# First-principles prediction of the information processing capacity of a simple genetic circuit

**DOI:** 10.1101/594325

**Authors:** Manuel Razo-Mejia, Sarah Marzen, Griffin Chure, Rachel Taubman, Muir Morrison, Rob Phillips

**Affiliations:** Division of Biology and Biological Engineering, California Institute of Technology, Pasadena, CA 91125, USA; Department of Physics, W. M. Keck Science Department; Department of Physics, California Institute of Technology, Pasadena, CA 91125, USA

## Abstract

Given the stochastic nature of gene expression, genetically identical cells exposed to the same environmental inputs will produce different outputs. This heterogeneity has been hypothesized to have consequences for how cells are able to survive in changing environments. Recent work has explored the use of information theory as a framework to understand the accuracy with which cells can ascertain the state of their surroundings. Yet the predictive power of these approaches is limited and has not been rigorously tested using precision measurements. To that end, we generate a minimal model for a simple genetic circuit in which all parameter values for the model come from independently published data sets. We then predict the information processing capacity of the genetic circuit for a suite of biophysical parameters such as protein copy number and protein-DNA affinity. We compare these parameter-free predictions with an experimental determination of protein expression distributions and the resulting information processing capacity of *E. coli* cells. We find that our minimal model captures the scaling of the cell-to-cell variability in the data and the inferred information processing capacity of our simple genetic circuit up to a systematic deviation.

---

As living organisms thrive in a given environment, they are faced with constant changes in their surroundings. From abiotic conditions such as temperature fluctuations or changes in osmotic pressure, to biological interactions such as cell-to-cell communication in a tissue or in a bacterial biofilm, living organisms of all types sense and respond to external signals. Fig. 1(A) shows a schematic of this process for a bacterial cell sensing a concentration of an extracellular chemical. At the molecular level where signal transduction unfolds mechanistically, there are physical constraints on the accuracy and precision of these responses given by intrinsic stochastic fluctuations [1]. This means that two genetically identical cells exposed to the same stimulus will not have identical responses [2].

**Figure 1.**
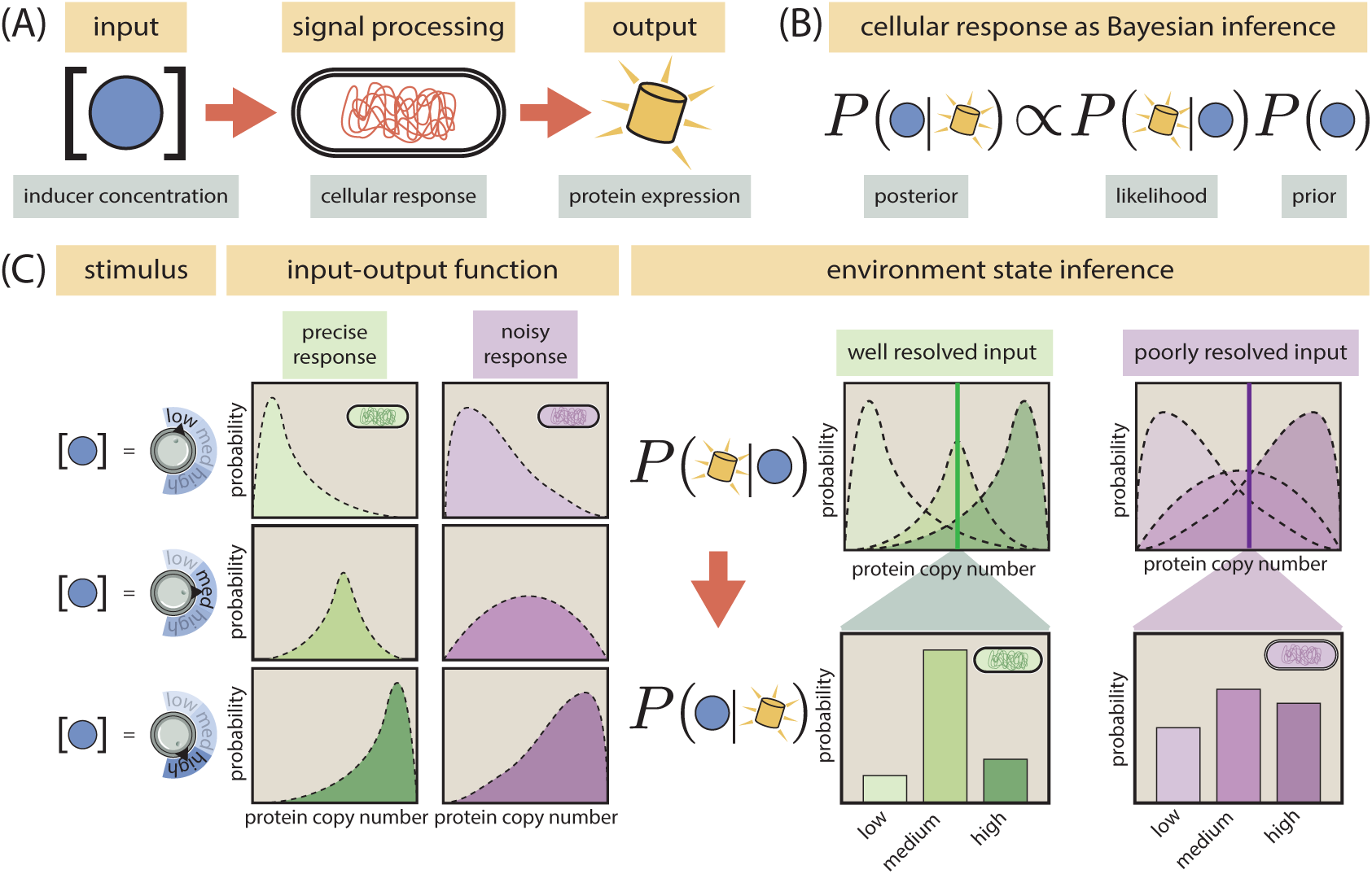
Cellular signaling systems sense the environment with different degrees of precision. (A) Schematic representation of a cell as a noisy communication channel. From an environmental input (inducer molecule concentration) to a phenotypic output (protein expression level), cellular signaling systems can be modeled as noisy communication channels. (B) We treat cellular response to an external stimulus as a Bayesian inference of the state of the environment. As the phenotype (protein level) serves as the internal representation of the environmental state (inducer concentration), the probability of a cell being in a specific environment given this internal representation *P* (*c*|*p*) is a function of the probability of the response given that environmental state *P* (*p*|*c*). (C) The precision of the inference of the environmental state depends on how well can cells resolve different inputs. For three different levels of input (left panel) the green strain responds more precisely than the purple strain since the output distributions overlap less (middle panel). This allows the green strain to make a more precise inference of the environmental state given a phenotypic response (right panel).

One implication of this noise in biological systems is that cells do not have an infinite resolution to distinguish signals and, as a consequence, there is a one-to-many mapping between inputs and outputs. Furthermore, given the limited number of possible outputs, there are overlapping responses between different inputs. This scenario can be map to a Bayesian inference problem where cells try to infer the state of the environment from their phenotypic response, as schematized in Fig. 1(B). The question then becomes this: how can one analyze this probabilistic, rather than deterministic, relationship between inputs and outputs? The abstract answer to this question was worked out in 1948 by Claude Shannon who, in his seminal work, founded the field of information theory [3]. Shannon developed a general framework for how to analyze information transmission through noisy communication channels. In his work, Shannon showed that the only quantity that satisfies three reasonable axioms for a measure of uncertainty was of the same functional form as the thermodynamic entropy – thereby christening his metric the information entropy [4]. He also gave a definition, based on this information entropy, for the relationship between inputs and outputs known as the mutual information. The mutual information *I* between input *c* and output *p*, given by

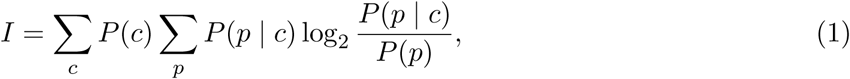

quantifies how much we learn about the state of the input *c* given that we get to observe the output *p*. In other words, the mutual information can be thought of as a generalized correlation coefficient that quantifies the degree to which the uncertainty about a random event decreases given the knowledge of the average outcome of another random event [5].

It is natural to conceive of scenarios in which living organisms that can better resolve signals might have an evolutionary benefit, making it more likely that their offspring will have a fitness advantage [6]. In recent years there has been a growing interest in understanding the theoretical limits on cellular information processing [7, 8], and in quantifying how close evolution has pushed cellular signaling pathways to these theoretical limits [9–11]. While these studies have treated the signaling pathway as a “black box,” explicitly ignoring all the molecular interactions taking place in them, other studies have explored the role that molecular players and regulatory architectures have on these information processing tasks [12–18]. Despite the great advances in our understanding of the information processing capabilities of molecular mechanisms, the field still lacks a rigorous experimental test of these detailed models with precision measurements on a simple system in which physical parameters can be perturbed. In this work we approach this task with a system that is both theoretically and experimentally tractable in which molecular parameters can be varied in a controlled manner.

Over the last decade the dialogue between theory and experiments in gene regulation has led to predictive power of models not only over the mean level of gene expression, but the noise as a function of relevant parameters such as regulatory protein copy numbers, affinity of these proteins to the DNA promoter, as well as the extracellular concentrations of inducer molecules [19–22]. These models based on equilibrium and non-equilibrium statistical physics have reached a predictive accuracy level such that, for simple cases, it is now possible to design input-output functions [23, 24]. This opens the opportunity to exploit these predictive models to tackle the question of how much information genetic circuits can process. This question lies at the heart of understanding the precision of the cellular response to environmental signals. Fig. 1(C) schematizes a scenario in which two bacterial strains respond with different levels of precision to three possible environmental states, i.e., inducer concentrations. The overlap between the three different responses is what precisely determines the resolution with which cells can distinguish different inputs. This is analogous to how the point spread function limits the ability to resolve two light emitting point sources.

In this work we follow the same philosophy of theory-experiment dialogue used to determine model parameters to predict from first principles the effect that biophysical parameters such as transcription factor copy number and protein-DNA affinity have on the information processing capacity of a simple genetic circuit. Specifically, to predict the mutual information between an extracellular chemical signal (input *c*) and the corresponding cellular response in the form of protein expression (output *p*), we must compute the input-output function *P* (*p* | *c*). To do so, we use a master-equation-based model to construct the protein copy number distribution as a function of an extracellular inducer concentration for different combinations of transcription factor copy numbers and binding sites. Having these input-output distributions allows us to compute the mutual information *I* between inputs and outputs for any arbitrary input distribution *P* (*c*). We opt to compute the channel capacity, i.e., the maximum information that can be processed by this gene regulatory architecture, defined as Eq. 1 maximized over all possible input distributions *P* (*c*). By doing so we examine the physical limits of what cells can do in terms of information processing by harboring these genetic circuits. Nevertheless, given the generality of the input-output function *P* (*p* | *c*) we derive, the model presented here can be used to compute the mutual information for any arbitrary input distribution *P* (*c*). All parameters used for our model were inferred from a series of studies that span several experimental techniques [20, 25–27], allowing us to make parameter-free predictions of this information processing capacity [28].

These predictions are then contrasted with experimental data, where the channel capacity is inferred from single-cell fluorescence distributions taken at different concentrations of inducer for cells with previously characterized biophysical parameters [20, 27]. We find that our parameter-free predictions quantitatively track the experimental data up to a systematic deviation. The lack of numerical agreement between our model and the experimental data poses new challenges towards having a foundational, first-principles understanding of the physics of cellular decision-making.

The reminder of the paper is organized as follows. In Section 1.1 we define the minimal theoretical model and parameter inference for a simple repression genetic circuit. Section 1.2 discusses how all parameters for the minimal model are determined from published datasets that explore different aspects of the simple repression motif. Section 1.3 computes the moments of the mRNA and protein distributions from this minimal model. In Section 1.4 we explore the consequences of variability in gene copy number during the cell cycle. In this section we compare experimental and theoretical quantities related to the moments of the distribution, specifically the predictions for the fold-change in gene expression (mean expression relative to an unregulated promoter) and the gene expression noise (standard deviation over mean). Section 1.5 follows with reconstruction of the full mRNA and protein distribution from the moments using the maximum entropy principle. Finally Section 1.6 uses the distributions from Section 1.5 to compute the maximum amount of information that the genetic circuit can process. Here we again contrast our zero-parameter fit predictions with experimental inferences of the channel capacity.

## 1 Results

### 1.1 Minimal model of transcriptional regulation

As a tractable circuit for which we have control over the parameters both theoretically and experimentally, we chose the so-called simple repression motif, a common regulatory scheme among prokaryotes [29]. This circuit consists of a single promoter with an RNA-polymerase (RNAP) binding site and a single binding site for a transcriptional repressor [20]. The regulation due to the repressor occurs via exclusion of the RNAP from its binding site when the repressor is bound, decreasing the likelihood of having a transcription event. As with many important macromolecules, we consider the repressor to be allosteric, meaning that it can exist in two conformations, one in which the repressor is able to bind to the specific binding site (active state) and one in which it cannot bind the specific binding site (inactive state). The environmental signaling occurs via passive import of an extracellular inducer that binds the repressor, shifting the equilibrium between the two conformations of the repressor [27]. In previous work we have extensively characterized the mean response of this circuit under different conditions using equilibrium based models [28]. Here we build upon these models to characterize the full distribution of gene expression with parameters such as repressor copy number and its affinity for the DNA being systematically varied.

As the copy number of molecular species is a discrete quantity, chemical master equations have emerged as a useful tool to model their inherent probability distribution [30]. In Fig. 2(A) we show the minimal model and the necessary set of parameters needed to compute the full distribution of mRNA and its protein gene product. Specifically, we assume a three-state model where the promoter can be found in a 1) transcriptionally active state (*A* state), 2) a transcriptionally inactive state without the repressor bound (*I* state) and 3) a transcriptionally inactive state with the repressor bound (*R* state). We do not assume that the transition between the active state *A* and the inactive state *I* occurs due to RNAP binding to the promoter as the transcription initiation kinetics involve several more steps than simple binding [31]. We coarse-grain all these steps into effective “on” and “off” states for the promoter, consistent with experiments demonstrating the bursty nature of gene expression in *E. coli* [19]. These three states generate a system of coupled differential equations for each of the three state distributions *P*_*A*_(*m, p*; *t*), *P*_*I*_(*m, p*; *t*) and *P*_*R*_(*m, p*; *t*), where *m* and *p* are the mRNA and protein count per cell, respectively and *t* is time. Given the rates depicted in Fig. 2(A) we define the system of ODEs for a specific *m* and *p*. For the transcriptionally active state, we have

**Figure 2.**
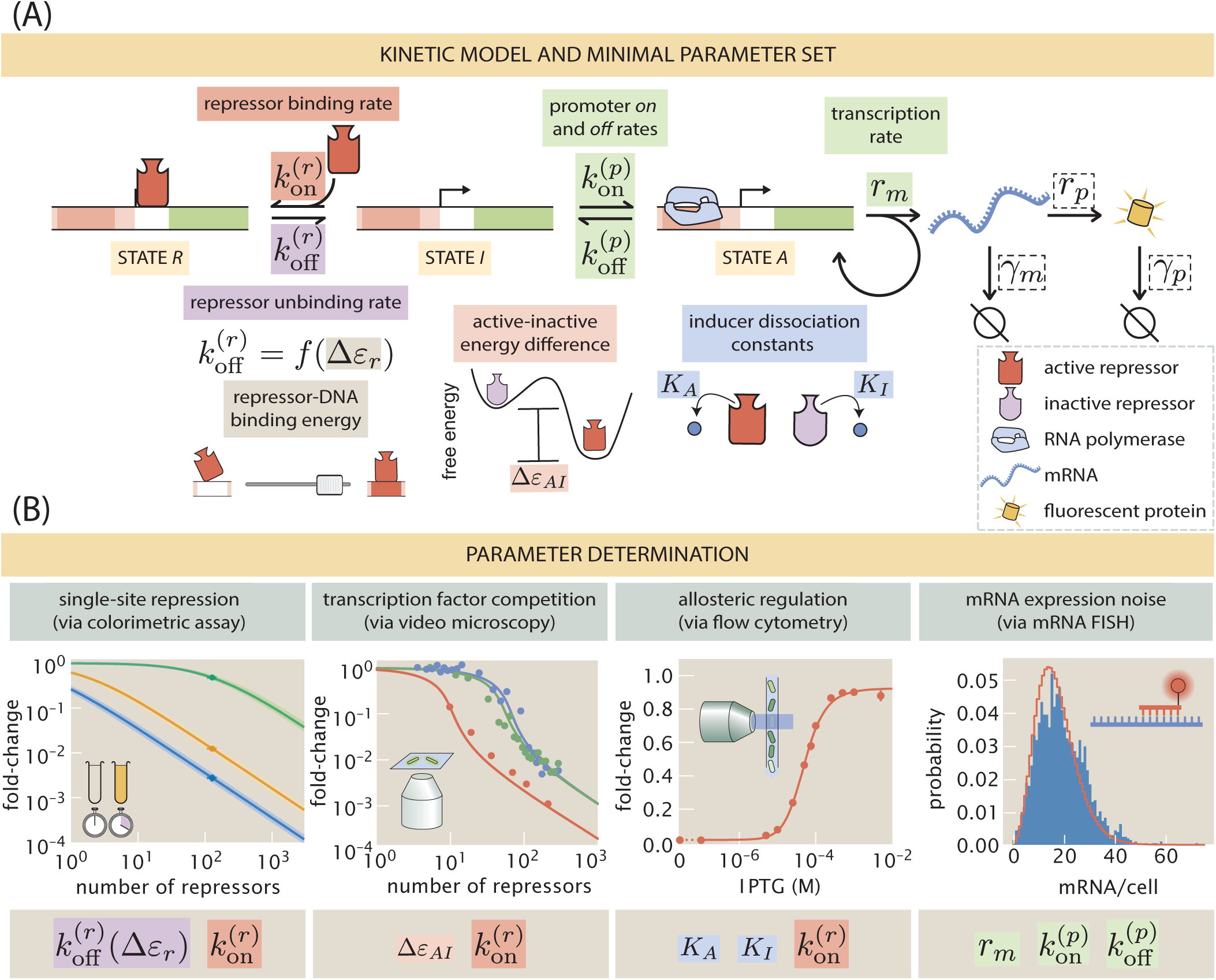
Minimal kinetic model of transcriptional regulation for a simple repression architecture. (A) Three-state promoter stochastic model of transcriptional regulation by a repressor. The regulation by the repressor occurs via exclusion of the transcription initiation machinery, not allowing the promoter to transition to the transcriptionally active state. All parameters highlighted with colored boxes were determined from published datasets based on the same genetic circuit. Parameters in dashed boxes were taken directly from values reported in the literature or adjusted to satisfy known biological restrictions. (B) Data sets used to infer the parameter values. From left to right Garcia & Phillips [20] is used to determine 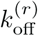 and 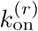, Brewster et al. [26] is used to determine Δ*ε*_*AI*_ and 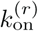, Razo-Mejia et al. [27] is used to determine *K*_*A*_, *K*_*I*_, and 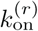 and Jones et al. [25] is used to determine 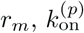, 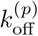.

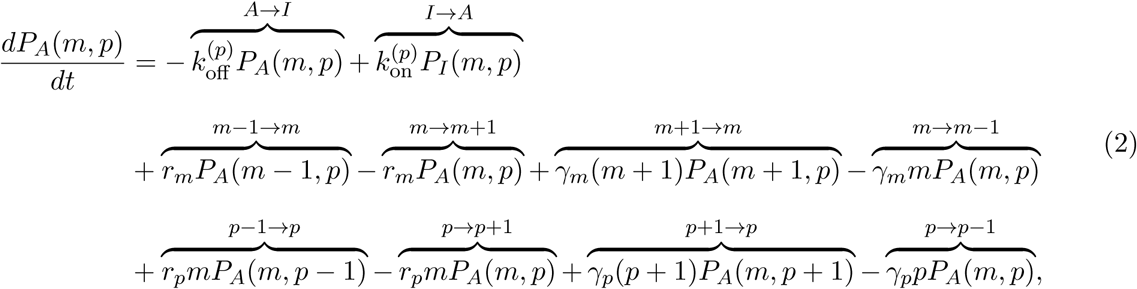

where the state transitions for each term are labeled by overbraces. For the transcriptionally inactive state *I*, we have

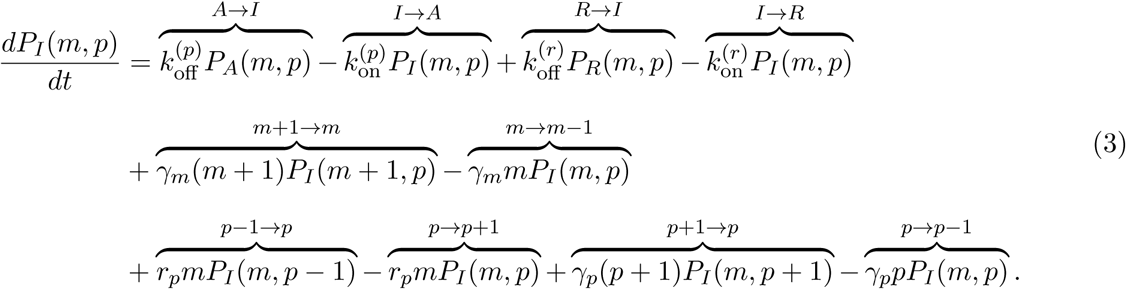

And finally, for the repressor bound state *R*,

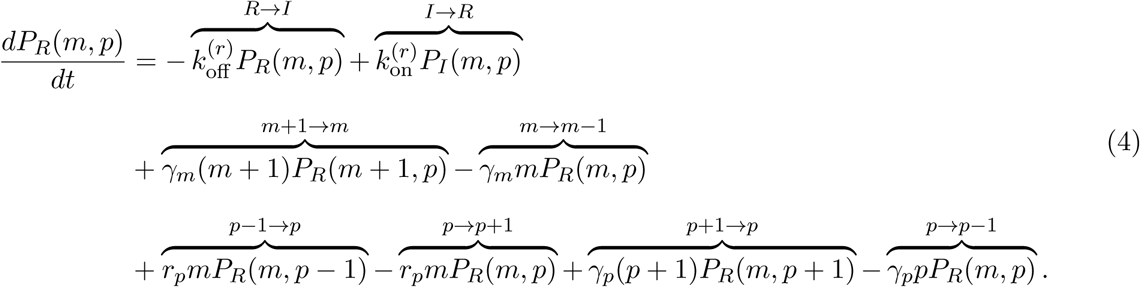

As we will discuss later in Section 1.4 the protein degradation term *γ*_*p*_ is set to zero since active protein degradation is slow compared to the cell cycle of exponentially growing bacteria, but rather we explicitly implement binomial partitioning of the proteins into daughter cells upon division [32].

It is convenient to rewrite these equations in a compact matrix notation [30]. For this we define the vector **P**(*m, p*) as

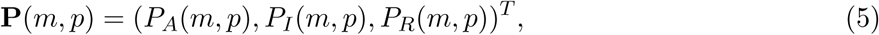

where ^*T*^ is the transpose. By defining the matrices **K** to contain the promoter state transitions, **R**_*m*_ and **Γ**_*m*_ to contain the mRNA production and degradation terms, respectively, and **R**_*p*_ and **Γ**_*p*_ to contain the protein production and degradation terms, respectively, the system of ODEs can then be written as (See Appendix S1 for full definition of these matrices)

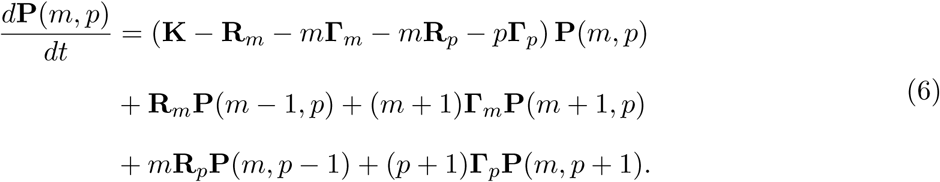

Having defined the gene expression dynamics we now proceed to determine all rate parameters in Eq. 6.

### 1.2 Inferring parameters from published data sets

A decade of research in our group has characterized the simple repression motif with an ever expanding array of predictions and corresponding experiments to uncover the physics of this genetic circuit [28]. In doing so we have come to understand the mean response of a single promoter in the presence of varying levels of repressor copy numbers and repressor-DNA affinities [20], due to the effect that competing binding sites and multiple promoter copies impose [26], and in recent work, assisted by the Monod-Wyman-Changeux (MWC) model, we expanded the scope to the allosteric nature of the repressor [27]. All of these studies have exploited the simplicity and predictive power of equilibrium approximations to these non-equilibrium systems [33]. We have also used a similar kinetic model to that depicted in Fig. 2(A) to study the noise in mRNA copy number [25]. As a test case of the depth of our theoretical understanding of this simple transcriptional regulation system we combine all of the studies mentioned above to inform the parameter values of the model presented in Fig. 2(A). Fig. 2(B) schematizes the data sets and experimental techniques used to measure gene expression along with the parameters that can be inferred from them.

Appendix S2 expands on the details of how the inference was performed for each of the parameters. Briefly, the promoter activation and inactivation rates 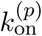 and 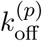, as well as the transcription rate *r*_*m*_ were obtained in units of the mRNA degradation rate *γ*_*m*_ by fitting a two-state promoter model (no state *R* from Fig. 2(A)) [34] to mRNA FISH data of an unregulated promoter (no repressor present in the cell) [25]. The repressor on rate is assumed to be of the form 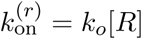 where *k*_*o*_ is a diffusion-limited on rate and [*R*] is the concentration of active repressor in the cell [25]. This concentration of active repressor is at the same time determined by the repressor copy number in the cell, and the fraction of these repressors that are in the active state, i.e. able to bind DNA. Existing estimates of the transition rates between conformations of allosteric molecules set them at the microsecond scale [35]. By considering this to be representative for our repressor of interest, the separation of time-scales between the rapid conformational changes of the repressor and the slower downstream processes such as the open-complex formation processes allow us to model the probability of the repressor being in the active state as an equilibrium MWC process. The parameters of the MWC model *K*_*A*_, *K*_*I*_ and Δ*ε*_*AI*_ were previously characterized from video-microscopy and flow-cytometry data [27]. For the repressor off rate, 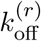, we take advantage of the fact that the mean mRNA copy number as derived from the model in Fig. 2(A) cast in the language of rates is of the same functional form as the equilibrium model cast in the language of binding energies [36]. Therefore the value of the repressor-DNA binding energy Δ*ε*_*r*_ constrains the value of the repressor off rate 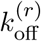. These constraints on the rates allow us to make self-consistent predictions under both the equilibrium and the kinetic framework. Having all parameters in hand, we can now proceed to solve he gene expression dynamics.

### 1.3 Computing the moments of the mRNA and protein distributions

Finding analytical solutions to chemical master equations is often fraught with difficulty. An alternative approach is to to approximate the distribution. One such scheme of approximation, the maximum entropy principle, makes use of the moments of the distribution to approximate the full distribution. In this section we will demonstrate an iterative algorithm to compute the mRNA and protein distribution moments.

The kinetic model for the simple repression motif depicted in Fig. 2(A) consists of an infinite system of ODEs for each possible pair of mRNA and protein copy number, (*m, p*). To compute any moment of the distribution, we define a vector

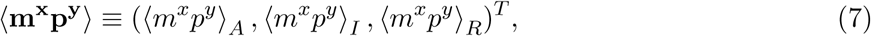

where ⟨*m*^*x*^*p*^*y*^⟩_*S*_ is the expected value of *m*^*x*^*p*^*y*^ in state *S* ∈ {*A, I, R*} for *x, y* ∈ ℕ. In other words, just as we defined the vector **P**(*m, p*), here we define a vector to collect the expected value of each of the promoter states. By definition, any of these moments ⟨*m*^*x*^*p*^*y*^⟩_*S*_ can be computed as

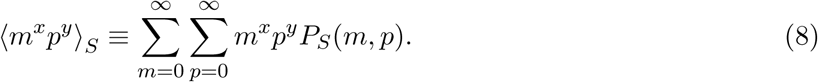

Summing over all possible values for *m* and *p* in Eq. 6 results in an ODE for any moment of the distribution of the form (See Appendix S3 for full derivation)

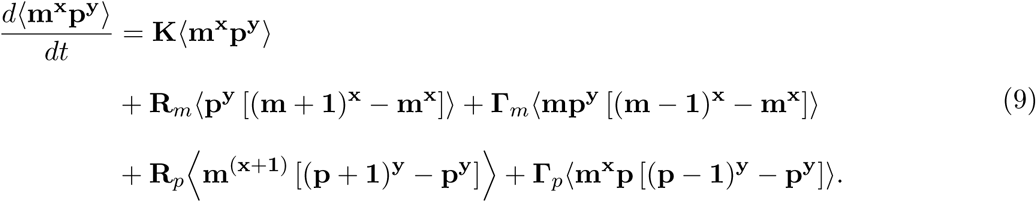

Given that all transitions in our stochastic model are first order reactions, Eq. 9 has no moment-closure problem [14]. This means that the dynamical equation for a given moment only depends on lower moments (See Appendix S3 for full proof). This feature of our model implies, for example, that the second moment of the protein distribution ⟨*p*^2^⟩ depends only on the first two moments of the mRNA distribution ⟨*m*⟩ and ⟨ *m*^*2*^⟩, the first protein moment ⟨ *p*⟩, and the cross-correlation term ⟨ *mp*⟩. We can therefore define ***µ***^(**x**,**y**)^ to be a vector containing all moments up to ⟨**m**^**x**^**p**^**y**^⟩ for all promoter states,

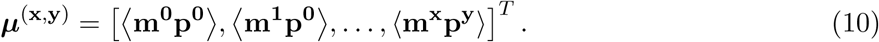

Explicitly for the three-state promoter model depicted in Fig. 2(A) this vector takes the form

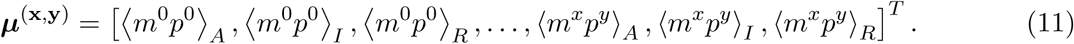

Given this definition we can compute the general moment dynamics as

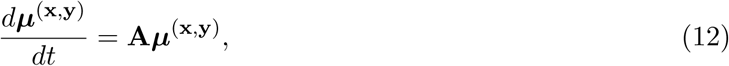

where **A** is a square matrix that contains all the numerical coefficients that relate each of the moments. We can then use Eq. 9 to build matrix **A** by iteratively substituting values for the exponents *x* and *y* up to a specified value. In the next section, we will use Eq. 12 to numerically integrate the dynamical equations for our moments of interest as cells progress through the cell cycle. We will then use the value of the moments of the distribution to approximate the full gene expression distribution. This method is computationally more efficient than trying to numerically integrate the infinite set of equations describing the full probability distribution **P**(*m, p*), or using a stochastic algorithm to sample from the distribution.

### 1.4 Accounting for cell-cycle dependent variability in gene dosage

As cells progress through the cell cycle, the genome has to be replicated to guarantee that each daughter cell receives a copy of the genetic material. As replication of the genome can take longer than the total cell cycle, this implies that cells spend part of the cell cycle with multiple copies of each gene depending on the cellular growth rate and the relative position of the gene with respect to the replication origin [37]. Genes closer to the replication origin spend a larger fraction of the cell cycle with multiple copies compared to genes closer to the replication termination site [37]. Fig. 3(A) depicts a schematic of this process where the replication origin (*oriC*) and the relevant locus for our experimental measurements (*galK*) are highlighted.

**Figure 3.**
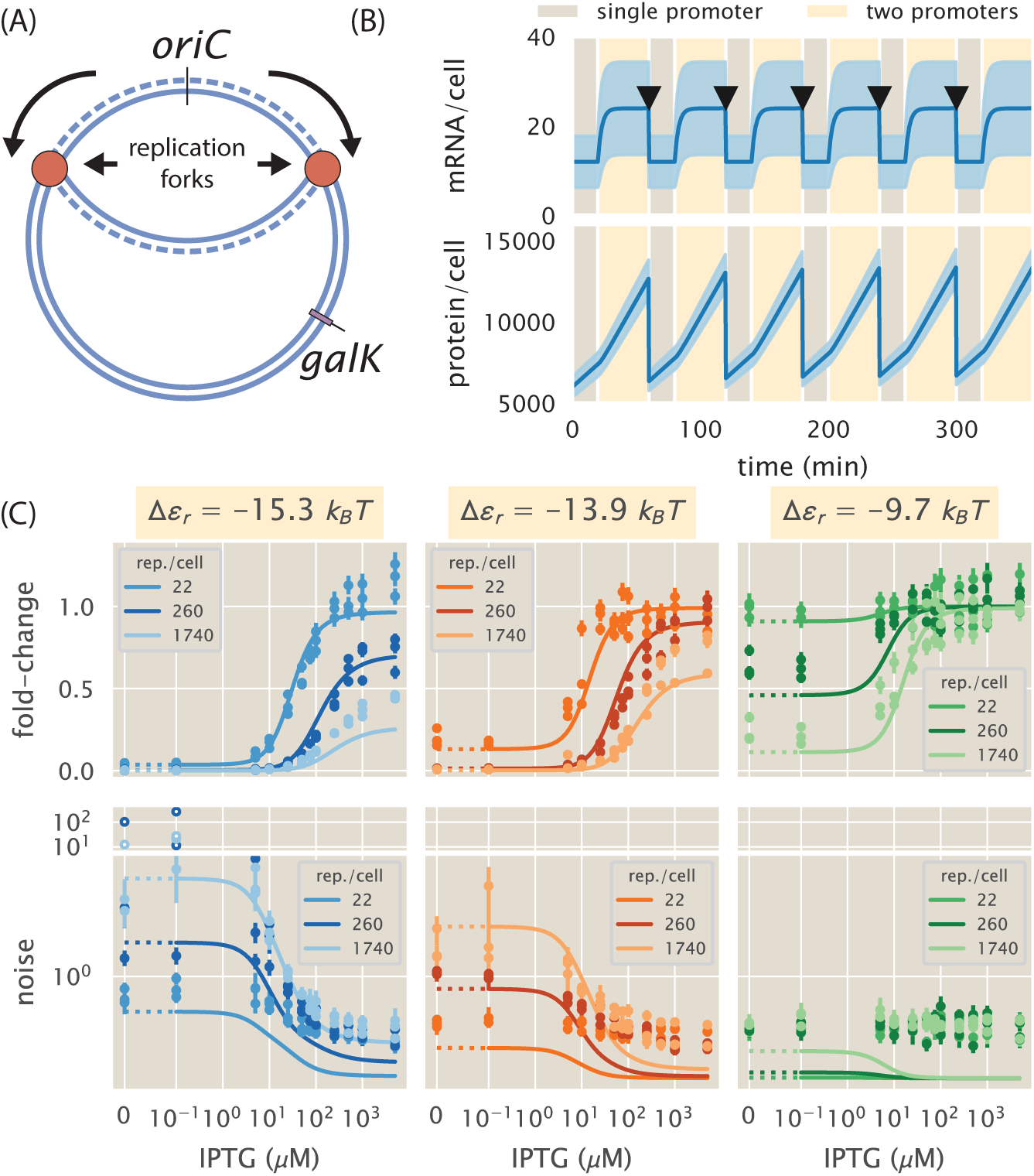
Accounting for gene copy number variability during the cell cycle. (A) Schematic of a replicating bacterial genome. As cells progress through the cell cycle the genome is replicated, duplicating gene copies for a fraction of the cell cycle before the cell divides. *oriC* indicates the replication origin, and *galK* indicates the locus at which the YFP reporter construct was integrated. (B) mean (solid line) *±* standard deviation (shaded region) for the mRNA (upper panel) and protein (lower panel) dynamics. Cells spend a fraction of the cell cycle with a single copy of the promoter (light brown) and the rest of the cell cycle with two copies (light yellow). Black arrows indicate time of cell division. (C) Zero parameter-fit predictions (lines) and experimental data (circles) of the gene expression fold-change (upper row) and noise (lower row) for repressor binding sites with different affinities (different columns) and different repressor copy numbers per cell (different lines on each panel). Error bars in data represent the 95% confidence interval on the quantities as computed from 10,000 bootstrap estimates generated from *>* 500 single-cell fluorescence measurements. In the theory curves, dotted lines indicate plot in linear scale to include zero, while solid lines indicate logarithmic scale. For visual clarity, data points in the noise panel with exceptionally large values coming from highly repressed strains are plotted on a separate panel.

Since this change in gene copy number has been shown to have an effect on cell-to-cell variability in gene expression [25, 38], we now extend our minimal model to account for these changes in gene copy number during the cell cycle. We reason that the only difference between the single-copy state and the two-copy state of the promoter is a doubling of the mRNA production rate *r*_*m*_. In particular, the promoter activation and inactivation rates 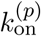 and 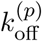 and the mRNA production rate *r*_*m*_ inferred in Section 1.1 assume that cells spend a fraction *f* of the cell cycle with one copy of the promoter (mRNA production rate *r*_*m*_) and a fraction (1 − *f*) of the cell cycle with two copies of the promoter (mRNA production rate 2*r*_*m*_). This inference was performed considering that at each cell state the mRNA level immediately reaches the steady state value for the corresponding mRNA production rate. This assumption is justified since the timescale to reach this steady state depends only on the degradation rate *γ*_*m*_, which for the mRNA is much shorter (≈ 3 min) than the length of the cell cycle (≈ 60 min for our experimental conditions) [39]. Appendix S2 shows that a model accounting for this gene copy number variability is able to capture data from single molecule mRNA counts of an unregulated (constitutively expressed) promoter.

Given that the protein degradation rate *γ*_*p*_ in our model is set by the cell division time, we do not expect that the protein count will reach the corresponding steady state value for each stage in the cell cycle. In other words, cells do not spend long enough with two copies of the promoter for the protein level to reach the steady state value corresponding to a transcription rate of 2*r*_*m*_. We therefore use the dynamical equations developed in Section 1.3 to numerically integrate the time trajectory of the moments of the distribution with the corresponding parameters for each phase of the cell cycle. Fig. 3(B) shows an example corresponding to the mean mRNA level (upper panel) and the mean protein level (lower panel) for the case of the unregulated promoter. Given that we inferred the promoter rate parameters considering that mRNA reaches steady state in each stage, we see that the numerical integration of the equations is consistent with the assumption of having the mRNA reach a stable value in each stage (See Fig. 3(B) upper panel). On the other hand, the mean protein level does not reach a steady state at either of the cellular stages. Nevertheless it is notable that after several cell cycles the trajectory from cycle to cycle follows a repetitive pattern (See Fig. 3(B) lower panel). Previously we have experimentally observed this repetitive pattern by tracking the expression level over time with video microscopy as observed in Fig. 18 of [28].

To test the effects of including this gene copy number variability in our model we now compare the predictions of the model with experimental data. As detailed in the Methods section, we obtained single-cell fluorescence values of different *E. coli* strains carrying a YFP gene under the control of the LacI repressor. Each strain was exposed to twelve different inducer concentrations for ≈ 8 generations for cells to adapt to the media. The strains imaged spanned three orders of magnitude in repressor copy number and three distinct repressor-DNA affinities. Since growth was asynchronous, we reason that cells were randomly sampled at all stages of the cell cycle. Therefore, when computing statistics from the data such as the mean fluorescence value, in reality we are averaging over the cell cycle. In other words, as depicted in Fig. 3(B), quantities such as the mean protein copy number change over time, i.e. ⟨*p*⟩≡ ⟨*p*(*t*)⟩. This means that computing the mean of a population of unsynchronized cells is equivalent to averaging this time-dependent mean protein copy number over the span of the cell cycle. Mathematically this is expressed as

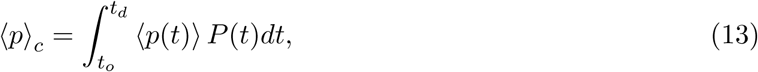

where ⟨*p*(*t*)⟩ represents the first moment of the protein distribution as computed from Eq. 9, ⟨*p*⟩_*c*_ represents the average protein copy number over a cell cycle, *t*_*o*_ represents the start of the cell cycle,*t*_*d*_ represents the time of cell division, and *P* (*t*) represents the probability of any cell being at time *t* ∈ [*t*_*o*_, *t*_*d*_] of their cell cycle. We do not consider cells uniformly distributed along the cell cycle since it is known that cells age is exponentially distributed, having more younger than older cells at any point in time [40] (See Appendix S9 for further details). All computations hereafter are therefore done by applying an average like that in Eq. 13 for the span of a cell cycle. We remind the reader that these time averages are done under a fixed environmental state. It is the trajectory of cells over cell cycles under a constant environment that we need to account for. It is through this averaging over the span of a cell cycle that we turn a periodic process as the one shown in Fig. 3(B) into a stationary process that we can compare with experimental data and, as we will see later, use to reconstruct the steady state gene expression distribution.

Fig. 3(C) compares zero-parameter fit predictions (lines) with experimentally determined quantities (points). The upper row shows the non-dimensional quantity known as the fold-change in gene expression [20]. This fold-change is defined as the relative mean gene expression level with respect to an unregulated promoter. For protein this is

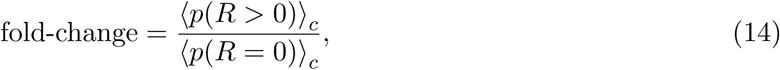

where ⟨*p*(*R>* 0)⟩_*c*_ represents the mean protein count for cells with non-zero repressor copy number count *R* over the entire cell cycle, and ⟨*p*(*R* = 0)⟩_*c*_ represents the equivalent for a strain with no repressors present. The experimental points were determined from the YFP fluorescent intensities of cells with varying repressor copy number and a Δ*lacI* strain with no repressor gene present (See Methods for further details). The fold-change in gene expression has previously served as a metric to test the validity of equilibrium-based models [36]. We note that the curves shown in the upper panel of Fig. 3(C) are consistent with the predictions from equilibrium models [27] despite being generated from a clearly non-equilibrium process as shown in Fig. 3(B). The kinetic model from Fig. 2(A) goes beyond the equilibrium picture to generate predictions for moments of the distribution other than the mean mRNA or mean protein count. To test this extended predictive power the lower row of Fig. 3(C) shows the noise in gene expression defined as the standard deviation over the mean protein count, accounting for the changes in gene dosage during the cell cycle. Although our model systematically underestimates the noise in gene expression, the zero-parameter fits capture the scaling of this noise. Possible origins of this systematic discrepancy could be the intrinsic cell-to-cell variability of rate parameters given the variability in the molecular components of the central dogma machinery [25], or noise generated by irreversible non-equilibrium reactions not explicitly taken into account in our minimal model [41]. The large errors for the highly repressed strains (lower left panel in Fig. 3(C)) are a result of having a small number in the denominator - mean fluorescence level - when computing the noise. Although the model is still highly informative about the physical nature of how cells regulate their gene expression, the lack of exact numerical agreement between theory and data opens an opportunity to gain new insights into the biophysical origin of cell-to-cell variability. In Appendix S8 we explore empirical ways to account for this systematic deviation. We direct the reader to Appendix S4 where equivalent predictions are done ignoring the changes in gene dosage due to the replication of the genome.

### 1.5 Maximum Entropy approximation

Having numerically computed the moments of the mRNA and protein distributions as cells progress through the cell cycle, we now proceed to make an approximate reconstruction of the full distributions given this limited information. As hinted in Section 1.3 the maximum entropy principle, first proposed by E.T. Jaynes in 1957 [42], approximates the entire distribution by maximizing the Shannon entropy subject to constraints given by the values of the moments of the distribution [42]. This procedure leads to a probability distribution of the form (See Appendix S5 for full derivation)

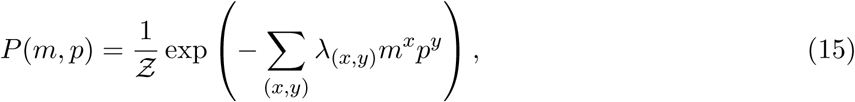

where *λ*_(*x,y*)_ is the Lagrange multiplier associated with the constraint set by the moment ⟨*m*^*x*^*p*^*y*^⟩, and 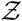 is a normalization constant. The more moments ⟨*m*^*x*^*p*^*y*^⟩ included as constraints, the more accurate the approximation resulting from Eq. 15 becomes.

The computational challenge then becomes an optimization routine in which the values for the Lagrange multipliers *λ*_(*x,y*)_ that are consistent with the constraints set by the moment values ⟨*m*^*x*^*p*^*y*^⟩ need to be found. This is computationally more efficient than sampling directly out of the master equation with a stochastic algorithm (see Appendix S6 for further comparison between maximum entropy estimates and the Gillespie algorithm). Appendix S5 details our implementation of a robust algorithm to find the values of the Lagrange multipliers. Fig. 4(A) shows example predicted protein distributions reconstructed using the first six moments of the protein distribution for a suite of different biophysical parameters and environmental inducer concentrations. As repressor-DNA binding affinity (columns in Fig. 4(A)) and repressor copy number (rows in Fig. 4(A)) are varied, the responses to different signals, i.e. inducer concentrations, overlap to varying degrees. For example, the upper right corner frame with a weak binding site (Δ*ε*_*r*_ = −9.7 *k*_*B*_*T*) and a low repressor copy number (22 repressors per cell) have virtually identical distributions regardless of the input inducer concentration. This means that cells with this set of parameters cannot resolve any difference in the concentration of the signal. As the number of repressors is increased, the degree of overlap between distributions decreases, allowing cells to better resolve the value of the signal input. On the opposite extreme the lower left panel shows a strong binding site (Δ*ε*_*r*_ = −15.3 *k*_*B*_*T*) and a high repressor copy number (1740 repressors per cell). This parameter combination shows overlap between distributions since the high degree of repression centers all distributions towards lower copy numbers, again giving little ability for the cells to resolve the inputs. In Fig. 4(B) and Appendix S5 we show the comparison of these predicted cumulative distributions with the experimental single-cell fluorescence distributions. Given the systematic deviation of our predictions for the protein copy number noise highlighted in Fig. 3(C), the theoretical distributions (dashed lines) underestimate the width of the experimental data. We again direct the reader to Appendix S8 for an exploration of empirical changes to the moments that improve the agreement of the predictions. In the following section we formalize the notion of how well cells can resolve different inputs from an information theoretic perspective via the channel capacity.

**Figure 4.**
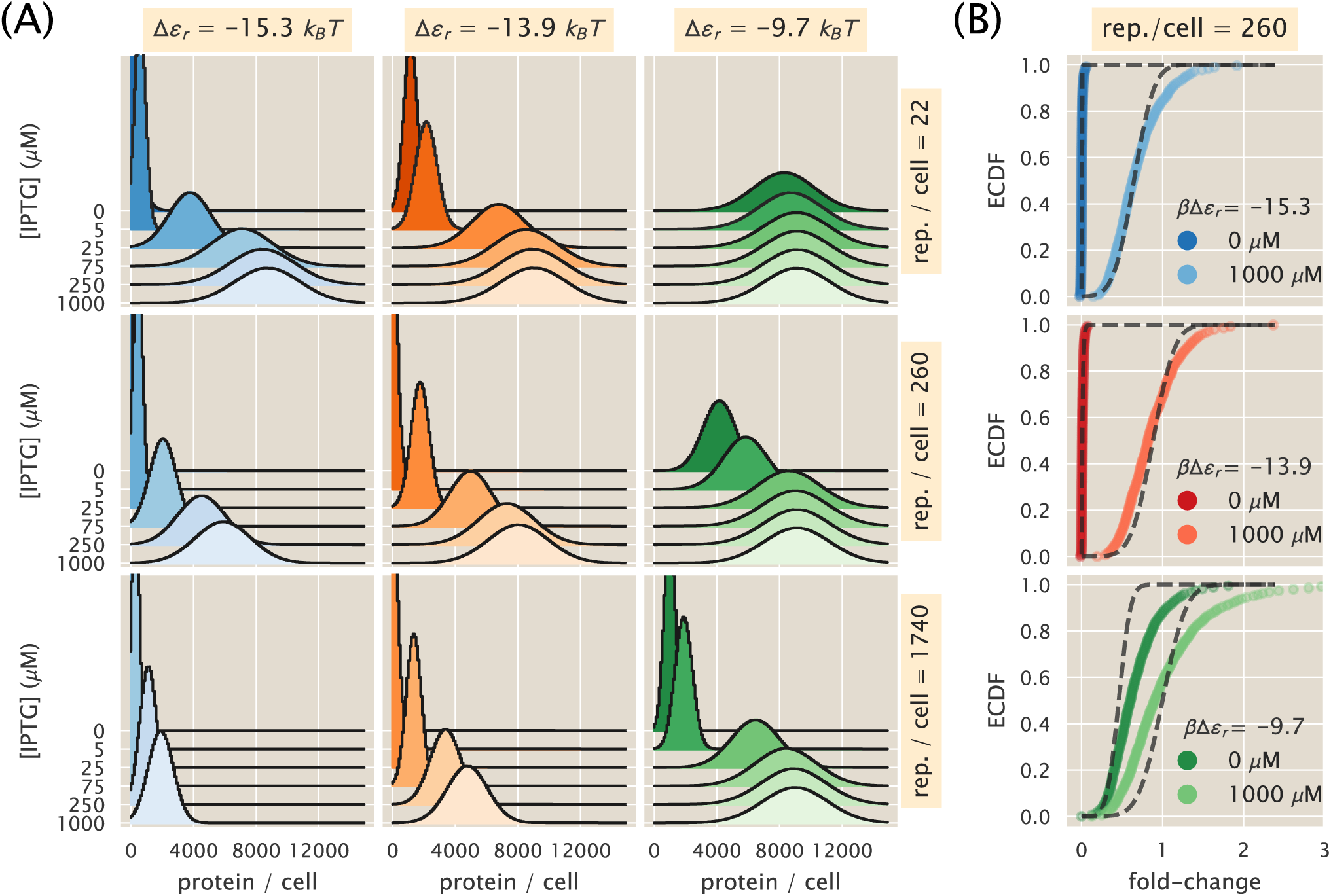
Maximum entropy protein distributions for varying physical parameters. (A) Predicted protein distributions under different inducer (IPTG) concentrations for different combinations of repressor-DNA affinities (columns) and repressor copy numbers (rows). The first six moments of the protein distribution used to constrain the maximum entropy approximation were computed by integrating Eq. 9 as cells progressed through the cell cycle as described in Section 1.4. (B) Theory-experiment comparison of predicted fold-change empirical cumulative distribution functions (ECDF). Each panel shows two example concentrations of inducer (colored curves) with their corresponding theoretical predictions (dashed lines). Distributions were normalized to the mean expression value of the unregulated strain in order to compare theoretical predictions in discrete protein counts with experimental fluorescent measurements in arbitrary units.

### 1.6 Theoretical prediction of the channel capacity

We now turn our focus to the channel capacity, which is a metric by which we can quantify the degree to which cells can measure the environmental state (in this context, the inducer concentration).

The channel capacity is defined as the mutual information *I* between input and output (Eq. 1), maximized over all possible input distributions *P* (*c*). If used as a metric of how reliably a signaling system can infer the state of the external signal, the channel capacity, when measured in bits, is commonly interpreted as the logarithm of the number of states that the signaling system can properly resolve. For example, a signaling system with a channel capacity of *C* bits is interpreted as being able to resolve 2^*C*^ states, though channel capacities with fractional values are allowed. We therefore prefer the Bayesian interpretation that the mutual information quantifies the improvement in the inference of the input when considering the output compared to just using the prior distribution of the input by itself for prediction [14, 43]. Under this interpretation a channel capacity of a fractional bit still quantifies an improvement in the ability of the signaling system to infer the value of the extracellular signal compared to having no sensing system at all.

Computing the channel capacity implies optimizing over an infinite space of possible distributions *P* (*c*). For special cases in which the noise is small compared to the dynamic range, approximate analytical equations have been derived [17]. But given the high cell-to-cell variability that our model predicts, the conditions of the so-called small noise approximation are not satisfied. We therefore appeal to a numerical solution known as the Blahut-Arimoto algorithm [44] (See Appendix S7 for further details). Fig. 5(A) shows zero-parameter fit predictions of the channel capacity as a function of the number of repressors for different repressor-DNA affinities (solid lines). These predictions are contrasted with experimental determinations of the channel capacity as inferred from single-cell fluorescence intensity distributions taken over 12 different concentrations of inducer. Briefly, from single-cell fluorescence measurements we can approximate the input-output distribution *P* (*p* | *c*). Once these conditional distributions are fixed, the task of finding the input distribution at channel capacity becomes a computational optimization routine that can be undertaken using conjugate gradient or similar algorithms. For the particular case of the channel capacity on a system with a discrete number of inputs and outputs the Blahut-Arimoto algorithm is built in such a way that it guarantees the convergence towards the optimal input distribution (See Appendix S7 for further details). Fig. 5(B) shows example input-output functions for different values of the channel capacity. This illustrates that having access to no information (zero channel capacity) is a consequence of having overlapping input-output functions (lower panel). On the other hand, the more separated the input-output distributions are (upper panel) the higher the channel capacity can be.

**Figure 5.**
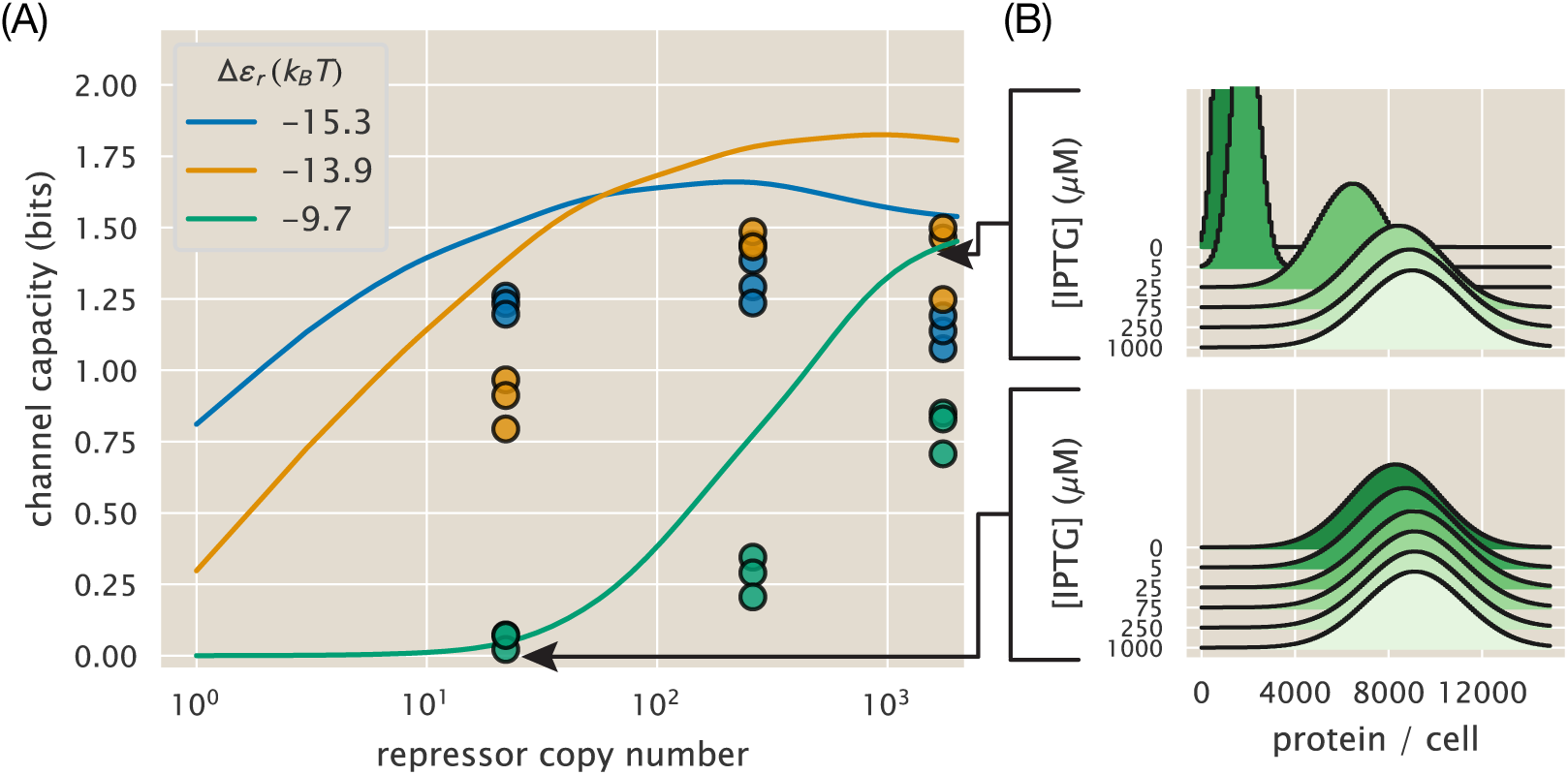
Comparison of theoretical and experimental channel capacity. (A) Channel capacity as inferred using the Blahut-Arimoto algorithm [44] for varying number of repressors and repressor-DNA affinities. All inferences were performed using 12 IPTG concentrations as detailed in the Methods. Curves represent zero-parameter fit predictions made with the maximum entropy distributions as shown in Fig. 4. Points represent inferences made from single cell fluorescence distributions (See Appendix S7 for further details). Theoretical curves were smoothed using a Gaussian kernel to remove numerical precision errors. (B) Example input-output functions in opposite limits of channel capacity. Lower panel illustrates that zero channel capacity indicates that all distributions overlap. Upper panel illustrates that as the channel capacity increases, the separation between distributions increases as well. Arrows point to the corresponding channel capacity computed from the predicted distributions.

All theoretical predictions in Fig. 5(A) are systematically above the experimental data. Although our theoretical predictions in Fig. 5(A) do not numerically match the experimental inference of the channel capacity, the model does capture interesting qualitative features of the data that are worth highlighting. On one extreme, for cells with no transcription factors, there is no information processing potential as this simple genetic circuit would be constitutively expressed regardless of the environmental state. As cells increase the transcription factor copy number, the channel capacity increases until it reaches a maximum before falling back down at high repressor copy number since the promoter would be permanently repressed. The steepness of the increment in channel capacity as well as the height of the maximum expression is highly dependent on the repressor-DNA affinity. For strong binding sites (blue curve in Fig. 5(A)) there is a rapid increment in the channel capacity, but the maximum value reached is smaller compared to a weaker binding site (orange curve in Fig. 5(A)). In Appendix S8 we show using the small noise approximation [9, 17] that if the systematic deviation of our predictions on the cell-to-cell variability was explained with a multiplicative constant, i.e. all noise predictions can be corrected by multiplying them by a single constant, we would expect the channel capacity to be off by a constant additive factor. This factor of ≈ 0.43 bits can recover the agreement between the model and the experimental data.

## Discussion

Building on Shannon’s formulation of information theory, there have been significant efforts using this theoretical framework to understand the information processing capabilities of biological systems, and the evolutionary consequences for organisms harboring signal transduction systems [1, 6, 9, 45–47]. Recently, with the mechanistic dissection of molecular signaling pathways, significant progress has been made on the question of the physical limits of cellular detection and the role that features such as feedback loops play in this task [7, 14, 16, 48, 49]. But the field still lacks a rigorous experimental test of these ideas with precision measurements on a system that is tractable both experimentally and theoretically.

In this paper we take advantage of the recent progress on the quantitative modeling of input-output functions of genetic circuits to build a minimal model of the simple repression motif [28]. By combining a series of studies on this circuit spanning diverse experimental methods for measuring gene expression under a myriad of different conditions, we possess complete a priori parametric knowledge – allowing us to generate parameter-free predictions for processes related to information processing. Some of the model parameters for our kinetic formulation of the input-output function are informed by inferences made from equilibrium models. We use the fact that if both kinetic and thermodynamic languages describe the same system, the predictions must be self-consistent. In other words, if the equilibrium model can only make statements about the mean mRNA and mean protein copy number because of the way these models are constructed, those predictions must be equivalent to what the kinetic model has to say about these same quantities. This condition therefore constrains the values that the kinetic rates in the model can take. To test whether or not the equilibrium picture can reproduce the predictions made by the kinetic model we compare the experimental and theoretical fold-change in protein copy number for a suite of biophysical parameters and environmental conditions (Fig. 3(C) upper row). The agreement between theory and experiment demonstrates that these two frameworks can indeed make consistent predictions.

The kinetic treatment of the system brings with it increasing predictive power compared to the equilibrium picture. Under the kinetic formulation, the predictions are not limited only to the mean but to any of the moments of the mRNA and protein distributions. We first test these novel predictions by comparing the noise in protein copy number (standard deviation / mean) with experimental data. Our minimal model predicts the noise up to a systematic deviation. The physical or biological origins of this discrepancy remain an open question. In that way the work presented here exposes the status quo of our understanding of gene regulation in bacteria, posing new questions to be answered with future refinements of the model. We then extend our analysis to infer entire protein distributions at different input signal concentrations by using the maximum entropy principle. What this means is that we compute moments of the protein distribution, and then use these moments to build an approximation to the full distribution. These predicted distributions are then compared with experimental single-cell distributions as shown in Fig. 4(B) and Appendix S5. Again, here although our minimal model systematically underestimates the width of the distributions, it informs how changes in parameters such as protein copy number or protein-DNA binding affinity will affect the full probabilistic input-output function of the genetic circuit, up to a multiplicative constant. We then use our model to predict the information processing capacity.

By maximizing the mutual information between input signal concentration and output protein distribution over all possible input distributions, we predict the channel capacity of the system over a suite of biophysical parameters such as varying repressor protein copy number and repressor-DNA binding affinity. Although there is no reason to assume the the simplified synthetic circuit we used as an experimental model operates optimally given the distribution of inputs, the relevance of the channel capacity comes from its interpretation as a metric of the physical limit of how precise an inference cells can make about what the state of the environment is. Our model, despite the systematic deviations, makes non-trivial predictions such as the existence of an optimal repressor copy number for a given repressor-DNA binding energy, predicting the channel capacity up to an additive constant (See Fig. 5). The origin of this optimal combination of repressor copy number and binding energy differs from previous publications in which an extra term associated with the cost of producing protein was included in the model [16]. This optimal parameter combination is a direct consequence of the fact that the LacI repressor cannot be fully deactivated [27]. This implies that as the number of repressors increases, a significant number of them are still able to bind to the promoter even at saturating concentrations of inducer. This causes all of the input-output functions to be shift towards low expression levels, regardless of the inducer concentration, decreasing the amount of information that the circuit is able to process.

We consider it important to highlight the limitations of the work presented here. The previously discussed systematic deviation for the noise and skewness of the predicted distributions (See Appendix S4), and therefore of the predicted distributions and channel capacity, remains an unresolved question that deserves to be addressed in further iterations of our minimal model. Also, as first reported in [27], our model fails to capture the steepness of the fold-change induction curve for the weakest repressor binding site (See Fig. 3(B)). Furthermore the minimal model in Fig. 2(A), despite being widely used, is an oversimplification of the physical picture of how the transcriptional machinery works. The coarse-graining of all the kinetic steps involved in transcription initiation into two effective promoter states – active and inactive – ignores potential kinetic regulatory mechanisms of intermediate states [50]. Moreover it has been argued that despite the fact that the mRNA count distribution does not follow a Poisson distribution, this effect could be caused by unknown factors not at the level of transcriptional regulation [51].

The findings of this work open the opportunity to accurately test intriguing ideas that connect Shannon’s metric of how accurately a signaling system can infer the state of the environment, with Darwinian fitness [6]. Beautiful work along these lines has been done in the context of the developmental program of the early *Drosophila* embryo [9, 11]. These studies demonstrated that the input-output function of the pair-rule genes works at channel capacity, suggesting that selection has acted on these signaling pathways, pushing them to operate at the limit of what the physics of these systems allows. Our system differs from the early embryo in the sense that we have a tunable circuit with variable amounts of information processing capabilities. Furthermore, compared with the fly embryo in which the organism tunes both the input and output distributions over evolutionary time, we have experimental control of the distribution of inputs that the cells are exposed to. Consequently this means that instead of seeing the final result of the evolutionary process, we would be able to set different environmental challenges, and track over time the evolution of the population. These experiments could shed light into the suggestive hypothesis of information bits as a trait on which natural selection acts. We see this exciting direction as part of the overall effort in quantitative biology of predicting evolution [52].

## 2 Materials and Methods

### 2.1 *E. coli* strains

All strains used in this study were originally made for [27]. We chose a subset of three repressor copy numbers that span two orders of magnitude. We refer the reader to [27] for details on the construction of these strains. Briefly, the strains have a construct consisting of the *lacUV5* promoter and one of three possible binding sites for the *lac* repressor (O1, O2, and O3) controlling the expression of a YFP reporter gene. This construct is integrated into the genome at the *galK* locus. The number of repressors per cell is varied by changing the ribosomal binding site controlling the translation of the *lac* repressor gene. The repressor constructs were integrated in the *ybcN* locus. Finally, all strains used in this work constitutively express an mCherry reporter from a low copy number plasmid. This serves as a volume marker that facilitates the segmentation of cells when processing microscopy images.

### 2.2 Growth conditions

For all experiments, cultures were initiated from a 50% glycerol frozen stock at −80*°*C. Three strains - autofluorescence (*auto*), Δ*lacI* (Δ), and a strain with a known binding site and repressor copy number (*R*) - were inoculated into individual tubes with 2 mL of Lysogeny Broth (LB Miller Powder, BD Medical) with 20 *µ*g/mL of chloramphenicol and 30 *µ*g/mL of kanamycin. These cultures were grown overnight at 37*°*C with rapid agitation to reach saturation. The saturated cultures were diluted 1:1000 into 500 *µ*L of M9 minimal media (M9 5X Salts, Sigma-Aldrich M6030; 2 mM magnesium sulfate, Mallinckrodt Chemicals 6066-04; 100 mM calcium chloride, Fisher Chemicals C79-500) supplemented with 0.5% (w/v) glucose on a 2 mL 96-deep-well plate. The *R* strain was diluted into 12 different wells with minimal media, each with a different IPTG concentration (0 *µ*M, 0.1 *µ*M, 5 *µ*M, 10 *µ*M, 25 *µ*M, 50 *µ*M, 75 *µ*M, 100 *µ*M, 250 *µ*M, 500 *µM*, 1000 *µ*M, 5000 *µ*M) while the *auto* and Δ strains were diluted into two wells (0 *µ*M, 5000 *µ*M). Each of the IPTG concentrations came from a single preparation stock kept in 100-fold concentrated aliquots. The 96 well plate was then incubated at 37*°*C with rapid agitation for 8 hours before imaging.

### 2.3 Microscopy imaging procedure

The microscopy pipeline used for this work exactly followed the steps from [27]. Briefly, twelve 2% agarose (Life Technologies UltraPure Agarose, Cat.No. 16500100) gels were made out of M9 media (or PBS buffer) with the corresponding IPTG concentration (see growth conditions) and placed between two glass coverslips for them to solidify after microwaving. After the 8 hour incubation in minimal media, 1 *µ*L of a 1:10 dilution of the cultures into fresh media or PBS buffer was placed into small squares (roughly 10 mm × 10 mm) of the different agarose gels. A total of 16 agarose squares - 12 concentrations of IPTG for the *R* strain, 2 concentrations for the Δ and 2 for the *auto* strain – were mounted into a single glass-bottom dish (Ted Pella Wilco Dish, Cat. No. 14027-20) that was sealed with parafilm.

All imaging was done on an inverted fluorescent microscope (Nikon Ti-Eclipse) with custom-built laser illumination system. The YFP fluorescence (quantitative reporter) was imaged with a CrystaLaser 514 nm excitation laser coupled with a laser-optimized (Semrock Cat. No. LF514-C-000) emission filter. All strains, including the *auto* strain, included a constitutively expressed mCherry protein to aid the segmentation. Therefore, for each image three channels (YFP, On average 30 images with roughly 20 cells per condition were taken. 25 images of a fluorescent slide and 25 images of the camera background noise were taken every imaging session in order to flatten the illumination. The image processing pipeline for this work is exactly the same as in [27].

### 2.4 Data and Code Availability

All data and custom scripts were collected and stored using Git version control. Code for raw data processing, theoretical analysis, and figure generation is available on the GitHub repository (https://github.com/RPGroup-PBoC/chann_cap). The code can also be accessed via the paper website (https://www.rpgroup.caltech.edu/chann_cap/). Raw microscopy data are stored on the CaltechDATA data repository and can be accessed via DOI https://doi.org/10.22002/d1.1184. Bootstrap estimates of experimental channel capacity are also available on the CaltechDATA data repository via https://doi.org/10.22002/D1.1185.

## Acknowledgements

We would like to also thank Nathan Belliveau, Michael Betancourt, William Bialek, Justin Bois, Emanuel Flores, Hernan Garcia, Alejandro Granados, Porfirio Quintero, Catherine Triandafillou, and Ned Wingreen for useful advice and discussion. We would especially like to thank Alvaro Sanchez, Gašper Tkačik, and Jane Kondev for critical observations on the manuscript. We thank Rob Brewster for providing the raw mRNA FISH data for inferences, and David Drabold for advice on the maximum entropy inferences. We are grateful to Heun Jin Lee for his key support with the quantitative microscopy. This work was supported by La Fondation Pierre-Gilles de Gennes, the Rosen Center at Caltech, and the NIH 1R35 GM118043 (MIRA). M.R.M. was supported by the Caldwell CEMI fellowship.

