## Supplemental Information for "First-principles prediction of the information processing capacity of a simple genetic circuit"

**S9 Derivation of the cell age distribution** **71**

### S1 Three-state promoter model for simple repression

In order to tackle the question of how much information the simple repression motif can process we require the joint probability distribution of mRNA and protein $P(m, p; t)$. To obtain this distribution we use the chemical master equation formalism as described in Section 1.1. Specifically, we assume a three-state model where the promoter can be found 1) in a transcriptionally active state ($A$ state), 2) in a transcriptionally inactive state without the repressor bound ($I$ state) and 3) with the repressor bound ($R$ state). (See Fig. 2(A)). These three states generate a system of coupled differential equations for each of the three state distributions $P_A(m, p)$, $P_I(m, p)$ and $P_R(m, p)$. Given the rates shown in Fig. 2(A) let us define the system of ODEs. For the transcriptionally active state we have

$$
\begin{aligned}
\frac{dP_A(m,p)}{dt} = & -\overbrace{k_{\text{off}}^{(p)} P_A(m,p)}^{A\to I} + \overbrace{k_{\text{on}}^{(p)} P_I(m,p)}^{I\to A} \\
& + \overbrace{r_m P_A(m-1,p)}^{m-1\to m} - \overbrace{r_m P_A(m,p)}^{m\to m+1} + \overbrace{\gamma_m (m+1) P_A(m+1,p)}^{m+1\to m} - \overbrace{\gamma_m m P_A(m,p)}^{m\to m-1} \\
& + \overbrace{r_p m P_A(m,p-1)}^{p-1\to p} - \overbrace{r_p m P_A(m,p)}^{p\to p+1} + \overbrace{\gamma_p (p+1) P_A(m,p+1)}^{p+1\to p} - \overbrace{\gamma_p p P_A(m,p)}^{p\to p-1}.
\end{aligned}
\tag{S1}
$$

For the inactive promoter state $I$ we have

$$\begin{aligned}\frac{dP_I(m,p)}{dt} = & \overbrace{k_{\text{off}}^{(p)} P_A(m,p)}^{A \to I} - \overbrace{k_{\text{on}}^{(p)} P_I(m,p)}^{I \to A} + \overbrace{k_{\text{off}}^{(r)} P_R(m,p)}^{R \to I} - \overbrace{k_{\text{on}}^{(r)} P_I(m,p)}^{I \to R} \\ & + \overbrace{\gamma_m (m+1) P_I(m+1,p)}^{m+1 \to m} - \overbrace{\gamma_m m P_I(m,p)}^{m \to m-1} \\ & + \overbrace{r_p m P_I(m,p-1)}^{p-1 \to p} - \overbrace{r_p m P_I(m,p)}^{p \to p+1} + \overbrace{\gamma_p (p+1) P_I(m,p+1)}^{p+1 \to p} - \overbrace{\gamma_p p P_I(m,p)}^{p \to p-1} .\end{aligned} \tag{S2}$$

And finally for the repressor bound state $R$ we have

$$\begin{aligned}\frac{dP_R(m,p)}{dt} = & - \overbrace{k_{\text{off}}^{(r)} P_R(m,p)}^{R \to I} + \overbrace{k_{\text{on}}^{(r)} P_I(m,p)}^{I \to R} \\ & + \overbrace{\gamma_m (m+1) P_R(m+1,p)}^{m+1 \to m} - \overbrace{\gamma_m m P_R(m,p)}^{m \to m-1} \\ & + \overbrace{r_p m P_R(m,p-1)}^{p-1 \to p} - \overbrace{r_p m P_R(m,p)}^{p \to p+1} + \overbrace{\gamma_p (p+1) P_R(m,p+1)}^{p+1 \to p} - \overbrace{\gamma_p p P_R(m,p)}^{p \to p-1} .\end{aligned} \tag{S3}$$

For an unregulated promoter, i.e. a promoter in a cell that has no repressors present, and therefore constitutively expresses the gene, we use a two-state model in which the state $R$ is not allowed. All the terms in the system of ODEs containing $k_{\text{on}}^{(r)}$ or $k_{\text{off}}^{(r)}$ are then set to zero.

As detailed in Section 1.1 it is convenient to express this system using matrix notation [30]. For this we define $\mathbf{P}(m,p) = (P_A(m,p), P_I(m,p), P_R(m,p))^T$. Then the system of ODEs can be expressed as

$$\begin{aligned}\frac{d\mathbf{P}(m,p)}{dt} = & \mathbf{K}\mathbf{P}(m,p) - \mathbf{R}_m \mathbf{P}(m,p) + \mathbf{R}_m \mathbf{P}(m-1,p) - m\mathbf{\Gamma}_m \mathbf{P}(m,p) + (m+1)\mathbf{\Gamma}_m \mathbf{P}(m+1,p) \\ & - m\mathbf{R}_p \mathbf{P}(m,p) + m\mathbf{R}_p \mathbf{P}(m,p-1) - p\mathbf{\Gamma}_p \mathbf{P}(m,p) + (p+1)\mathbf{\Gamma}_p \mathbf{P}(m,p+1),\end{aligned} \tag{S4}$$

where we defined matrices representing the promoter state transition $\mathbf{K}$,

$$\mathbf{K} \equiv \begin{bmatrix} -k_{\text{off}}^{(p)} & k_{\text{on}}^{(p)} & 0 \\ k_{\text{off}}^{(p)} & -k_{\text{on}}^{(p)} - k_{\text{on}}^{(r)} & k_{\text{off}}^{(r)} \\ 0 & k_{\text{on}}^{(r)} & -k_{\text{off}}^{(r)} \end{bmatrix}, \tag{S5}$$

mRNA production, $\mathbf{R}_m$, and degradation, $\mathbf{\Gamma}_m$, as

$$\mathbf{R}_m \equiv \begin{bmatrix} r_m & 0 & 0 \\ 0 & 0 & 0 \\ 0 & 0 & 0 \end{bmatrix}, \tag{S6}$$

and

$$\mathbf{\Gamma}_m \equiv \begin{bmatrix} \gamma_m & 0 & 0 \\ 0 & \gamma_m & 0 \\ 0 & 0 & \gamma_m \end{bmatrix}. \tag{S7}$$

For the protein we also define production $\mathbf{R}_p$ and degradation $\mathbf{\Gamma}_p$ matrices as

$$\mathbf{R}_p \equiv \begin{bmatrix} r_p & 0 & 0 \\ 0 & r_p & 0 \\ 0 & 0 & r_p \end{bmatrix} \tag{S8}$$

and

$$\mathbf{\Gamma}_p \equiv \begin{bmatrix} \gamma_p & 0 & 0 \\ 0 & \gamma_p & 0 \\ 0 & 0 & \gamma_p \end{bmatrix}. \tag{S9}$$

The corresponding equation for the unregulated two-state promoter takes the exact same form with the definition of the matrices following the same scheme without including the third row and third column, and setting $k_{\text{on}}^{(r)}$ and $k_{\text{off}}^{(r)}$ to zero.

A closed-form solution for this master equation might not even exist. The approximate solution of chemical master equations of this kind is an active area of research. As we will see in Appendix S2 the two-state promoter master equation has been analytically solved for the mRNA [34] and protein distributions [53]. For our purposes, in Appendix S5 we will detail how to use the Maximum Entropy principle to approximate the full distribution for the two- and three-state promoter.

### S2 Parameter inference

(Note: The Python code used for the calculations presented in this section can be found in the following link as an annotated Jupyter notebook)

With the objective of generating falsifiable predictions with meaningful parameters, we infer the kinetic rates for this three-state promoter model using different data sets generated in our lab over the last decade concerning different aspects of the regulation of the simple repression motif. For example, for the unregulated promoter transition rates $k_{\text{on}}^{(p)}$ and $k_{\text{off}}^{(p)}$ and the mRNA production rate $r_m$, we use single-molecule mRNA FISH counts from an unregulated promoter [25]. Once these parameters are fixed, we use the values to constrain the repressor rates $k_{\text{on}}^{(r)}$ and $k_{\text{off}}^{(r)}$. These repressor rates are obtained using information from mean gene expression measurements from bulk LacZ colorimetric assays [20]. We also expand our model to include the allosteric nature of the repressor protein, taking advantage of video microscopy measurements done in the context of multiple promoter copies [26] and flow-cytometry measurements of the mean response of the system to different levels of induction [27]. In what follows of this section we detail the steps taken to infer the parameter values. At each step the values of the parameters inferred in previous steps constrain the values of the parameters that are not yet determined, building in this way a self-consistent model informed by work that spans several experimental techniques.

#### S2.1 Unregulated promoter rates

We begin our parameter inference problem with the promoter on and off rates $k_{\text{on}}^{(p)}$ and $k_{\text{off}}^{(p)}$, as well as the mRNA production rate $r_m$. In this case there are only two states available to the promoter – the inactive state $I$ and the transcriptionally active state $A$. That means that the third ODE for $P_R(m, p)$ is removed from the system. The mRNA steady state distribution for this particular two-state promoter model was solved analytically by Peccoud and Ycart [34]. This distribution $P(m) \equiv P_I(m) + P_A(m)$ is of the form

$$P(m \mid k_{\text{on}}^{(p)}, k_{\text{off}}^{(p)}, r_m, \gamma_m) = \frac{\Gamma\left(\frac{k_{\text{on}}^{(p)}}{\gamma_m} + m\right)}{\Gamma(m+1)\Gamma\left(\frac{k_{\text{off}}^{(p)}+k_{\text{on}}^{(p)}}{\gamma_m} + m\right)} \frac{\Gamma\left(\frac{k_{\text{off}}^{(p)}+k_{\text{on}}^{(p)}}{\gamma_m}\right)}{\Gamma\left(\frac{k_{\text{on}}^{(p)}}{\gamma_m}\right)} \left(\frac{r_m}{\gamma_m}\right)^m F_1^1\left(\frac{k_{\text{on}}^{(p)}}{\gamma_m} + m, \frac{k_{\text{off}}^{(p)} + k_{\text{on}}^{(p)}}{\gamma_m} + m, -\frac{r_m}{\gamma_m}\right), \tag{S10}$$

where $\Gamma(\cdot)$ is the gamma function, and $F_1^1$ is the confluent hypergeometric function of the first kind. This rather complicated expression will aid us to find parameter values for the rates. The inferred rates $k_{\text{on}}^{(p)}$, $k_{\text{off}}^{(p)}$ and $r_m$ will be expressed in units of the mRNA degradation rate $\gamma_m$. This is because the model in Eq. S10 is homogeneous in time, meaning that if we divide all rates by a constant it would be equivalent to multiplying the characteristic time scale by the same constant. As we will discuss in the next section, Eq. S10 has degeneracy in the parameter values. What this means is that a change in one of the parameters, specifically $r_m$, can be compensated by a change in another parameter, specifically $k_{\text{off}}^{(p)}$, to obtain the exact same distribution. To work around this intrinsic limitation of the model we will include in our inference prior information from what we know from equilibrium-based models.

#### Bayesian parameter inference of RNAP rates

In order to make progress at inferring the unregulated promoter state transition rates, we make use of the single-molecule mRNA FISH data from Jones et al. [25]. Fig. S1 shows the distribution of mRNA per cell for the *lacUV5* promoter used for our inference. This promoter, being very strong, has a mean copy number of $\langle m \rangle \approx 18$ mRNA/cell.

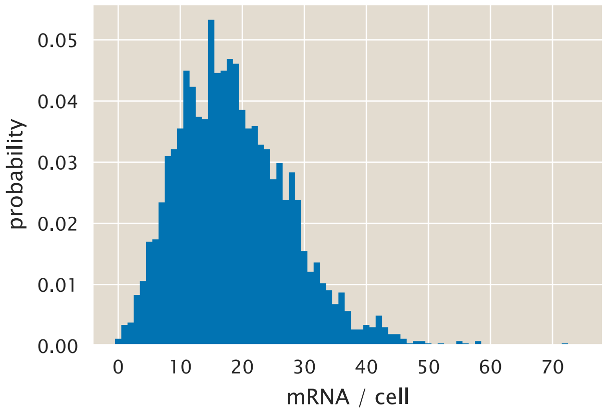

**Figure S1. *lacUV5* mRNA per cell distribution.** Data from [25] of the unregulated *lacUV5* promoter as inferred from single molecule mRNA FISH.

Having this data in hand we now turn to Bayesian parameter inference. Writing Bayes theorem we have

$$P(k_{\text{on}}^{(p)}, k_{\text{off}}^{(p)}, r_m \mid D) = \frac{P(D \mid k_{\text{on}}^{(p)}, k_{\text{off}}^{(p)}, r_m) P(k_{\text{on}}^{(p)}, k_{\text{off}}^{(p)}, r_m)}{P(D)}, \tag{S11}$$

where $D$ represents the data. For this case the data consists of single-cell mRNA counts $D = \{m_1, m_2, \ldots, m_N\}$, where $N$ is the number of cells. We assume that each cell's measurement is independent of the others such that we can rewrite Eq. S11 as

$$P(k_{\text{on}}^{(p)}, k_{\text{off}}^{(p)}, r_m \mid \{m_i\}) \propto \left[\prod_{i=1}^{N} P(m_i \mid k_{\text{on}}^{(p)}, k_{\text{off}}^{(p)}, r_m)\right] P(k_{\text{on}}^{(p)}, k_{\text{off}}^{(p)}, r_m), \tag{S12}$$

where we ignore the normalization constant $P(D)$. The likelihood term $P(m_i \mid k_{\text{on}}^{(p)}, k_{\text{off}}^{(p)}, r_m)$ is exactly given by Eq. S10 with $\gamma_m = 1$. Given that we have this functional form for the distribution, we can use Markov Chain Monte Carlo (MCMC) sampling to explore the 3D parameter space in order to fit Eq. S10 to the mRNA-FISH data.

**Constraining the rates given prior thermodynamic knowledge.**

One of the strengths of the Bayesian approach is that we can include all the prior knowledge on the parameters when performing an inference [4]. Basic features such as the fact that the rates have to be strictly positive constrain the values that these parameters can take. For the specific rates analyzed in this section we know more than the simple constraint of non-negative values. The expression of an unregulated promoter has been studied from a thermodynamic perspective [23]. Given the underlying assumptions of these equilibrium models, in which the probability of finding the RNAP bound to the promoter is proportional to the transcription rate [54], they can only make statements about the mean expression level. Nevertheless if both the thermodynamic and the kinetic model describe the same process, the predictions for the mean gene expression level must agree. That means that we can use what we know about the mean gene expression, and how this is related to parameters such as molecule copy numbers and binding affinities, to constrain the values that the rates in question can take.

In the case of this two-state promoter it can be shown that the mean number of mRNA is given by [30] (See Appendix S3 for moment computation)

$$\langle m \rangle = \frac{r_m}{\gamma_m} \frac{k_{\text{on}}^{(p)}}{k_{\text{on}}^{(p)} + k_{\text{off}}^{(p)}}. \tag{S13}$$

Another way of expressing this is as $\frac{r_m}{\gamma_m} \times p_{\text{active}}^{(p)}$, where $p_{\text{active}}^{(p)}$ is the probability of the promoter being in the transcriptionally active state. The thermodynamic picture has an equivalent result where the mean number of mRNA is given by [23, 54]

$$\langle m \rangle = \frac{r_m}{\gamma_m} \frac{\frac{P}{N_{\text{NS}}} e^{-\beta \Delta\varepsilon_p}}{1 + \frac{P}{N_{\text{NS}}} e^{-\beta \Delta\varepsilon_p}}, \tag{S14}$$

where $P$ is the number of RNAP per cell, $N_{\text{NS}}$ is the number of non-specific binding sites, $\Delta\varepsilon_p$ is the RNAP binding energy in $k_BT$ units and $\beta \equiv (k_BT)^{-1}$. Using Eq. S13 and Eq. S14 we can easily see that if these frameworks are to be equivalent, then it must be true that

$$\frac{k_{\text{on}}^{(p)}}{k_{\text{off}}^{(p)}} = \frac{P}{N_{\text{NS}}} e^{-\beta \Delta\varepsilon_p}, \tag{S15}$$

or equivalently

$$\ln\left(\frac{k_{\text{on}}^{(p)}}{k_{\text{off}}^{(p)}}\right) = -\beta \Delta\varepsilon_p + \ln P - \ln N_{\text{NS}}. \tag{S16}$$

To put numerical values into these variables we can use information from the literature. The RNAP copy number is order $P \approx 1000-3000$ RNAP/cell for a 1 hour doubling time [55]. As for the number of non-specific binding sites and the binding energy, we have that $N_{\text{NS}} = 4.6\times10^6$ [54] and $-\beta\Delta\varepsilon_p \approx 5-7$ [23]. Given these values we define a Gaussian prior for the log ratio of these two quantities of the form

$$P\left(\ln\left(\frac{k_{\text{on}}^{(p)}}{k_{\text{off}}^{(p)}}\right)\right) \propto \exp\left\{-\frac{\left(\ln\left(\frac{k_{\text{on}}^{(p)}}{k_{\text{off}}^{(p)}}\right) - (-\beta\Delta\varepsilon_p + \ln P - \ln N_{\text{NS}})\right)^2}{2\sigma^2}\right\}, \tag{S17}$$

where $\sigma$ is the variance that accounts for the uncertainty in these parameters. We include this prior as part of the prior term $P(k_{\text{on}}^{(p)}, k_{\text{off}}^{(p)}, r_m)$ of Eq. S12. We then use MCMC to sample out of the posterior distribution given by Eq. S12. Fig. S2 shows the MCMC samples of the posterior distribution. For the case of the $k_{\text{on}}^{(p)}$ parameter there is a single symmetric peak. $k_{\text{off}}^{(p)}$ and $r_m$ have a rather long tail towards large values. In fact, the 2D projection of $k_{\text{off}}^{(p)}$ vs $r_m$ shows that the model is sloppy, meaning that the two parameters are highly correlated. This feature is a common problem for many non-linear systems used in biophysics and systems biology [56]. What this implies is that we can change the value of $k_{\text{off}}^{(p)}$, and then compensate by a change in $r_m$ in order to maintain the shape of the mRNA distribution. Therefore it is impossible from the data and the model themselves to narrow down a single value for the parameters. Nevertheless since we included the prior information on the rates as given by the analogous form between the equilibrium and non-equilibrium expressions for the mean mRNA level, we obtained a more constrained parameter value for the RNAP rates and the transcription rate that we will take as the peak of this long-tailed distribution.

The inferred values $k_{\text{on}}^{(p)} = 4.3^{+1}_{-0.3}$, $k_{\text{off}}^{(p)} = 18.8^{+120}_{-10}$ and $r_m = 103.8^{+423}_{-37}$ are given in units of the mRNA degradation rate $\gamma_m$. Given the asymmetry of the parameter distributions we report the upper and lower bound of the 95$^{\text{th}}$ percentile of the posterior distributions. Assuming a mean life-time for mRNA of $\approx 3$ min (from this link) we have an mRNA degradation rate of $\gamma_m \approx 2.84\times10^{-3}s^{-1}$. Using this value gives the following values for the inferred rates: $k_{\text{on}}^{(p)} = 0.024^{+0.005}_{-0.002}s^{-1}$, $k_{\text{off}}^{(p)} = 0.11^{+0.66}_{-0.05}s^{-1}$, and $r_m = 0.3^{+2.3}_{-0.2}s^{-1}$.

Fig. S3 compares the experimental data from Fig. S1 with the resulting distribution obtained by substituting the most likely parameter values into Eq. S10. As we can see this two-state model fits the data adequately.

#### S2.2 Accounting for variability in the number of promoters

As discussed in ref. [25] and further expanded in [38] an important source of cell-to-cell variability in gene expression in bacteria is the fact that, depending on the growth rate and the position relative to the chromosome replication origin, cells can have multiple copies of any given gene. Genes closer to the replication origin have on average higher gene copy number compared to genes at the opposite end. For the locus in which our reporter construct is located (*galK*) and the doubling time of the mRNA FISH experiments we expect to have $\approx 1.66$ copies of the gene [25, 37]. This implies that the cells spend 2/3 of the cell cycle with two copies of the promoter and the rest with a single copy.

To account for this variability in gene copy we extend the model assuming that when cells have two copies of the promoter the mRNA production rate is $2r_m$ compared to the rate $r_m$ for a single

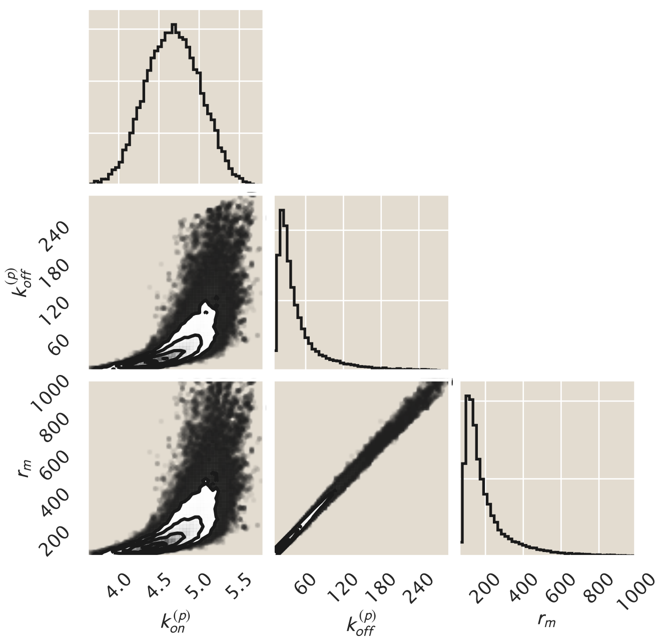

**Figure S2. MCMC posterior distribution.** Sampling out of Eq. S12 the plot shows 2D and 1D projections of the 3D parameter space. The parameter values are (in units of the mRNA degradation rate $\gamma_m$) $k^{(p)}_{\text{on}} = 4.3^{+1}_{-0.3}$, $k^{(p)}_{\text{off}} = 18.8^{+120}_{-10}$ and $r_m = 103.8^{+423}_{-37}$ which are the modes of their respective distributions, where the superscripts and subscripts represent the upper and lower bounds of the 95$^{\text{th}}$ percentile of the parameter value distributions

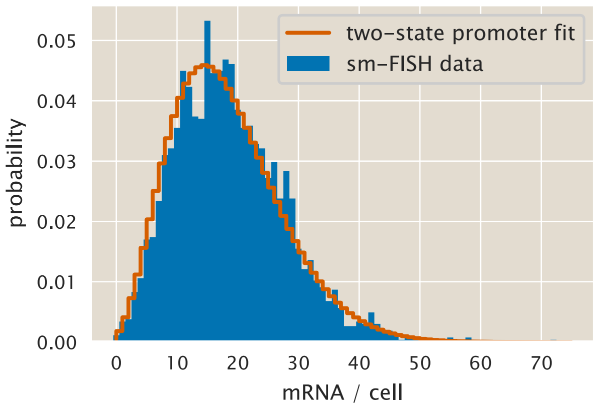

**Figure S3. Experimental vs. theoretical distribution of mRNA per cell using parameters from Bayesian inference.** Dotted line shows the result of using Eq. S10 along with the parameters inferred for the rates. Blue bars are the same data as Fig. S1 obtained from [25].

promoter copy. The probability of observing a certain mRNA copy $m$ is therefore given by

$$P(m) = P(m \mid \text{one promoter}) \cdot P(\text{one promoter}) + P(m \mid \text{two promoters}) \cdot P(\text{two promoters}). \quad \text{(S18)}$$

Both terms $P(m \mid \text{promoter copy})$ are given by Eq. S10 with the only difference being the rate $r_m$. It is important to acknowledge that Eq. S18 assumes that once the gene is replicated the time scale in which the mRNA count relaxes to the new steady state is much shorter than the time that the cells spend in this two promoter copies state. This approximation should be valid for a short lived mRNA molecule, but the assumption is not applicable for proteins whose degradation rate is comparable to the cell cycle length as explored in Section 1.4.

In order to repeat the Bayesian inference including this variability in gene copy number we must split the mRNA count data into two sets – cells with a single copy of the promoter and cells with two copies of the promoter. For the single molecule mRNA FISH data there is no labeling of the locus, making it impossible to determine the number of copies of the promoter for any given cell. We therefore follow Jones et al. [25] in using the cell area as a proxy for stage in the cell cycle. In their approach they sorted cells by area, considering cells below the 33$^{\text{th}}$ percentile having a single promoter copy and the rest as having two copies. This approach ignores that cells are not uniformly distributed along the cell cycle. As first derived in [40] populations of cells in a log-phase are exponentially distributed along the cell cycle. This distribution is of the form

$$P(a) = (\ln 2) \cdot 2^{1-a}, \tag{S19}$$

where $a \in [0, 1]$ is the stage of the cell cycle, with $a = 0$ being the start of the cycle and $a = 1$ being the cell division (See Appendix S9 for a derivation of Eq. S19). Fig. S4 shows the separation of the two groups based on area where Eq. S19 was used to weight the distribution along the cell cycle.

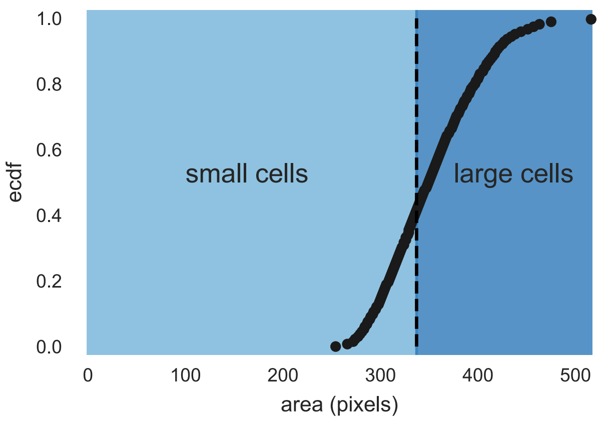

**Figure S4. Separation of cells based on cell size.** Using the area as a proxy for position in the cell cycle, cells can be sorted into two groups – small cells (with one promoter copy) and large cells (with two promoter copies). The vertical black line delimits the threshold that divides both groups as weighted by Eq. S19.

A subtle, but important consequence of Eq. S19 is that computing any quantity for a single cell is not equivalent to computing the same quantity for a population of cells. For example, let us assume that we want to compute the mean mRNA copy number $\langle m \rangle$. For a single cell this would be of the form

$$\langle m \rangle_{\text{cell}} = \langle m \rangle_1 \cdot f + \langle m \rangle_2 \cdot (1 - f), \tag{S20}$$

where $\langle m \rangle_i$ is the mean mRNA copy number with $i$ promoter copies in the cell, and $f$ is the fraction of the cell cycle that cells spend with a single copy of the promoter. For a single cell the probability of having a single promoter copy is equivalent to this fraction $f$. But Eq. S19 tells us that if we sample unsynchronized cells we are not sampling uniformly across the cell cycle. Therefore for a population of cells the mean mRNA is given by

$$\langle m \rangle_{\text{population}} = \langle m \rangle_1 \cdot \phi + \langle m \rangle_2 \cdot (1 - \phi) \tag{S21}$$

where the probability of sampling a cell with one promoter $\phi$ is given by

$$\phi = \int_0^f P(a) da, \tag{S22}$$

where $P(a)$ is given by Eq. S106. What this equation computes is the probability of sampling a cell during a stage of the cell cycle $< f$ where the reporter gene hasn't been replicated yet. Fig. S5 shows the distribution of both groups. As expected larger cells have a higher mRNA copy number on average.

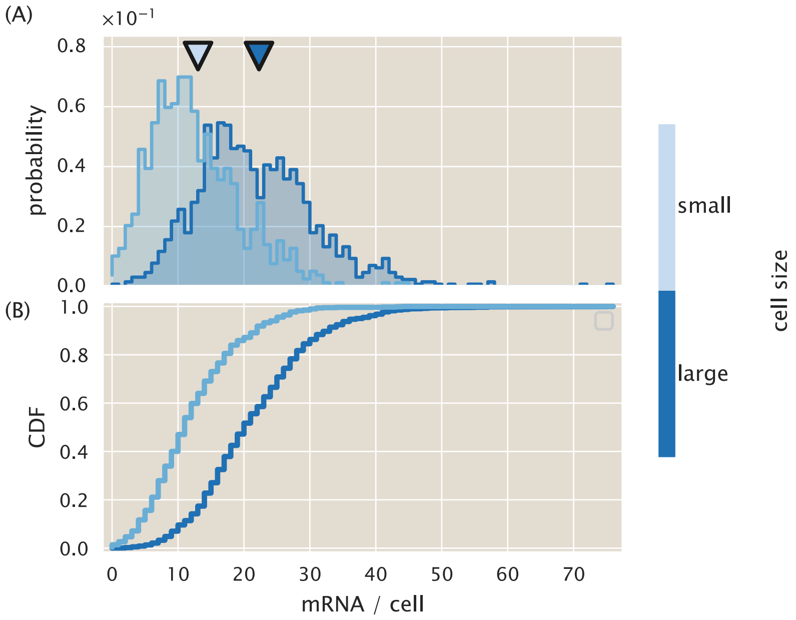

**Figure S5. mRNA distribution for small and large cells.** (A) histogram and (B) cumulative distribution function of the small and large cells as determined in Fig. S4. The triangles above histograms in (A) indicate the mean mRNA copy number for each group.

We modify Eq. S12 to account for the two separate groups of cells. Let $N_s$ be the number of cells in the small size group and $N_l$ the number of cells in the large size group. Then the posterior distribution for the parameters is of the form

$$P(k_{\text{on}}^{(p)}, k_{\text{off}}^{(p)}, r_m \mid \{m_i\}) \propto \left[ \prod_{i=1}^{N_s} P(m_i \mid k_{\text{on}}^{(p)}, k_{\text{off}}^{(p)}, r_m) \right] \left[ \prod_{j=1}^{N_l} P(m_j \mid k_{\text{on}}^{(p)}, k_{\text{off}}^{(p)}, 2r_m) \right] P(k_{\text{on}}^{(p)}, k_{\text{off}}^{(p)}, r_m), \tag{S23}$$

where we split the product of small and large cells.

For the two-promoter model the prior shown in Eq. S17 requires a small modification. Eq. S21 gives the mean mRNA copy number of a population of asynchronous cells growing at steady state. Given that we assume that the only difference between having one vs. two promoter copies is the change in transcription rate from $r_m$ in the single promoter case to $2r_m$ in the two-promoter case we can write Eq. S21 as

$$\langle m \rangle = \phi \cdot \frac{r_m}{\gamma_m} \frac{k_{\text{on}}^{(p)}}{k_{\text{on}}^{(p)} + k_{\text{off}}^{(p)}} + (1-\phi) \cdot \frac{2r_m}{\gamma_m} \frac{k_{\text{on}}^{(p)}}{k_{\text{on}}^{(p)} + k_{\text{off}}^{(p)}}. \tag{S24}$$

This can be simplified to

$$\langle m \rangle = (2-\phi) \frac{r_m}{\gamma_m} \frac{k_{\text{on}}^{(p)}}{k_{\text{on}}^{(p)} + k_{\text{off}}^{(p)}}. \tag{S25}$$

Equating Eq. S25 and Eq. S14 to again require self-consistent predictions of the mean mRNA from the equilibrium and kinetic models gives

$$(2-\phi) \frac{k_{\text{on}}^{(p)}}{k_{\text{on}}^{(p)} + k_{\text{off}}^{(p)}} = \frac{\frac{P}{N_{\text{NS}}} e^{-\beta\Delta\varepsilon_p}}{1 + \frac{P}{N_{\text{NS}}} e^{-\beta\Delta\varepsilon_p}}. \tag{S26}$$

Solving for $\frac{k_{\text{on}}^{(p)}}{k_{\text{off}}^{(p)}}$ results in

$$\left( \frac{k_{\text{on}}^{(p)}}{k_{\text{off}}^{(p)}} \right) = \frac{\rho}{[(1+\rho)(2-\phi) - \rho]}, \tag{S27}$$

where we define $\rho \equiv \frac{P}{N_{\text{NS}}} e^{-\beta\Delta\varepsilon_p}$. To simplify things further we notice that for the specified values of $P = 1000 - 3000$ per cell, $N_{\text{NS}} = 4.6 \times 10^6$ bp, and $-\beta\Delta\varepsilon_p = 5 - 7$, we can safely assume that $\rho \ll 1$. This simplifying assumption has been previously called the weak promoter approximation [20]. Given this we can simplify Eq. S27 as

$$\frac{k_{\text{on}}^{(p)}}{k_{\text{off}}^{(p)}} = \frac{1}{2-\phi} \frac{P}{N_{\text{NS}}} e^{-\beta\Delta\varepsilon_p}. \tag{S28}$$

Taking the log of both sides gives

$$\ln \left( \frac{k_{\text{on}}^{(p)}}{k_{\text{off}}^{(p)}} \right) = -\ln(2-\phi) + \ln P - \ln N_{\text{NS}} - \beta\Delta\varepsilon_p. \tag{S29}$$

With this we can set as before a Gaussian prior to constrain the ratio of the RNAP rates as

$$P \left( \ln \left( \frac{k_{\text{on}}^{(p)}}{k_{\text{off}}^{(p)}} \right) \right) \propto \exp \left\{ - \frac{\left( \ln \left( \frac{k_{\text{on}}^{(p)}}{k_{\text{off}}^{(p)}} \right) - \left[ -\ln(2-\phi) - \beta\Delta\varepsilon_p + \ln P - \ln N_{\text{NS}} \right) \right]^2}{2\sigma^2} \right\}. \tag{S30}$$

Fig. S6 shows the result of sampling out of Eq. S23. Again we see that the model is highly sloppy with large credible regions obtained for $k_{\text{off}}^{(p)}$ and $r_m$. Nevertheless, again the use of the prior information allows us to get a parameter values consistent with the equilibrium picture.

Using again a mRNA mean lifetime of $\approx 3$ min gives the following values for the parameters: $k_{\text{on}}^{(p)} = 0.03^{+0.004}_{-0.002} s^{-1}$, $k_{\text{off}}^{(p)} = 0.7^{+4.1}_{-0.4} s^{-1}$, and $r_m = 1.4^{+7.3}_{-0.7} s^{-1}$. Fig. S7 shows the result of applying Eq. S18

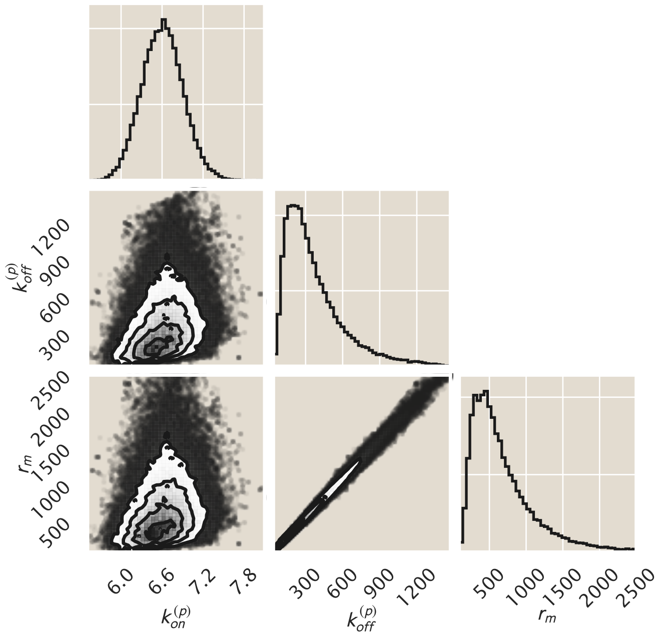

**Figure S6. MCMC posterior distribution for a multi-promoter model.** Sampling out of Eq. S23 the plot shows 2D and 1D projections of the 3D parameter space. The parameter values are (in units of the mRNA degradation rate $\gamma_m$) $k_{\text{on}}^{(p)} = 6.4^{+0.8}_{-0.4}$, $k_{\text{off}}^{(p)} = 132^{+737}_{-75}$ and $r_m = 257^{+1307}_{-132}$ which are the modes of their respective distributions, where the superscripts and subscripts represent the upper and lower bounds of the $95^{\text{th}}$ percentile of the parameter value distributions. The sampling was bounded to values $< 1000$ for numerical stability when computing the confluent hypergeometric function.

using these parameter values. Specifically Fig. S7(A) shows the global distribution including cells with one and two promoters and Fig. S7(B) splits the distributions within the two populations. Given that the model adequately describes both populations independently and pooled together we confirm that using the cell area as a proxy for stage in the cell cycle and the doubling of the transcription rate once cells have two promoters are reasonable approximations.

It is hard to make comparisons with literature reported values because these kinetic rates are effective parameters hiding a lot of the complexity of transcription initiation [31]. Also the non-identifiability of the parameters restricts our explicit comparison of the actual numerical values of the inferred rates. Nevertheless from the model we can see that the mean burst size for each transcription event is given by $r_m/k_{\text{off}}^{(p)}$. From our inferred values we obtain then a mean burst size of $\approx 1.9$ transcripts per cell. This is similar to the reported burst size of 1.15 on a similar system on *E. coli* [57].

#### S2.3 Repressor rates from three-state regulated promoter.

Having determined the unregulated promoter transition rates we now proceed to determine the repressor rates $k_{\text{on}}^{(r)}$ and $k_{\text{off}}^{(r)}$. The values of these rates are constrained again by the correspondence between our kinetic picture and what we know from equilibrium models [36]. For this analysis we again exploit the feature that, at the mean, both the kinetic language and the thermodynamic language should have equivalent predictions. Over the last decade there has been great effort in developing

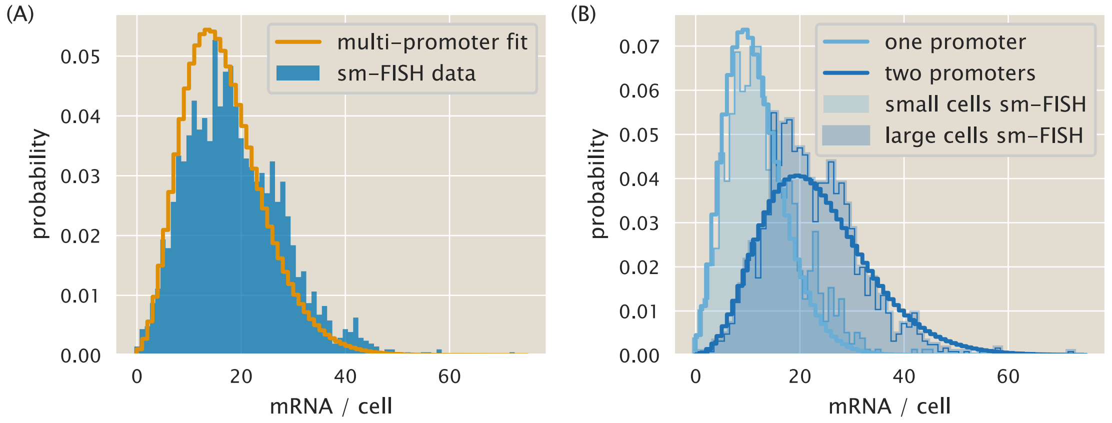

**Figure S7. Experimental vs. theoretical distribution of mRNA per cell using parameters for multi-promoter model.** (A) Solid line shows the result of using Eq. S18 with the parameters inferred by sampling Eq. S23. Blue bars are the same data as Fig. S1 from [25]. (B) Split distributions of small cells (light blue bars) and large cells (dark blue) with the corresponding theoretical predictions with transcription rate $r_m$ (light blue line) and transcription rate $2r_m$ (dark blue line)

equilibrium models for gene expression regulation [33, 54, 58]. In particular our group has extensively characterized the simple repression motif using this formalism [20, 26, 27].

The dialogue between theory and experiments has led to simplified expressions that capture the phenomenology of the gene expression response as a function of natural variables such as molecule count and affinities between molecular players. A particularly interesting quantity for the simple repression motif used by Garcia & Phillips [20] is the fold-change in gene expression, defined as

$$\text{fold-change} = \frac{\langle \text{gene expression}(R \neq 0) \rangle}{\langle \text{gene expression}(R = 0) \rangle}, \tag{S31}$$

where $R$ is the number of repressors per cell and $\langle \cdot \rangle$ is the population average. The fold-change is simply the mean expression level in the presence of the repressor relative to the mean expression level in the absence of regulation. In the language of statistical mechanics this quantity takes the form

$$\text{fold-change} = \left( 1 + \frac{R}{N_{\text{NS}}} e^{-\beta \Delta\varepsilon_r} \right)^{-1}, \tag{S32}$$

where $\Delta\varepsilon_r$ is the repressor-DNA binding energy, and as before $N_{\text{NS}}$ is the number of non-specific binding sites where the repressor can bind [20].

To compute the fold-change in the chemical master equation language we compute the first moment of the steady sate mRNA distribution $\langle m \rangle$ for both the three-state promoter ($R \neq 0$) and the two-state promoter case ($R = 0$) (See Appendix S3 for moment derivation). The unregulated (two-state) promoter mean mRNA copy number is given by Eq. S25. For the regulated (three-state) promoter we have an equivalent expression of the form

$$\langle m(R \neq 0) \rangle = (2 - \phi) \frac{r_m}{\gamma_m} \frac{k^{(r)}_{\text{off}} k^{(p)}_{\text{on}}}{k^{(p)}_{\text{off}} k^{(r)}_{\text{off}} + k^{(p)}_{\text{off}} k^{(r)}_{\text{on}} + k^{(r)}_{\text{off}} k^{(p)}_{\text{on}}}. \tag{S33}$$

Computing the fold-change then gives

$$\text{fold-change} = \frac{\langle m(R \neq 0)\rangle}{\langle m(R = 0)\rangle} = \frac{k_{\text{off}}^{(r)}\left(k_{\text{off}}^{(p)} + k_{\text{on}}^{(p)}\right)}{k_{\text{off}}^{(p)}k_{\text{on}}^{(r)} + k_{\text{off}}^{(r)}\left(k_{\text{off}}^{(p)} + k_{\text{on}}^{(p)}\right)}, \tag{S34}$$

where the factor $(2-\phi)$ due to the multiple promoter copies, the transcription rate $r_m$ and the mRNA degradation rate $\gamma_m$ cancel out.

Given that the number of repressors per cell $R$ is an experimental variable that we can control, we assume that the rate at which the promoter transitions from the transcriptionally inactive state to the repressor bound state, $k_{\text{on}}^{(r)}$, is given by the concentration of repressors $[R]$ times a diffusion limited on rate $k_o$. For the diffusion limited constant $k_o$ we use the value used by Jones et al. [25]. With this in hand we can rewrite Eq. S34 as

$$\text{fold-change} = \left(1 + \frac{k_0[R]}{k_{\text{off}}^{(r)}}\frac{k_{\text{off}}^{(p)}}{k_{\text{on}}^{(p)} + k_{\text{off}}^{(p)}}\right)^{-1}. \tag{S35}$$

We note that both Eq. S32 and Eq. S35 have the same functional form. Therefore if both languages predict the same output for the mean gene expression level, it must be true that

$$\frac{k_o[R]}{k_{\text{off}}^{(r)}}\frac{k_{\text{off}}^{(p)}}{k_{\text{on}}^{(p)} + k_{\text{off}}^{(p)}} = \frac{R}{N_{\text{NS}}}e^{-\beta\Delta\varepsilon_r}. \tag{S36}$$

Solving for $k_{\text{off}}^{(r)}$ gives

$$k_{\text{off}}^{(r)} = \frac{k_o[R]N_{\text{NS}}e^{\beta\Delta\varepsilon_r}}{R}\frac{k_{\text{off}}^{(p)}}{k_{\text{on}}^{(p)} + k_{\text{off}}^{(p)}}. \tag{S37}$$

Since the reported value of $k_o$ is given in units of $\text{nM}^{-1}\text{s}^{-1}$ in order for the units to cancel properly the repressor concentration has to be given in nM rather than absolute count. If we consider that the repressor concentration is equal to

$$[R] = \frac{R}{V_{cell}} \cdot \frac{1}{N_A}, \tag{S38}$$

where $R$ is the absolute repressor copy number per cell, $V_{cell}$ is the cell volume and $N_A$ is Avogadro's number. The *E. coli* cell volume is 2.1 fL [59], and Avogadro's number is $6.022 \times 10^{23}$. If we further include the conversion factor to turn M into nM we find that

$$[R] = \frac{R}{2.1 \times 10^{-15}L} \cdot \frac{1}{6.022 \times 10^{23}} \cdot \frac{10^9 \text{ nmol}}{1 \text{ mol}} \approx 0.8 \times R. \tag{S39}$$

Using this we simplify Eq. S37 as

$$k_{\text{off}}^{(r)} \approx 0.8 \cdot k_o \cdot N_{\text{NS}}e^{\beta\Delta\varepsilon_r} \cdot \frac{k_{\text{off}}^{(p)}}{k_{\text{on}}^{(p)} + k_{\text{off}}^{(p)}}. \tag{S40}$$

What Eq. S40 shows is the direct relationship that must be satisfied if the equilibrium model is set to be consistent with the non-equilibrium kinetic picture. Table S1 summarizes the values obtained for

**Table S1. Binding sites and corresponding parameters.**

| Operator | $\Delta\varepsilon_r$ $(k_BT)$ | $k_{\text{off}}^{(r)}$ $(s^{-1})$ |
|---|---|---|
| O1 | -15.3 | 0.002 |
| O2 | -13.9 | 0.008 |
| O3 | -9.7 | 0.55 |

the three operator sequences used throughout this work. To compute these numbers the number of non-specific binding sites $N_{\text{NS}}$ was taken to be $4.6 \times 10^6$ bp, i.e. the size of the *E. coli* K12 genome.

*In-vivo* measurements of the Lac repressor off rate have been done with single-molecule resolution [60]. The authors report a mean residence time of 5.3±0.2 minutes for the repressor on an O1 operator. The corresponding rate is $k_{\text{off}}^{(r)} \approx 0.003$ $(s^{-1})$, very similar value to what we inferred from our model. In this same reference the authors determined that on average the repressor takes 30.9±0.5 seconds to bind to the operator [60]. Given the kinetic model presented in Fig. 2(A) this time can be converted to the probability of not being on the repressor bound state $P_{\text{not } R}$. This is computed as

$$P_{\text{not } R} = \frac{\tau_{\text{not } R}}{\tau_{\text{not } R} + \tau_R}, \tag{S41}$$

where $\tau_{\text{not } R}$ is the average time that the operator is not occupied by the repressor and $\tau_R$ is the average time that the repressor spends bound to the operator. Substituting the numbers from [60] gives $P_{\text{not } R} \approx 0.088$. From our model we can compute the zeroth moment $\left\langle m^0 p^0 \right\rangle$ for each of the three promoter states. This moment is equivalent to the probability of being on each of the promoter states. Upon substitution of our inferred rate parameters we can compute $P_{\text{not } R}$ as

$$P_{\text{not } R} = 1 - P_R \approx 0.046, \tag{S42}$$

where $P_R$ is the probability of the promoter being bound by the repressor. The value we obtained is within a factor of two from the one reported in [60].

### S3 Computing moments from the master equation

In this section we will compute the moment equations for the distribution $P(m, p)$. Without lost of generality here we will focus on the three-state regulated promoter. The computation of the moments for the two-state promoter follows the exact same procedure, changing only the definition of the matrices in the master equation.

#### S3.1 Computing moments of a distribution

(Note: The Python code used for the calculations presented in this section can be found in the following link as an annotated Jupyter notebook)

To compute any moment of our chemical master equation (Eq. 6) let us define a vector

$$\langle \mathbf{m^x p^y} \rangle \equiv (\langle m^x p^y \rangle_A, \langle m^x p^y \rangle_I, \langle m^x p^y \rangle_R)^T, \tag{S43}$$

where $\langle m^x p^y \rangle_S$ is the expected value of $m^x p^y$ in state $S \in \{A, I, R\}$ with $x, y \in \mathbb{N}$. In other words, just as we defined the vector $\mathbf{P}(m, p)$, here we define a vector to collect the expected value of each of the promoter states. By definition, these moments $\langle m^x p^y \rangle_S$ are computed as

$$\langle m^x p^y \rangle_S \equiv \sum_{m=0}^{\infty} \sum_{p=0}^{\infty} m^x p^y P_S(m, p). \tag{S44}$$

To simplify the notation, let $\sum_x \equiv \sum_{x=0}^{\infty}$. Since we are working with a system of three ODEs, one for each state, let us define the following operation:

$$\langle \mathbf{m^x p^y} \rangle = \sum_m \sum_p m^x p^y \mathbf{P}(m, p) \equiv \begin{bmatrix} \sum_m \sum_p m^x p^y P_A(m, p) \\ \sum_m \sum_p m^x p^y P_I(m, p) \\ \sum_m \sum_p m^x p^y P_R(m, p) \end{bmatrix}. \tag{S45}$$

With this in hand we can then apply this sum over $m$ and $p$ to Eq. 6. For the left-hand side we have

$$\sum_m \sum_p m^x p^y \frac{d\mathbf{P}(m, p)}{dt} = \frac{d}{dt} \left[ \sum_m \sum_p m^x p^y \mathbf{P}(m, p) \right], \tag{S46}$$

where we made use of the linearity property of the derivative to switch the order between the sum and the derivative. Notice that the right-hand side of Eq. S46 contains the definition of a moment from Eq. S44. That means that we can rewrite it as

$$\frac{d}{dt} \left[ \sum_m \sum_p m^x p^y \mathbf{P}(m, p) \right] = \frac{d\langle \mathbf{m^x p^y} \rangle}{dt}. \tag{S47}$$

Distributing the sum on the right-hand side of Eq. 6 gives

$$\begin{aligned} \frac{d\langle \mathbf{m^x p^y} \rangle}{dt} = & \mathbf{K} \sum_m \sum_p m^x p^y \mathbf{P}(m, p) \\ & - \mathbf{R}_m \sum_m \sum_p m^x p^y \mathbf{P}(m, p) + \mathbf{R}_m \sum_m \sum_p m^x p^y \mathbf{P}(m-1, p) \\ & - \mathbf{\Gamma}_m \sum_m \sum_p (m) m^x p^y \mathbf{P}(m, p) + \mathbf{\Gamma}_m \sum_m \sum_p (m+1) m^x p^y \mathbf{P}(m+1, p) \\ & - \mathbf{R}_p \sum_m \sum_p (m) m^x p^y \mathbf{P}(m, p) + \mathbf{R}_p \sum_m \sum_p (m) m^x p^y \mathbf{P}(m, p-1) \\ & - \mathbf{\Gamma}_p \sum_m \sum_p (p) m^x p^y \mathbf{P}(m, p) + \mathbf{\Gamma}_p \sum_m \sum_p (p+1) m^x p^y \mathbf{P}(m, p+1). \end{aligned} \tag{S48}$$

Let's look at each term on the right-hand side individually. For the terms in Eq. S48 involving $\mathbf{P}(m, p)$ we can again use Eq. S44 to rewrite them in a more compact form. This means that we can rewrite the state transition term as

$$\mathbf{K} \sum_m \sum_p m^x p^y \mathbf{P}(m, p) = \mathbf{K} \langle \mathbf{m^x p^y} \rangle. \tag{S49}$$

The mRNA production term involving $\mathbf{P}(m, p)$ can be rewritten as

$$\mathbf{R}_m \sum_m \sum_p m^x p^y \mathbf{P}(m, p) = \mathbf{R}_m \langle \mathbf{m^x p^y} \rangle. \tag{S50}$$

In the same way the mRNA degradation term gives

$$\mathbf{\Gamma}_m \sum_m \sum_p (m) m^x p^y \mathbf{P}(m, p) = \mathbf{\Gamma}_m \left\langle \mathbf{m^{(x+1)} p^y} \right\rangle. \tag{S51}$$

For the protein production and degradation terms involving $\mathbf{P}(m, p)$ we have

$$\mathbf{R}_p \sum_m \sum_p (m) m^x p^y \mathbf{P}(m, p) = \mathbf{R}_p \left\langle \mathbf{m^{(x+1)} p^y} \right\rangle, \tag{S52}$$

and

$$\mathbf{\Gamma}_p \sum_m \sum_p (p) m^x p^y \mathbf{P}(m, p) = \mathbf{\Gamma}_p \left\langle \mathbf{m^x p^{(y+1)}} \right\rangle, \tag{S53}$$

respectively.

For the sums terms in Eq. S48 involving $\mathbf{P}(m \pm 1, p)$ or $\mathbf{P}(m, p \pm 1)$ we can reindex the sum to work around this mismatch. To be more specific let's again look at each term case by case. For the mRNA production term involving $\mathbf{P}(m - 1, p)$ we define $m' \equiv m - 1$. Using this we write

$$\mathbf{R}_m \sum_m \sum_p m^x p^y \mathbf{P}(m - 1, p) = \mathbf{R}_m \sum_{m'=-1}^{\infty} \sum_p (m' + 1)^x p^y \mathbf{P}(m', p). \tag{S54}$$

Since having negative numbers of mRNA or protein doesn't make physical sense we have that $\mathbf{P}(-1, p) = 0$. Therefore we can rewrite the sum starting from 0 rather than from -1, obtaining

$$\mathbf{R}_m \sum_{m'=-1}^{\infty} \sum_p (m' + 1)^x p^y \mathbf{P}(m', p) = \mathbf{R}_m \sum_{m'=0}^{\infty} \sum_p (m' + 1)^x p^y \mathbf{P}(m', p). \tag{S55}$$

Recall that our distribution $\mathbf{P}(m, p)$ takes $m$ and $p$ as numerical inputs and returns a probability associated with such a molecule count. Nevertheless, $m$ and $p$ themselves are dimensionless quantities that serve as indices of how many molecules are in the cell. The distribution is the same whether the variable is called $m$ or $m'$; for a specific number, let's say $m = 5$, or $m' = 5$, $\mathbf{P}(5, p)$ will return the same result. This means that the variable name is arbitrary, and the right-hand side of Eq. S55 can be written as

$$\mathbf{R}_m \sum_{m'=0}^{\infty} \sum_p (m' + 1)^x p^y \mathbf{P}(m', p) = \mathbf{R}_m \langle (\mathbf{m} + \mathbf{1})^{\mathbf{x}} \mathbf{p^y} \rangle, \tag{S56}$$

since the left-hand side corresponds to the definition of a moment.

For the mRNA degradation term involving $\mathbf{P}(m + 1, p)$ we follow a similar procedure in which we define $m' = m + 1$ to obtain

$$\mathbf{\Gamma}_m \sum_m \sum_p (m + 1) m^x p^y \mathbf{P}(m + 1, p) = \mathbf{\Gamma}_m \sum_{m'=1}^{\infty} \sum_p m' (m' - 1)^x p^y \mathbf{P}(m', p). \tag{S57}$$

In this case since the term on the right-hand side of the equation is multiplied by $m'$, starting the sum over $m'$ from 0 rather than from 1 will not affect the result since this factor will not contribute to the total sum. Nevertheless this is useful since our definition of a moment from Eq. S44 requires the sum to start at zero. This means that we can rewrite this term as

$$\mathbf{\Gamma}_m \sum_{m'=1}^{\infty} m' \sum_p (m'-1)^x p^y \mathbf{P}(m', p) = \mathbf{\Gamma}_m \sum_{m'=0}^{\infty} m' \sum_p (m'-1)^x p^y \mathbf{P}(m', p). \tag{S58}$$

Here again we can change the arbitrary label $m'$ back to $m$ obtaining

$$\mathbf{\Gamma}_m \sum_{m'=0}^{\infty} m' \sum_p (m'-1)^x p^y \mathbf{P}(m', p) = \mathbf{\Gamma}_m \langle \mathbf{m}(\mathbf{m}-\mathbf{1})^{\mathbf{x}} \mathbf{p}^{\mathbf{y}} \rangle. \tag{S59}$$

The protein production term involving $\mathbf{P}(m, p-1)$ can be reindexed by defining $p' \equiv p-1$. This gives

$$\mathbf{R}_p \sum_m \sum_p (m) m^x p^y \mathbf{P}(m, p-1) = \mathbf{R}_p \sum_m \sum_{p'=-1}^{\infty} m^{(x+1)} (p+1)^y \mathbf{P}(m, p'). \tag{S60}$$

We again use the fact that negative molecule copy numbers are assigned with probability zero to begin the sum from 0 rather than -1 and the arbitrary nature of the label $p'$ to write

$$\mathbf{R}_p \sum_m \sum_{p'=0}^{\infty} m^{(x+1)} (p+1)^y \mathbf{P}(m, p') = \mathbf{R}_p \left\langle \mathbf{m}^{(\mathbf{x}+\mathbf{1})} (\mathbf{p}+\mathbf{1})^{\mathbf{y}} \right\rangle. \tag{S61}$$

Finally, we take care of the protein degradation term involving $\mathbf{P}(m, p+1)$. As before, we define $p' = p+1$ and substitute this to obtain

$$\mathbf{\Gamma}_p \sum_m \sum_p (p+1) m^x p^y \mathbf{P}(m, p+1) = \mathbf{\Gamma}_p \sum_m \sum_{p'=1}^{\infty} (p') m^x (p'-1)^y \mathbf{P}(m, p'). \tag{S62}$$

Just as with the mRNA degradation term, having a term $p'$ inside the sum allows us to start the sum over $p'$ from 0 rather than 1. We can therefore write

$$\mathbf{\Gamma}_p \sum_m \sum_{p'=0}^{\infty} (p') m^x (p'-1)^y \mathbf{P}(m, p') = \mathbf{\Gamma}_p \langle \mathbf{m}^{\mathbf{x}} \mathbf{p}(\mathbf{p}-\mathbf{1})^{\mathbf{y}} \rangle. \tag{S63}$$

Putting all these terms together we can write the general moment ODE. This is of the form

$$\begin{aligned} \frac{d\langle \mathbf{m}^{\mathbf{x}} \mathbf{p}^{\mathbf{y}} \rangle}{dt} &= \mathbf{K} \langle \mathbf{m}^{\mathbf{x}} \mathbf{p}^{\mathbf{y}} \rangle \text{ (promoter state transition)} \\ &- \mathbf{R}_m \langle \mathbf{m}^{\mathbf{x}} \mathbf{p}^{\mathbf{y}} \rangle + \mathbf{R}_m \langle (\mathbf{m}+\mathbf{1})^{\mathbf{x}} \mathbf{p}^{\mathbf{y}} \rangle \text{ (mRNA production)} \\ &- \mathbf{\Gamma}_m \left\langle \mathbf{m}^{(\mathbf{x}+\mathbf{1})} \mathbf{p}^{\mathbf{y}} \right\rangle + \mathbf{\Gamma}_m \langle \mathbf{m}(\mathbf{m}-\mathbf{1})^{\mathbf{x}} \mathbf{p}^{\mathbf{y}} \rangle \text{ (mRNA degradation)} \\ &- \mathbf{R}_p \left\langle \mathbf{m}^{(\mathbf{x}+\mathbf{1})} \mathbf{p}^{\mathbf{y}} \right\rangle + \mathbf{R}_p \left\langle \mathbf{m}^{(\mathbf{x}+\mathbf{1})} (\mathbf{p}+\mathbf{1})^{\mathbf{y}} \right\rangle \text{ (protein production)} \\ &- \mathbf{\Gamma}_p \left\langle \mathbf{m}^{\mathbf{x}} \mathbf{p}^{(\mathbf{y}+\mathbf{1})} \right\rangle + \mathbf{\Gamma}_p \langle \mathbf{m}^{\mathbf{x}} \mathbf{p}(\mathbf{p}-\mathbf{1})^{\mathbf{y}} \rangle \text{ (protein degradation).} \end{aligned} \tag{S64}$$

#### S3.2 Moment closure of the simple-repression distribution

A very interesting and useful feature of Eq. S64 is that for a given value of $x$ and $y$ the moment $\langle \mathbf{m^x p^y} \rangle$ is only a function of lower moments. Specifically $\langle \mathbf{m^x p^y} \rangle$ is a function of moments $\left\langle \mathbf{m^{x'} p^{y'}} \right\rangle$ that satisfy two conditions:

$$
\begin{aligned}
&1) y' \leq y, \\
&2) x' + y' \leq x + y.
\end{aligned} \tag{S65}
$$

To prove this we rewrite Eq. S64 as

$$
\begin{aligned}
\frac{d\langle \mathbf{m^x p^y} \rangle}{dt} &= \mathbf{K} \langle \mathbf{m^x p^y} \rangle \\
&+ \mathbf{R}_m \langle \mathbf{p^y} \left[ (\mathbf{m} + \mathbf{1})^{\mathbf{x}} - \mathbf{m^x} \right] \rangle \\
&+ \mathbf{\Gamma}_m \langle \mathbf{m p^y} \left[ (\mathbf{m} - \mathbf{1})^{\mathbf{x}} - \mathbf{m^x} \right] \rangle \\
&+ \mathbf{R}_p \left\langle \mathbf{m^{(x+1)}} \left[ (\mathbf{p} + \mathbf{1})^{\mathbf{y}} - \mathbf{p^y} \right] \right\rangle \\
&+ \mathbf{\Gamma}_p \langle \mathbf{m^x p} \left[ (\mathbf{p} - \mathbf{1})^{\mathbf{y}} - \mathbf{p^y} \right] \rangle,
\end{aligned} \tag{S66}
$$

where the factorization is valid given the linearity of expected values. Now the objective is to find the highest moment for each term once the relevant binomial, such as $(m-1)^x$, is expanded. Take, for example, a simple case in which we want to find the second moment of the mRNA distribution. We then set $x = 2$ and $y = 0$. Eq. S66 then becomes

$$
\begin{aligned}
\frac{d\left\langle \mathbf{m^2 p^0} \right\rangle}{dt} &= \mathbf{K} \left\langle \mathbf{m^2 p^0} \right\rangle \\
&+ \mathbf{R}_m \left\langle \mathbf{p^0} \left[ (\mathbf{m} + \mathbf{1})^{\mathbf{2}} - \mathbf{m^2} \right] \right\rangle \\
&+ \mathbf{\Gamma}_m \left\langle \mathbf{m p^0} \left[ (\mathbf{m} - \mathbf{1})^{\mathbf{2}} - \mathbf{m^2} \right] \right\rangle \\
&+ \mathbf{R}_p \left\langle \mathbf{m^{(2+1)}} \left[ (\mathbf{p} + \mathbf{1})^{\mathbf{0}} - \mathbf{p^0} \right] \right\rangle \\
&+ \mathbf{\Gamma}_p \left\langle \mathbf{m^2 p} \left[ (\mathbf{p} - \mathbf{1})^{\mathbf{0}} - \mathbf{p^0} \right] \right\rangle.
\end{aligned} \tag{S67}
$$

Simplifying this equation gives

$$
\frac{d\left\langle \mathbf{m^2} \right\rangle}{dt} = \mathbf{K} \left\langle \mathbf{m^2} \right\rangle + \mathbf{R}_m \langle [\mathbf{2m} + \mathbf{1}] \rangle + \mathbf{\Gamma}_m \left\langle \left[ -\mathbf{2m^2} + \mathbf{m} \right] \right\rangle. \tag{S68}
$$

Eq. S68 satisfies both of our conditions. Since we set $y$ to be zero, none of the terms depend on any moment that involves the protein number, therefore $y' \leq y$ is satisfied. Also the highest moment in Eq. S68 also satisfies $x' + y' \leq x + y$ since the second moment of mRNA doesn't depend on any moment higher than $\left\langle \mathbf{m^2} \right\rangle$. To demonstrate that this is true for any $x$ and $y$ we now rewrite Eq. S66, making use of the binomial expansion

$$
(z \pm 1)^n = \sum_{k=0}^{n} \binom{n}{k} (\pm 1)^k z^{n-k}. \tag{S69}
$$

Just as before let's look at each term individually. For the mRNA production term we have

$$\mathbf{R}_m \langle \mathbf{p^y} \left[(\mathbf{m}+\mathbf{1})^{\mathbf{x}} - \mathbf{m^x}\right] \rangle = \mathbf{R}_m \left\langle \mathbf{p^y} \left[ \sum_{\mathbf{k=0}}^{\mathbf{x}} \binom{\mathbf{x}}{\mathbf{k}} \mathbf{m^{x-k}} - \mathbf{m^x} \right] \right\rangle. \tag{S70}$$

When $k = 0$, the term inside the sum on the right-hand side cancels with the other $m^x$, so we can simplify to

$$\mathbf{R}_m \langle \mathbf{p^y} \left[(\mathbf{m}+\mathbf{1})^{\mathbf{x}} - \mathbf{m^x}\right] \rangle = \mathbf{R}_m \left\langle \mathbf{p^y} \left[ \sum_{\mathbf{k=1}}^{\mathbf{x}} \binom{\mathbf{x}}{\mathbf{k}} \mathbf{m^{x-k}} \right] \right\rangle. \tag{S71}$$

Once the sum is expanded we can see that the highest moment in this sum is given by $\left\langle \mathbf{m^{(x-1)}p^y} \right\rangle$ which satisfies both of the conditions on Eq. S65.

For the mRNA degradation term we similarly have

$$\mathbf{\Gamma}_m \langle \mathbf{mp^y} \left[(\mathbf{m}-\mathbf{1})^{\mathbf{x}} - \mathbf{m^x}\right] \rangle = \mathbf{\Gamma}_m \left\langle \mathbf{mp^y} \left[ \sum_{\mathbf{k=0}}^{\mathbf{x}} \binom{\mathbf{x}}{\mathbf{k}} (-\mathbf{1})^{\mathbf{k}} \mathbf{m^{x-k}} - \mathbf{m^x} \right] \right\rangle. \tag{S72}$$

Simplifying terms we obtain

$$\mathbf{\Gamma}_m \left\langle \mathbf{mp^y} \left[ \sum_{\mathbf{k=0}}^{\mathbf{x}} \binom{\mathbf{x}}{\mathbf{k}} (-\mathbf{1})^{\mathbf{k}} \mathbf{m^{x-k}} - \mathbf{m^x} \right] \right\rangle = \mathbf{\Gamma}_m \left\langle \mathbf{p^y} \left[ \sum_{\mathbf{k=1}}^{\mathbf{x}} \binom{\mathbf{x}}{\mathbf{k}} (-\mathbf{1})^{\mathbf{k}} \mathbf{m^{x+1-k}} \right] \right\rangle. \tag{S73}$$

The largest moment in this case is $\langle \mathbf{m^x p^y} \rangle$, which again satisfies the conditions on Eq. S65.

The protein production term gives

$$\mathbf{R}_p \left\langle \mathbf{m^{(x+1)}} \left[(\mathbf{p}+\mathbf{1})^{\mathbf{y}} - \mathbf{p^y}\right] \right\rangle = \mathbf{R}_p \left\langle \mathbf{m^{(x+1)}} \left[ \sum_{\mathbf{k=0}}^{\mathbf{y}} \binom{\mathbf{y}}{\mathbf{k}} (-\mathbf{1})^{\mathbf{k}} \mathbf{p^{y-k}} - \mathbf{p^y} \right] \right\rangle. \tag{S74}$$

Upon simplification we obtain

$$\mathbf{R}_p \left\langle \mathbf{m^{(x+1)}} \left[ \sum_{\mathbf{k=0}}^{\mathbf{y}} \binom{\mathbf{y}}{\mathbf{k}} (-\mathbf{1})^{\mathbf{k}} \mathbf{p^{y-k}} - \mathbf{p^y} \right] \right\rangle = \mathbf{R}_p \left\langle \mathbf{m^{(x+1)}} \left[ \sum_{\mathbf{k=1}}^{\mathbf{y}} \binom{\mathbf{y}}{\mathbf{k}} (-\mathbf{1})^{\mathbf{k}} \mathbf{p^{y-k}} \right] \right\rangle. \tag{S75}$$

Here the largest moment is given by $\left\langle \mathbf{m^{x+1}p^{y-1}} \right\rangle$, that again satisfies both of our conditions. For the last term, for protein degradation we have

$$Rp \left\langle \mathbf{m^{(x+1)}} \left[(\mathbf{p}+\mathbf{1})^{\mathbf{y}} - \mathbf{p^y}\right] \right\rangle = Rp \left\langle \mathbf{m^{(x+1)}} \left[ \sum_{\mathbf{k=1}}^{\mathbf{y}} \binom{\mathbf{y}}{\mathbf{k}} (-\mathbf{1^k}) \mathbf{p^{y-k}} \right] \right\rangle. \tag{S76}$$

The largest moment involved in this term is therefore $\left\langle \mathbf{m^x p^{y-1}} \right\rangle$. With this, we show that the four terms involved in our general moment equation depend only on lower moments that satisfy Eq. S65.

As a reminder, what we showed in this section is that the kinetic model introduced in Fig. 2(A) has no moment-closure problem. In other words, moments of the joint mRNA and protein distribution can be computed just from knowledge of lower moments. This allows us to cleanly integrate the moments of the distribution dynamics as cells progress through the cell cycle.

#### S3.3 Computing single promoter steady-state moments

(Note: The Python code used for the calculations presented in this section can be found in the following link as an annotated Jupyter notebook)

As discussed in Section 1.4, one of the main factors contributing to cell-to-cell variability in gene expression is the change in gene copy number during the cell cycle as cells replicate their genome before cell division. Our minimal model accounts for this variability by considering the time trajectory of the distribution moments as given by Eq. S66. These predictions will be contrasted with the predictions from a kinetic model that doesn't account for changes in gene copy number during the cell cycle in Appendix S4.

If we do not account for change in gene copy number during the cell cycle or for the partition of proteins during division, the dynamics of the moments of the distribution described in this section will reach a steady state. In order to compute the steady-state moments of the kinetic model with a single gene across the cell cycle, we use the moment closure property of our master equation. By equating Eq. S66 to zero for a given $\mathbf{x}$ and $\mathbf{y}$, we can solve the resulting linear system and obtain a solution for $\langle \mathbf{m}^{\mathbf{x}} \mathbf{p}^{\mathbf{y}} \rangle$ at steady state as a function of moments $\left\langle \mathbf{m}^{\mathbf{x}'} \mathbf{p}^{\mathbf{y}'} \right\rangle$ that satisfy Eq. S65. Then, by solving for the zero$^{\text{th}}$ moment $\langle \mathbf{m}^{\mathbf{0}} \mathbf{p}^{\mathbf{0}} \rangle$ subject to the constraint that the probability of the promoter being in any state should add up to one, we can substitute back all of the solutions in terms of moments $\left\langle \mathbf{m}^{\mathbf{x}'} \mathbf{p}^{\mathbf{y}'} \right\rangle$ with solutions in terms of the rates shown in Fig. 2. In other words, through an iterative process, we can get at the value of any moment of the distribution. We start by solving for the zero$^{\text{th}}$ moment. Since all higher moments, depend on lower moments we can use the solution of the zero$^{\text{th}}$ moment to compute the first mRNA moment. This solution is then used for higher moments in a hierarchical iterative process.

### S4 Accounting for the variability in gene copy number during the cell cycle

(Note: The Python code used for the calculations presented in this section can be found in the following link as an annotated Jupyter notebook)

When growing in rich media, bacteria can double every $\approx 20$ minutes. With two replication forks each traveling at $\approx 1000$ bp per second, and a genome of $\approx 5$ Mbp for *E. coli* [61], a cell would need $\approx 40$ minutes to replicate its genome. The apparent paradox of growth rates faster than one division per 40 minutes is solved by the fact that cells have multiple replisomes, i.e. molecular machines that replicate the genome running in parallel. Cells can have up to 8 copies of the genome being replicated simultaneously depending on the growth rate [37].

This observation implies that during the cell cycle gene copy number varies. This variation depends on the growth rate and the relative position of the gene with respect to the replication origin, having genes close to the replication origin spending more time with multiple copies compare to genes closer to the replication termination site. This change in gene dosage has a direct effect on the cell-to-cell variability in gene expression [25, 38].

#### S4.1 Numerical integration of moment equations

(Note: The Python code used for the calculations presented in this section can be found in the following link as an annotated Jupyter notebook)

For our specific locus (*galK*) and a doubling time of $\approx 60$ min for our experimental conditions, cells have on average 1.66 copies of the reporter gene during the cell cycle [25]. What this means is that cells spend 60% of the time having one copy of the gene and 40% of the time with two copies. To account for this variability in gene copy number across the cell cycle we numerically integrate the moment equations derived in Appendix S3 for a time $t = [0, t_s]$ with an mRNA production rate $r_m$, where $t_s$ is the time point at which the replication fork reaches our specific locus. For the remaining time before the cell division $t = [t_s, t_d]$ that the cell spends with two promoters, we assume that the only parameter that changes is the mRNA production rate from $r_m$ to $2r_m$. This simplifying assumption ignores potential changes in protein translation rate $r_p$ or changes in the repressor copy number that would be reflected in changes on the repressor on rate $k_{\text{on}}^{(r)}$.

##### S4.1.1 Computing distribution moments after cell division

(Note: The Python code used for the calculations presented in this section can be found in the following link as an annotated Jupyter notebook)

We have already solved a general form for the dynamics of the moments of the distribution, i.e. we wrote differential equations for the moments $\frac{d\langle m^x p^y \rangle}{dt}$. Given that we know all parameters for our model we can simply integrate these equations numerically to compute how the moments of the distribution evolve as cells progress through their cell cycle. Once the cell reaches a time $t_d$ when is going to divide the mRNA and proteins that we are interested in undergo a binomial partitioning between the two daughter cells. In other words, each molecule flips a coin and decides whether to go to either daughter. The question then becomes given that we have a value for the moment $\langle m^x p^y \rangle_{t_d}$ at a time before the cell division, what would the value of this moment be after the cell division takes place $\langle m^x p^y \rangle_{t_o}$?

The probability distribution of mRNA and protein after the cell division $P_{t_o}(m, p)$ must satisfy

$$P_{t_o}(m, p) = \sum_{m'=m}^{\infty} \sum_{p'=p}^{\infty} P(m, p \mid m', p') P_{t_d}(m', p'), \tag{S77}$$

where we are summing over all the possibilities of having $m'$ mRNA and $p'$ proteins before cell division. Note that the sums start at $m$ and $p$; this is because for a cell to have these copy numbers before cell division it is a requirement that the mother cell had at least such copy number since we are not assuming that there is any production at the instantaneous cell division time. Since we assume that the partition of mRNA is independent from the partition of protein, the conditional probability $P(m, p \mid m', p')$ is simply given by a product of two binomial distributions, one for the mRNA and one for the protein, i.e.

$$P(m, p \mid m', p') = \binom{m'}{m} \left(\frac{1}{2}\right)^{m'} \cdot \binom{p'}{p} \left(\frac{1}{2}\right)^{p'}. \tag{S78}$$

Because of these product of binomial probabilities are allowed to extend the sum from Eq. S77 to start at $m' = 0$ and $p' = 0$ as

$$P_{t_o}(m, p) = \sum_{m'=0}^{\infty} \sum_{p'=0}^{\infty} P(m, p \mid m', p') P_{t_d}(m', p'), \tag{S79}$$

since the product of the binomial distributions in Eq. S78 is zero for all $m' < m$ and/or $p' < 0$. So from now on in this section we will assume that a sum of the form $\sum_x \equiv \sum_{x=0}^{\infty}$ to simplify notation.

We can then compute the distribution moments after the cell division $\langle m^x p^y \rangle_{t_o}$ as

$$\langle m^x p^y \rangle_{t_o} = \sum_m \sum_p m^x p^y P_{t_o}(m, p), \tag{S80}$$

for all $x, y \in \mathbb{N}$. Substituting Eq. S77 results in

$$\langle m^x p^y \rangle_{t_o} = \sum_m \sum_p m^x p^y \sum_{m'} \sum_{p'} P(m, p \mid m', p') P_{t_d}(m', p'). \tag{S81}$$

We can rearrange the sums to be

$$\langle m^x p^y \rangle_{t_o} = \sum_{m'} \sum_{p'} P_{t_d}(m', p') \sum_m \sum_p m^x p^y P(m, p \mid m', p'). \tag{S82}$$

The fact that Eq. S78 is the product of two independent events allows us to rewrite the joint probability $P(m, p \mid m', p')$ as

$$P(m, p \mid m', p') = P(m \mid m') \cdot P(p \mid p'). \tag{S83}$$

With this we can then write the moment $\langle m^x p^y \rangle_{t_o}$ as

$$\langle m^x p^y \rangle_{t_o} = \sum_{m'} \sum_{p'} P_{t_d}(m', p') \sum_m m^x P(m \mid m') \sum_p p^y P(p \mid p'). \tag{S84}$$

Notice that both terms summing over $m$ and over $p$ are the conditional expected values, i.e.

$$\sum_z z^x P(z \mid z') \equiv \left\langle z^x \mid z' \right\rangle, \quad \text{for } z \in \{m, p\}. \tag{S85}$$

These conditional expected values are the expected values of a binomial random variable $z \sim \text{Bin}(z', 1/2)$, which can be easily computed as we will show later in this section. We then rewrite the expected values after the cell division in terms of these moments of a binomial distribution

$$\langle m^x p^y \rangle_{t_o} = \sum_{m'} \sum_{p'} \left\langle m^x \mid m' \right\rangle \left\langle p^y \mid p' \right\rangle P_{t_d}(m', p'). \tag{S86}$$

To see how this general formula for the moments after the cell division works let's compute the mean protein per cell after the cell division $\langle p \rangle_{t_o}$. That is setting $x = 0$, and $y = 1$. This results in

$$\langle p \rangle_{t_o} = \sum_{m'} \sum_{p'} \left\langle m^0 \mid m' \right\rangle \left\langle p \mid p' \right\rangle P_{t_d}(m', p'). \tag{S87}$$

The zeroth moment $\left\langle m^0 \mid m' \right\rangle$ by definition must be one since we have

$$\left\langle m^0 \mid m' \right\rangle = \sum_m m^0 P(m \mid m') = \sum_m P(m \mid m') = 1, \tag{S88}$$

since the probability distribution must be normalized. This leaves us then with

$$\langle p \rangle_{t_o} = \sum_{m'} \sum_{p'} P_{t_d}(m', p') \left\langle p \mid p' \right\rangle. \tag{S89}$$

If we take the sum over $m'$ we simply compute the marginal probability distribution $\sum_{m'} P_{t_d}(m', p') = P_{t_d}(p')$, then we have

$$\langle p \rangle_{t_o} = \sum_{p'} \left\langle p \mid p' \right\rangle P_{t_d}(p'). \tag{S90}$$

For the particular case of the first moment of the binomial distribution with parameters $p'$ and $1/2$ we know that

$$\left\langle p \mid p' \right\rangle = \frac{p'}{2}. \tag{S91}$$

Therefore the moment after division is equal to

$$\langle p \rangle_{t_o} = \sum_{p'} \frac{p'}{2} P_{t_d}(p') = \frac{1}{2} \sum_{p'} p' P_{t_d}(p'). \tag{S92}$$

Notice that this is just $1/2$ of the expected value of $p'$ averaging over the distribution prior to cell division, i.e.

$$\langle p \rangle_{t_o} = \frac{\langle p' \rangle_{t_d}}{2}, \tag{S93}$$

where $\langle \cdot \rangle_{t_d}$ highlights that is the moment of the distribution prior to the cell division. This result makes perfect sense. What this is saying is that the mean protein copy number right after the cell divides is half of the mean protein copy number just before the cell division. That is exactly we would expect. So in principle to know the first moment of either the mRNA distribution $\langle m \rangle_{t_o}$ or the protein distribution $\langle m \rangle_{t_o}$ right after cell division it suffices to multiply the moments before the cell division $\langle m \rangle_{t_d}$ or $\langle p \rangle_{t_d}$ by $1/2$. Let's now explore how this generalizes to any other moment $\langle m^x p^y \rangle_{t_o}$.

##### S4.1.2 Computing the moments of a binomial distribution

The result from last section was dependent on us knowing the functional form of the first moment of the binomial distribution. For higher moments we need some systematic way to compute such moments. Luckily for us we can do so by using the so-called moment generating function (MGF). The MGF of a random variable $X$ is defined as

$$M_X(t) = \left\langle e^{tX} \right\rangle, \tag{S94}$$

where $t$ is a dummy variable. Once we know the MGF we can obtain any moment of the distribution by simply computing

$$\langle X^n \rangle = \left. \frac{d^n}{dt^n} M_X(t) \right|_{t=0}, \tag{S95}$$

i.e. taking the $n$-th derivative of the MGF returns the $n$-th moment of the distribution. For the particular case of the binomial distribution $X \sim \text{Bin}(N, q)$ it can be shown that the MGF is of the form

$$M_X(t) = \left[(1-q) + qe^t\right]^N. \tag{S96}$$

As an example let's compute the first moment of this binomially distributed variable. For this, the first derivative of the MGF results in

$$\frac{dM_X(t)}{dt} = N[(1-q) + qe^t]^{N-1} qe^t. \tag{S97}$$

We just need to follow Eq. S95 and set $t = 0$ to obtain the first moment

$$\left.\frac{dM_X(t)}{dt}\right|_{t=0} = Nq, \tag{S98}$$

which is exactly the expected value of a binomially distributed random variable.

So according to Eq. S86 to compute any moment $\langle m^x p^y \rangle$ after cell division we can just take the $x$-th derivative and the $y$-th derivative of the binomial MGF to obtain $\langle m^x \mid m' \rangle$ and $\langle p^y \mid p' \rangle$, respectively, and take the expected value of the result. Let's follow on detail the specific case for the moment $\langle mp \rangle$. When computing the moment after cell division $\langle mp \rangle_{t_o}$ which is of the form

$$\langle mp \rangle_{t_o} = \sum_{m'} \sum p' \left\langle m \mid m' \right\rangle \left\langle p \mid p' \right\rangle P_{t_d}(m', p'), \tag{S99}$$

the product $\langle m \mid m' \rangle \langle p \mid p' \rangle$ is then

$$\left\langle m \mid m' \right\rangle \left\langle p \mid p' \right\rangle = \frac{m'}{2} \cdot \frac{p'}{2}, \tag{S100}$$

where we used the result in Eq. S98, substituting $m$ and $p$ for $X$, respectively, and $q$ for $1/2$. Substituting this result into the moment gives

$$\langle mp \rangle_{t_o} = \sum_{m'} \sum_{p'} \frac{m'p'}{4} P_{t_d}(m', p') = \frac{\langle m'p' \rangle_{t_d}}{4}. \tag{S101}$$

Therefore to compute the moment after cell division $\langle mp \rangle_{t_o}$ we simply have to divide by 4 the corresponding equivalent moment before the cell division.

Not all moments after cell division depend only on the equivalent moment before cell division. For example if we compute the third moment of the protein distribution $\left\langle p^3 \right\rangle_{t_o}$, we find

$$\left\langle p^3 \right\rangle_{t_o} = \frac{\left\langle p^3 \right\rangle_{t_d}}{8} + \frac{3 \left\langle p^2 \right\rangle_{t_d}}{8}. \tag{S102}$$

So for this particular case the third moment of the protein distribution depends on the third moment and the second moment before the cell division. In general all moments after cell division $\langle m^x p^y \rangle_{t_o}$ linearly depend on moments before cell division. Furthermore, there is "moment closure" for this specific case in the sense that all moments after cell division depend on lower moments before cell division. To generalize these results to all the moments computed in this work let us then define a vector to collect all moments before the cell division up the $\langle m^x p^y \rangle_{t_d}$ moment, i.e.

$$\langle \mathbf{m^x p^y} \rangle_{t_d} = \left( \left\langle m^0 p^0 \right\rangle_{t_d}, \left\langle m^1 \right\rangle_{t_d}, \ldots, \langle m^x p^y \rangle_{t_d} \right). \tag{S103}$$

Then any moment after cell division $\left\langle m^{x'} p^{y'} \right\rangle_{t_o}$ for $x' \leq x$ and $y' \leq y$ can be computed as

$$\left\langle m^{x'} p^{y'} \right\rangle_{t_o} = \mathbf{z}_{x'y'} \cdot \langle \mathbf{m^x p^y} \rangle_{t_d},$$

where we define the vector $\mathbf{z}_{x'y'}$ as the vector containing all the coefficients that we obtain with the product of the two binomial distributions. For example for the case of the third protein moment $\left\langle p^3 \right\rangle_{t_o}$ the vector $\mathbf{z}_{x'y'}$ would have zeros for all entries except for the corresponding entry for $\left\langle p^2 \right\rangle_{t_d}$ and for $\left\langle p^3 \right\rangle_{t_d}$, where it would have 3/8 and 1/8 accordingly.

If we want then to compute all the moments after the cell division up to $\langle m^x p^y \rangle_{t_o}$ let us define an equivalent vector

$$\langle \mathbf{m^x p^y} \rangle_{t_o} = \left( \langle m^0 p^0 \rangle_{t_o}, \langle m^1 \rangle_{t_o}, \ldots, \langle m^x p^y \rangle_{t_o} \right). \tag{S104}$$

Then we need to build a square matrix $\mathbf{Z}$ such that each row of the matrix contains the corresponding vector $\mathbf{z}_{x'y'}$ for each of the moments. Having this matrix we would simply compute the moments after the cell division as

$$\langle \mathbf{m^x p^x} \rangle_{t_o} = \mathbf{Z} \cdot \langle \mathbf{m^x p^x} \rangle_{t_d}. \tag{S105}$$

In other words, matrix $\mathbf{Z}$ will contain all the coefficients that we need to multiply by the moments before the cell division in order to obtain the moments after cell division. Matrix $\mathbf{Z}$ was then generated automatically using Python's analytical math library sympy [62].

Fig. S8 (adapted from Fig. 3(B)) shows how the first moment of both mRNA and protein changes over several cell cycles. The mRNA quickly relaxes to the steady state corresponding to the parameters for both a single and two promoter copies. This is expected since the parameters for the mRNA production were determined in the first place under this assumption (See Appendix S1). We note that there is no apparent delay before reaching steady state of the mean mRNA count after the cell divides. This is because the mean mRNA count for the two promoters copies state is exactly twice the expected mRNA count for the single promoter state (See Appendix S1). Therefore once the mean mRNA count is halved after the cell division, it is already at the steady state value for the single promoter case. On the other hand, given that the relaxation time to steady state is determined by the degradation rate, the mean protein count does not reach its corresponding steady state value for either promoter copy number state. Interestingly once a couple of cell cycles have passed the first moment has a repetitive trajectory over cell cycles. We have observed this experimentally by tracking cells as they grow under the microscope. Comparing cells at the beginning of the cell cycle with the daughter cells that appear after cell division shown that on average all cells have the same amount of protein at the beginning of the cell cycle (See Fig. 18 of [28]), suggesting that these dynamical steady state takes place *in vivo*.

In principle when measuring gene expression levels experimentally from an asynchronous culture, cells are sampled from any time point across their individual cell cycles. This means that the moments determined experimentally correspond to an average over the cell cycle. In the following section we discuss how to account for the fact that cells are not uniformly distributed across the cell cycle in order to compute these averages.

#### S4.2 Exponentially distributed ages

As mentioned in Appendix S2, cells in exponential growth have exponentially distributed ages across the cell cycle, having more young cells compared to old ones. Specifically the probability of a

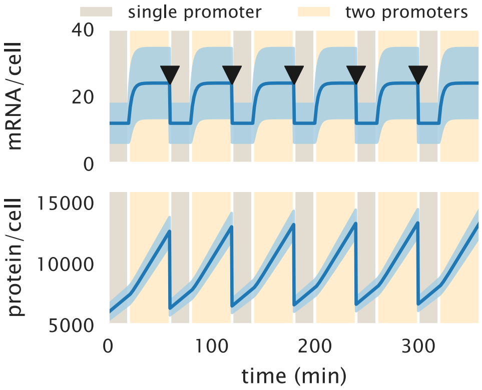

**Figure S8. First and second moment dynamics over cell the cell cycle.** Mean ± standard deviation mRNA (upper panel) and mean ± standard deviation protein copy number (lower panel) as the cell cycle progresses. The dark shaded region delimits the fraction of the cell cycle that cells spend with a single copy of the promoter. The light shaded region delimits the fraction of the cell cycle that cells spend with two copies of the promoter. For a 100 min doubling time at the *galK* locus cells spend 60% of the time with one copy of the promoter and the rest with two copies.

cell being at any time point in the cell cycle is given by [40]

$$P(a) = (\ln 2) \cdot 2^{1-a}, \tag{S106}$$

where $a \in [0, 1]$ is the stage of the cell cycle, with $a = 0$ being the start of the cycle and $a = 1$ being the cell division. In Appendix S9 we reproduce this derivation. It is a surprising result, but can be intuitively thought as follows: If the culture is growing exponentially, that means that all the time there is an increasing number of cells. That means for example that if in a time interval $\Delta t$ $N$ "old" cells divided, these produced $2N$ "young" cells. So at any point there is always more younger than older cells.

Our numerical integration of the moment equations gave us a time evolution of the moments as cells progress through the cell cycle. Since experimentally we sample asynchronous cells that follow Eq. S106, each time point along the moment dynamic must be weighted by the probability of having sampled a cell at such specific time point of the cell cycle. Without loss of generality let's focus on the first mRNA moment $\langle m(t) \rangle$ (the same can be applied to all other moments). As mentioned before, in order to calculate the first moment across the entire cell cycle we must weigh each time point by the corresponding probability that a cell is found in such point of its cell cycle. This translates to computing the integral

$$\langle m \rangle_c = \int_{\text{beginning cell cycle}}^{\text{end cell cycle}} \langle m(t) \rangle \, P(t) dt, \tag{S107}$$

where $\langle m \rangle_c$ is the mean mRNA copy number averaged over the entire cell cycle trajectory, and $P(t)$ is the probability of a cell being at a time $t$ of its cell cycle.

If we set the time in units of the cell cycle length we can use Eq. S106 and compute instead

$$\langle m \rangle = \int_0^1 \langle m(a) \rangle \, P(a) da, \tag{S108}$$

where $P(a)$ is given by Eq. S106.

What Eq. S108 implies is that in order to compute the first moment (or any moment of the distribution) we must weigh each point in the moment dynamics by the corresponding probability of a cell being at that point along its cell cycle. That is why when computing a moment we take the time trajectory of a single cell cycle as the ones shown in Fig. S8 and compute the average using Eq. S106 to weigh each time point. We perform this integral numerically for all moments using Simpson's rule.

#### S4.3 Reproducing the equilibrium picture

Given the large variability of the first moments depicted in Fig. S8 it is worth considering why a simplistic equilibrium picture has shown to be very successful in predicting the mean expression level under diverse conditions [20, 24, 26, 27]. In this section we compare the simple repression thermodynamic model with this dynamical picture of the cell cycle. But before diving into this comparison, it is worth recapping the assumptions that go into the equilibrium model.

##### S4.3.1 Steady state under the thermodynamic model

Given the construction of the thermodynamic model of gene regulation for which the probability of the promoter microstates rather than the probability of mRNA or protein counts is accounted for, we are only allowed to describe the dynamics of the first moment using this theoretical framework [36]. Again let's only focus on the mRNA first moment $\langle m \rangle$. The same principles apply if we consider the protein first moment. We can write a dynamical system of the form

$$\frac{d\langle m \rangle}{dt} = r_m \cdot p_{\text{bound}} - \gamma_m \langle m \rangle, \tag{S109}$$

where as before $r_m$ and $\gamma_m$ are the mRNA production and degradation rates respectively, and $p_{\text{bound}}$ is the probability of finding the RNAP bound to the promoter [54]. This dynamical system is predicted to have a single stable fixed point that we can find by computing the steady state. When we solve for the mean mRNA copy number at steady state $\langle m \rangle_{ss}$ we find

$$\langle m \rangle_{ss} = \frac{r_m}{\gamma_m} p_{\text{bound}}. \tag{S110}$$

Since we assume that the only effect that the repressor has over the regulation of the promoter is exclusion of the RNAP from binding to the promoter, we assume that only $p_{\text{bound}}$ depends on the repressor copy number $R$. Therefore when computing the fold-change in gene expression we are left with

$$\text{fold-change} = \frac{\langle m(R \neq 0) \rangle_{ss}}{\langle m(R = 0) \rangle_{ss}} = \frac{p_{\text{bound}}(R \neq 0)}{p_{\text{bound}}(R = 0)}. \tag{S111}$$

As derived in [20] this can be written in the language of equilibrium statistical mechanics as

$$\text{fold-change} = \left(1 + \frac{R}{N_{\text{NS}}} e^{-\beta \Delta\varepsilon_r}\right)^{-1}, \tag{S112}$$

where $\beta \equiv (k_B T)^{-1}$, $\Delta\varepsilon_r$ is the repressor-DNA binding energy, and $N_{\text{NS}}$ is the number of non-specific binding sites where the repressor can bind.

To arrive at Eq. S112 we ignore the physiological changes that occur during the cell cycle; one of the most important being the variability in gene copy number that we are exploring in this section.

It is therefore worth thinking about whether or not the dynamical picture exemplified in Fig. S8 can be reconciled with the predictions made by Eq. S112 both at the mRNA and protein level.

Fig. S9 compares the predictions of both theoretical frameworks for varying repressor copy numbers and repressor-DNA affinities. The solid lines are directly computed from Eq. S112. The hollow triangles and the solid circles, represent the fold-change in mRNA and protein respectively as computed from the moment dynamics. To compute the fold-change from the kinetic picture we first numerically integrate the moment dynamics for both the two- and the three-state promoter (See Fig. S8 for the unregulated case) and then average the time series accounting for the probability of cells being sampled at each stage of the cell cycle as defined in Eq. S108. The small systematic deviations between both models come partly from the simplifying assumption that the repressor copy number, and therefore the repressor on rate $k_{\text{on}}^{(r)}$ remains constant during the cell cycle. In principle the gene producing the repressor protein itself is also subjected to the same duplication during the cell cycle, changing therefore the mean repressor copy number for both stages.

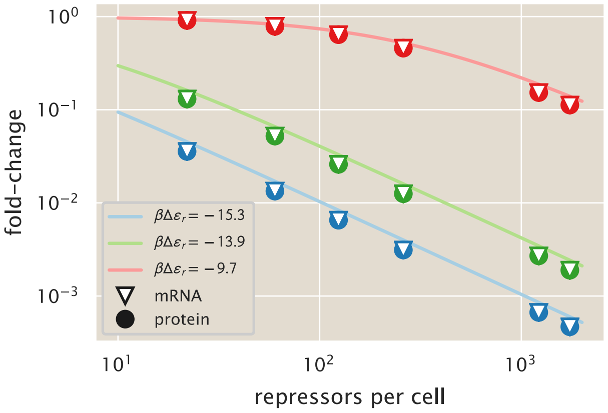

**Figure S9. Comparison of the equilibrium and kinetic reressor titration predictions.** The equilibrium model (solid lines) and the kinetic model with variation over the cell cycle (solid circles and white triangles) predictions are compared for varying repressor copy numbers and operator binding energy. The equilibrium model is directly computed from Eq. S112 while the kinetic model is computed by numerically integrating the moment equations over several cell cycles, and then averaging over the extent of the cell cycle as defined in Eq. S108.

For completeness Fig. S10 compares the kinetic and equilibrium models for the extended model of [27] in which the inducer concentration enters into the equation. The solid line is directly computed from Eq. 5 of [27]. The hollow triangles and solid points follow the same procedure as for Fig. S9, where the only effect that the inducer is assume to have in the kinetics is an effective change in the number of active repressors, affecting therefore $k_{\text{on}}^{(r)}$.

#### S4.4 Comparison between single- and multi-promoter kinetic model

After these calculations it is worth questioning whether the inclusion of this change in gene dosage is drastically different with respect to the simpler picture of a kinetic model that ignores the gene copy number variability during the cell cycle. To this end we systematically computed the average moments for varying repressor copy number and repressor-DNA affinities. We then compare these results with the moments obtained from a single-promoter model and their corresponding parameters.

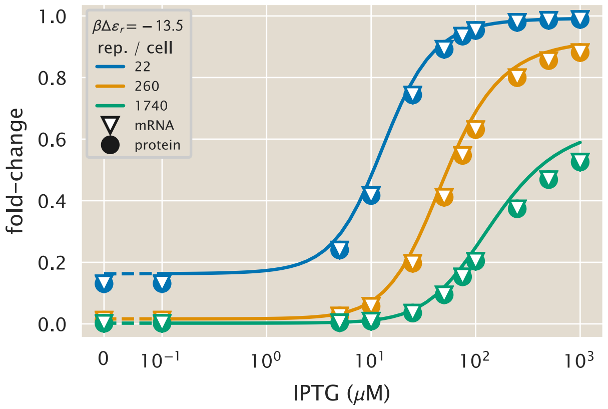

**Figure S10. Comparison of the equilibrium and kinetic inducer titration predictions.** The equilibrium model (solid lines) and the kinetic model with variation over the cell cycle (solid circles and white triangles) predictions are compared for varying repressor copy numbers and inducer concentrations. The equilibrium model is directly computed as Eq. 5 of reference [27] with repressor-DNA binding energy $\Delta\varepsilon_r = -13.5\ k_BT$ while the kinetic model is computed by numerically integrating the moment dynamics over several cell cycles, and then averaging over the extent of a single cell cycle as defined in Eq. S108.

The derivation of the steady-state moments of the distribution for the single-promoter model are detailed in Appendix S3.

Fig. S9 and Fig. S10 both suggest that since the dynamic multi-promoter model can reproduce the results of the equilibrium model at the first moment level it must then also be able to reproduce the results of the single-promoter model at this level (See Appendix S2). The interesting comparison comes with higher moments. A useful metric to consider for gene expression variability is the noise in gene expression [53]. This quantity, defined as the standard deviation divided by the mean, is a dimensionless metric of how much variability there is with respect to the mean of a distribution. As we will show below this quantity differs from the also commonly used metric known as the Fano factor (variance / mean) in the sense that for experimentally determined expression levels in fluorescent arbitrary units, the noise is a dimensionless quantity while the Fano factor is not.

Fig. S11 shows the comparison of the predicted protein noise between the single- (dashed lines) and the multi-promoter model (solid lines) for different operators and repressor copy numbers. A striking difference between both is that the single-promoter model predicts that as the inducer concentration increases, the standard deviation grows much slower than the mean, giving a very small noise. In comparison the multi-promoter model has a much higher floor for the lowest value of the noise, reflecting the expected result that the variability in gene copy number across the cell cycle should increase the cell-to-cell variability in gene expression [25, 38]

#### S4.5 Comparison with experimental data

Having shown that the kinetic model presented in this section can not only reproduce the results from the equilibrium picture at the mean level (See Fig. S9 and Fig. S10), but make predictions for the cell-to-cell variability as quantified by the noise (See Fig. S11), we can assess whether or not this model is able to predict experimental measurements of the noise. For this we take the single cell intensity measurements (See Methods) to compute the noise at the protein level.

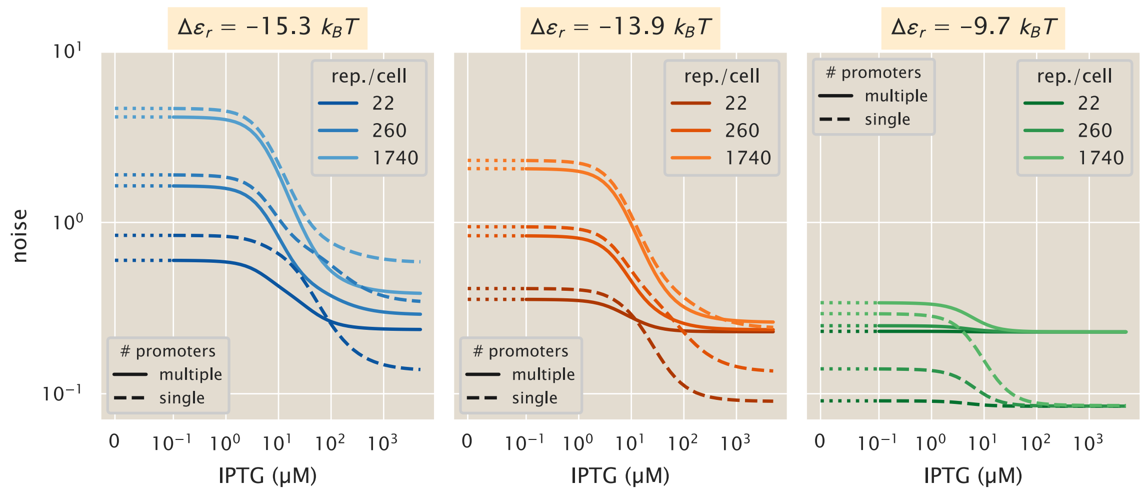

**Figure S11. Comparison of the predicted protein noise between a single- and a multi-promoter kinetic model.** Comparison of the noise (standard deviation/mean) between a kinetic model that considers a single promoter at all times (dashed line) and the multi-promoter model developed in this section (solid line) for different repressor operators. (A) Operator O1, $\Delta\varepsilon_r = -15.3\ k_BT$, (B) O2, $\Delta\varepsilon_r = -13.9\ k_BT$, (C) O3, $\Delta\varepsilon_r = -9.7\ k_BT$

As mentioned before this metric differs from the Fano factor since for fluorescent arbitrary units the noise is a dimensionless quantity. To see why consider that the noise is defined as

$$\text{noise} \equiv \frac{\sqrt{\langle p^2 \rangle - \langle p \rangle^2}}{\langle p \rangle}. \tag{S113}$$

We assume that the intensity level of a cell $I$ is linearly proportional to the absolute protein count, i.e.

$$I = \alpha p, \tag{S114}$$

where $\alpha$ is the proportionality constant between arbitrary units and protein absolute number $p$. Substituting this definition on Eq. S113 gives

$$\text{noise} = \frac{\sqrt{\langle (\alpha I)^2 \rangle - \langle \alpha I \rangle^2}}{\langle \alpha I \rangle}. \tag{S115}$$

Since $\alpha$ is a constant it can be taken out of the average operator $\langle \cdot \rangle$, obtaining

$$\text{noise} = \frac{\sqrt{\alpha^2 \left( \langle I^2 \rangle - \langle I \rangle^2 \right)}}{\alpha \langle I \rangle} = \frac{\sqrt{\left( \langle I^2 \rangle - \langle I \rangle^2 \right)}}{\langle I \rangle}. \tag{S116}$$

Notice that in Eq. S114 the linear proportionality between intensity and protein count has no intercept. This ignores the autofluorescence that cells without reporter would generate. To account for this, in practice we compute

$$\text{noise} = \frac{\sqrt{\left( \langle (I - \langle I_{\text{auto}} \rangle)^2 \rangle - \langle I - \langle I_{\text{auto}} \rangle \rangle^2 \right)}}{\langle I - \langle I_{\text{auto}} \rangle \rangle}. \tag{S117}$$

where $I$ is the intensity of the strain of interest and $\langle I_{\text{auto}} \rangle$ is the mean autofluorescence intensity, obtained from a strain that does not carry the fluorescent reporter gene.

Fig. S12 shows the comparison between theoretical predictions and experimental measurements for the unregulated promoter. The reason we split the data by operator despite the fact that since these are unregulated promoters, they should in principle have identical expression profiles is to precisely make sure that this is the case. We have found in the past that sequences downstream of the RNAP binding site can affect the expression level of constitutively expressed genes. We can see that both models, the single-promoter (gray dotted line) and the multi-promoter (black dashed line) underestimate the experimental noise to different degrees. The single-promoter model does a worse job at predicting the experimental data since it doesn't account for the differences in gene dosage during the cell cycle. But still we can see that accounting for this variability takes us to within a factor of two of the experimentally determined noise for these unregulated strains.

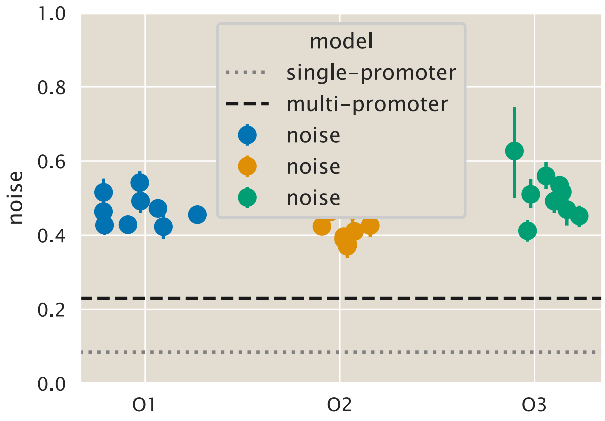

**Figure S12. Protein noise of the unregulated promoter.** Comparison of the experimental noise for different operators with the theoretical predictions for the single-promoter (gray dotted line) and the multi-promoter model (black dashed line). Each datum represents a single date measurement of the corresponding $\Delta lacI$ strain with $\geq 300$ cells. The points correspond to the median, and the error bars correspond to the 95% confidence interval as determined by 10,000 bootstrap samples.

To further test the model predictive power we compare the predictions for the three-state regulated promoter. Fig. S13 shows the theoretical predictions for the single- and multi-promoter model for varying repressor copy numbers and repressor-DNA binding affinities as a function of the inducer concentration. We can see again that our zero-parameter fits systematically underestimates the noise for all strains and all inducer concentrations. We highlight that the $y$-axis is shown in a log-scale to emphasize more this deviation; but, as we will show in the next section, our predictions still fall within a factor of two from the experimental data.

##### S4.5.1 Systematic deviation of the noise in gene expression

Fig. S12 and Fig. S13 highlight that our model underestimates the cell-to-cell variability as measured by the noise. To further explore this systematic deviation Fig. S14 shows the theoretical vs. experimental noise both in linear and log scale. As we can see the data is systematically above the identity line. The data is colored by their corresponding experimental fold-change values. The data that has the largest deviations from the identity line also corresponds to the data with the largest

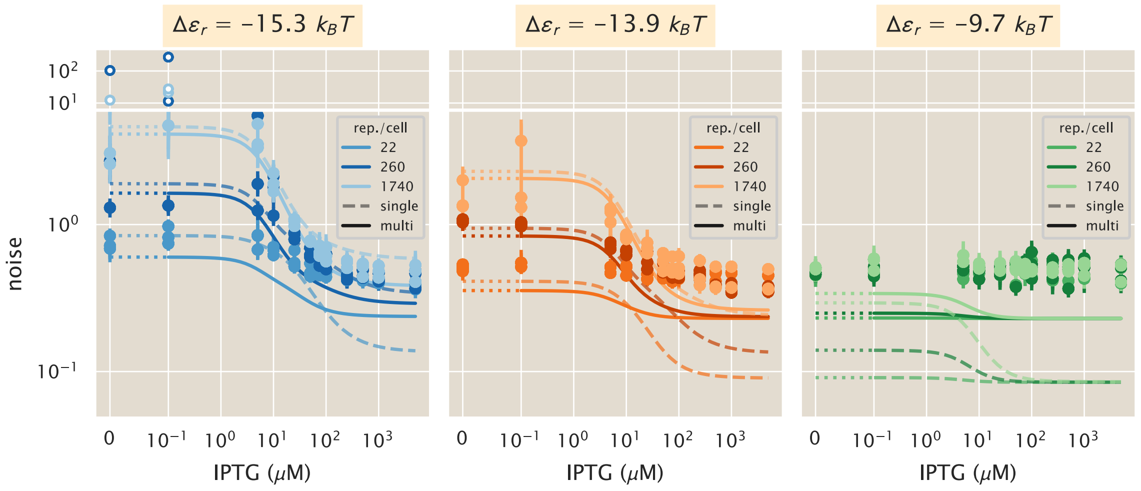

**Figure S13. Protein noise of the regulated promoter.** Comparison of the experimental noise for different operators ((A) O1, $\Delta\varepsilon_r = -15.3\ k_BT$, (B) O2, $\Delta\varepsilon_r = -13.9\ k_BT$, (C) O3, $\Delta\varepsilon_r = -9.7\ k_BT$) with the theoretical predictions for the single-promoter (dashed lines) and the multi-promoter model (solid lines). Points represent the experimental noise as computed from single-cell fluorescence measurements of different *E. coli* strains under 12 different inducer concentrations. Dotted line indicates plot in linear rather than logarithmic scale. Each datum represents a single date measurement of the corresponding strain and IPTG concentration with $\geq 300$ cells. The points correspond to the median, and the error bars correspond to the 95% confidence interval as determined by 10,000 bootstrap samples. White-filled dots are plot at a different scale for better visualization.

error bars and the smallest fold-change. This is because measurements with very small fold-changes correspond to intensities very close to the autofluorescence background. Therefore minimal changes when computing the noise are amplified given the ratio of std/mean. In Appendix S8 we will explore empirical ways to improve the agreement between our minimal model and the experimental data to guide future efforts to improve the minimal.

### S5 Maximum entropy approximation of distributions

(Note: The Python code used for the calculations presented in this section can be found in the following link as an annotated Jupyter notebook)

On the one hand the solution of chemical master equations like the one in Section 1.1 represent a hard mathematical challenge. As presented in Appendix S2 Peccoud and Ycart derived a closed-form solution for the two-state promoter at the mRNA level [34]. In an impressive display of mathematical skills, Shahrezaei and Swain were able to derive an approximate solution for the one- (not considered in this work) and two-state promoter master equation at the protein level [53]. Nevertheless both of these solutions do not give instantaneous insights about the distributions as they involve complicated terms such as confluent hypergeometric functions.

On the other hand there has been a great deal of work to generate methods that can approximate the solution of these discrete state Markovian models [63–67]. In particular for master equations like the one that concerns us here whose moments can be easily computed, the moment expansion method provides a simple method to approximate the full joint distribution of mRNA and protein [67]. In

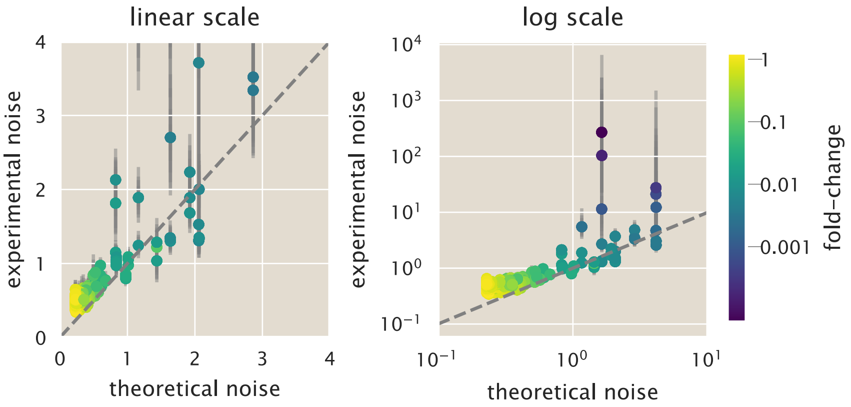

**Figure S14. Systematic comparison of theoretical vs experimental noise in gene expression.** Theoretical vs. experimental noise both in linear (left) and log (right) scale. The dashed line shows the identity line of slope 1 and intercept zero. All data are colored by the corresponding value of the experimental fold-change in gene expression as indicated by the color bar. Each datum represents a single date measurement of the corresponding strain and IPTG concentration with $\geq 300$ cells. The points correspond to the median, and the error bars correspond to the 95% confidence interval as determined by 10,000 bootstrap samples.

this section we will explain the principles behind this method and show the implementation for our particular case study.

#### S5.1 The MaxEnt principle

The principle of maximum entropy (MaxEnt) first proposed by E. T. Jaynes in 1957 tackles the question of given limited information what is the least biased inference one can make about a particular probability distribution [42]. In particular Jaynes used this principle to show the correspondence between statistical mechanics and information theory, demonstrating, for example, that the Boltzmann distribution is the probability distribution that maximizes Shannon's entropy subject to a constraint that the average energy of the system is fixed.

To illustrate the principle let us focus on a univariate distribution $P_X(x)$. The $n^{\text{th}}$ moment of the distribution for a discrete set of possible values of $x$ is given by

$$\langle x^n \rangle \equiv \sum_x x^n P_X(x). \tag{S118}$$

Now assume that we have knowledge of the first $m$ moments $\langle \mathbf{x} \rangle_m = (\langle x \rangle, \langle x^2 \rangle, \ldots, \langle x^m \rangle)$. The question is then how can we use this information to build an estimator $P_H(x \mid \langle \mathbf{x} \rangle_m)$ of the distribution such that

$$\lim_{m \to \infty} P_H(x \mid \langle \mathbf{x} \rangle_m) \to P_X(x), \tag{S119}$$

i.e. that the more moments we add to our approximation, the more the estimator distribution converges to the real distribution.

The MaxEnt principle tells us that our best guess for this estimator is to build it on the base of maximizing the Shannon entropy, constrained by the information we have about these $m$ moments.

The maximization of Shannon's entropy guarantees that we are the least committed possible to information that we do not posses. The Shannon entropy for an univariate discrete distribution is given by [3]

$$H(x) \equiv -\sum_x P_X(x) \log P_X(x). \tag{S120}$$

For an optimization problem subject to constraints we make use of the method of the Lagrange multipliers. For this we define the constraint equation $\mathcal{L}(x)$ as

$$\mathcal{L}(x) \equiv H(x) - \sum_{i=0}^{m} \left[ \lambda_i \left( \left\langle x^i \right\rangle - \sum_x x^i P_X(x) \right) \right], \tag{S121}$$

where $\lambda_i$ is the Lagrange multiplier associated with the $i^{\text{th}}$ moment. The inclusion of the zeroth moment is an additional constraint to guarantee the normalization of the resulting distribution. Since $P_X(x)$ has a finite set of discrete values, when taking the derivative of the constraint equation with respect to $P_X(x)$, we chose a particular value of $X = x$. Therefore from the sum over all possible $x$ values only a single term survives. With this in mind we take the derivative of the constraint equation obtaining

$$\frac{d\mathcal{L}}{dP_X(x)} = -\log P_X(x) - 1 - \sum_{i=0}^{m} \lambda_i x^i. \tag{S122}$$

Equating this derivative to zero and solving for the distribution (that we now start calling $P_H(x)$, our MaxEnt estimator) gives

$$P_H(x) = \exp\left( -1 - \sum_{i=0}^{m} \lambda_i x^i \right) = \frac{1}{\mathcal{Z}} \exp\left( -\sum_{i=1}^{m} \lambda_i x^i \right), \tag{S123}$$

where $\mathcal{Z}$ is the normalization constant that can be obtained by substituting this solution into the normalization constraint. This results in

$$\mathcal{Z} \equiv \exp\left(1 + \lambda_0\right) = \sum_x \exp\left( -\sum_{i=1}^{m} \lambda_i x^i \right). \tag{S124}$$

Eq. S123 is the general form of the MaxEnt distribution for a univariate distribution. The computational challenge then consists in finding numerical values for the Lagrange multipliers $\{\lambda_i\}$ such that $P_H(x)$ satisfies our constraints. In other words, the Lagrange multipliers weight the contribution of each term in the exponent such that when computing any of the moments we recover the value of our constraint. Mathematically what this means is that $P_H(x)$ must satisfy

$$\sum_x x^n P_H(x) = \sum_x \frac{x^n}{\mathcal{Z}} \exp\left( -\sum_{i=1}^{m} \lambda_i x^i \right) = \langle x^n \rangle. \tag{S125}$$

As an example of how to apply the MaxEnt principle let us use the classic problem of a six-face die. If we are only told that after a large number of die rolls the mean value of the face is $\langle x \rangle = 4.5$

(note that a fair die has a mean of 3.5), what would the least biased guess for the distribution look like? The MaxEnt principle tells us that our best guess would be of the form

$$P_H(x) = \frac{1}{\mathcal{Z}} \exp\left(\lambda x\right). \tag{S126}$$

Using any numerical minimization package we can easily find the value of the Lagrange multiplier $\lambda$ that satisfies our constraint. Fig. S15 shows two two examples of distributions that satisfy the constraint. Panel (A) shows a distribution consistent with the 4.5 average where both 4 and 5 are equally likely. Nevertheless in the information we got about the nature of the die it was never stated that some of the faces were forbidden. In that sense the distribution is committing to information about the process that we do not posses. Panel (B) by contrast shows the MaxEnt distribution that satisfies this constraint. Since this distribution maximizes Shannon's entropy it is guaranteed to be the least biased distribution given the available information.

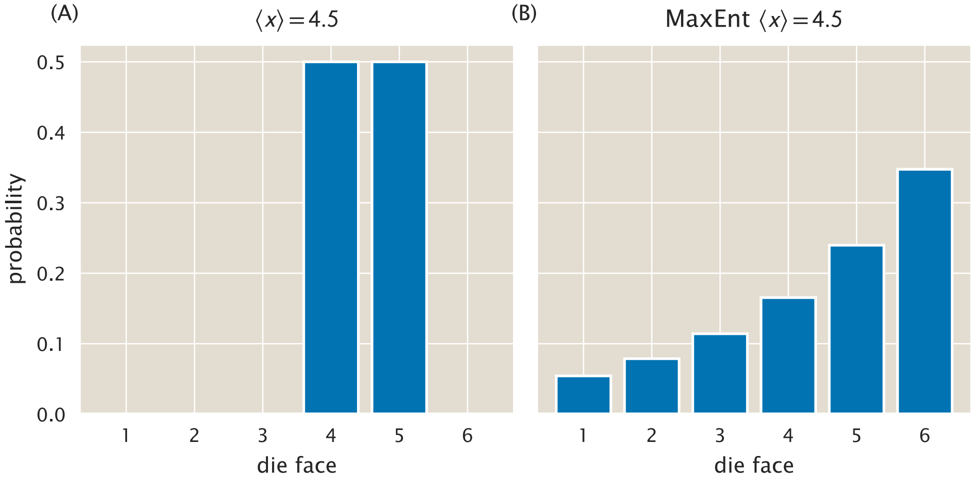

**Figure S15. Maximum entropy distribution of six-face die.** (A)biased distribution consistent with the constraint $\langle x \rangle = 4.5$. (B) MaxEnt distribution also consistent with the constraint.

##### S5.1.1 The mRNA and protein joint distribution

The MaxEnt principle can easily be extended to multivariate distributions. For our particular case we are interested in the mRNA and protein joint distribution $P(m, p)$. The definition of a moment $\langle m^x p^y \rangle$ is a natural extension of Eq. S118 of the form

$$\langle m^x p^y \rangle = \sum_m \sum_p m^x p^y P(m, p). \tag{S127}$$

As a consequence the MaxEnt joint distribution $P_H(m, p)$ is of the form

$$P_H(m, p) = \frac{1}{\mathcal{Z}} \exp\left( - \sum_{(x,y)} \lambda_{(x,y)} m^x p^y \right), \tag{S128}$$

where $\lambda_{(x,y)}$ is the Lagrange multiplier associated with the moment $\langle m^x p^y \rangle$, and again $\mathcal{Z}$ is the normalization constant given by

$$\mathcal{Z} = \sum_m \sum_p \exp\left( - \sum_{(x,y)} \lambda_{(x,y)} m^x p^y \right). \tag{S129}$$

Note that the sum in the exponent is taken over all available $(x, y)$ pairs that define the moment constraints for the distribution.

##### S5.2 The Bretthorst rescaling algorithm

The determination of the Lagrange multipliers suffer from a numerical under and overflow problem due to the difference in magnitude between the constraints. This becomes a problem when higher moments are taken into account. The resulting numerical values for the Lagrange multipliers end up being separated by several orders of magnitude. For routines such as Newton-Raphson or other minimization algorithms that can be used to find these Lagrange multipliers these different scales become problematic.

To get around this problem we implemented a variation to the algorithm due to G. Larry Bretthorst, E.T. Jaynes' last student. With a very simple argument we can show that linearly rescaling the constraints, the Lagrange multipliers and the "rules" for how to compute each of the moments, i.e. each of the individual products that go into the moment calculation, should converge to the same MaxEnt distribution. In order to see this let's consider again a univariate distribution $P_X(x)$ that we are trying to reconstruct given the first two moments $\langle x \rangle$, and $\langle x^2 \rangle$. The MaxEnt distribution can be written as

$$P_H(x) = \frac{1}{\mathcal{Z}} \exp\left(-\lambda_1 x - \lambda_2 x^2\right) = \frac{1}{\mathcal{Z}} \exp\left(-\lambda_1 x\right) \exp\left(-\lambda_2 x^2\right). \tag{S130}$$

We can always rescale the terms in any way and obtain the same result. Let's say that for some reason we want to rescale the quadratic terms by a factor $a$. We can define a new Lagrange multiplier $\lambda_2' \equiv \frac{\lambda_2}{a}$ that compensates for the rescaling of the terms, obtaining

$$P_H(x) = \frac{1}{\mathcal{Z}} \exp\left(-\lambda_1 x\right) \exp\left(-\lambda_2' a x^2\right). \tag{S131}$$

Computationally it might be more efficient to find the numerical value of $\lambda_2'$ rather than $\lambda_2$ maybe because it is of the same order of magnitude as $\lambda_1$. Then we can always multiply $\lambda_2'$ by $a$ to obtain back the constraint for our quadratic term. What this means is that that we can always rescale the MaxEnt problem to make it numerically more stable, then we can rescale back to obtain the value of the Lagrange multipliers. The key to the Bretthorst algorithm lies in the selection of what rescaling factor to choose in order to make the numerical inference more efficient.

Bretthorst's algorithm goes even further by further transforming the constraints and the variables to make the constraints orthogonal, making the computation much more effective. We now explain the implementation of the algorithm for our joint distribution of interest $P(m, p)$.

###### S5.2.1 Algorithm implementation

Let the $M \times N$ matrix $\mathbf{A}$ contain all the factors used to compute the moments that serve as constraints, where each entry is of the form

$$A_{ij} = m_i^{x_j} \cdot p_i^{y_j}. \tag{S132}$$

In other words, recall that to obtain any moment $\langle m^x p^y \rangle$ we compute

$$\langle m^x p^y \rangle = \sum_m \sum_p m^x p^y P(m, x). \tag{S133}$$

If we have $M$ possible $(m, p)$ pairs in our truncated sample space (because we can't include the sample space up to infinity) $\{(m, p)_1, (m, p)_2, \ldots (m, p)_N\}$, and we have $N$ exponent pairs $(x, y)$ corresponding to the $N$ moments used to constraint the maximum entropy distribution $\{(x, y)_1, (x, y)_2, \ldots, (x, y)_N\}$, then matrix $\mathbf{A}$ contains all the possible $M$ by $N$ terms of the form described in Eq. S132. Let also $\mathbf{v}$ be a vector of length $N$ containing all the constraints with each entry of the form

$$v_j = \langle m^{x_j} p^{y_j} \rangle, \tag{S134}$$

i.e. the information that we have about the distribution. That means that the constraint equation $\mathcal{L}$ to be used for this problem takes the form

$$\mathcal{L} = -\sum_i P_i \ln P_i + \lambda_0 \left(1 - \sum_i P_i\right) + \sum_{j>0} \lambda_j \left(v_j - \sum_i A_{ij} P_i\right), \tag{S135}$$

where $\lambda_0$ is the Lagrange multiplier associated with the normalization constraint, and $\lambda_j$ is the Lagrange multiplier associated with the $j^{\text{th}}$ constraint. This constraint equation is equivalent to Eq. S121, but now all the details of how to compute the moments are specified in matrix $\mathbf{A}$.

With this notation in hand we now proceed to rescale the problem. The first step consists of rescaling the terms to compute the entries of matrix $\mathbf{A}$. As mentioned before, this is the key feature of the Bretthorst algorithm; the particular choice of rescaling factor used in the algorithm empirically promotes that the rescaled Lagrange multipliers are of the same order of magnitude. The rescaling takes the form

$$A'_{ij} = \frac{A_{ij}}{G_j}, \tag{S136}$$

where $G_j$ serves to rescale the moments, providing numerical stability to the inference problem. Bretthorst proposes an empirical rescaling that satisfies

$$G_j^2 = \sum_i A_{ij}^2, \tag{S137}$$

or in terms of our particular problem

$$G_j^2 = \sum_m \sum_p \left(m^{x_j} p^{y_j}\right)^2. \tag{S138}$$

What this indicates is that each pair $m_i^{x_j} p_i^{y_j}$ is normalized by the square root of the sum of the all pairs of the same form squared.

Since we rescale the factors involved in computing the constraints, the constraints must also be rescaled simply as

$$v'_j = \langle m^{x_j} p^{y_j} \rangle' = \frac{\langle m^{x_j} p^{y_j} \rangle}{G_j}. \tag{S139}$$

The Lagrange multipliers must compensate this rescaling since at the end of the day the probability must add up to the same value. Therefore we rescale the $\lambda_j$ terms as

$$\lambda_j' = \lambda_j G_j, \tag{S140}$$

such that any $\lambda_j A_{ij} = \lambda_j' A_{ij}'$. If this empirical value for the rescaling factor makes the rescaled Lagrange multipliers $\lambda_j'$ be of the same order of magnitude, this by itself would already improve the algorithm convergence. Bretthorst proposes another linear transformation to make the optimization routine even more efficient. For this we generate orthogonal constraints that make Newton-Raphson and similar algorithms converge faster. The transformation is as follows

$$A_{ik}'' = \sum_j e_{jk} A_{ij}', \tag{S141}$$

for the entires of matrix $\mathbf{A}$, and

$$v_k'' = \sum_j e_{jk} u_j', \tag{S142}$$

for entires of the constraint vector $\mathbf{v}$, finally

$$\lambda_k'' = \sum_j e_{jk} \beta_j, \tag{S143}$$

for the Lagrange multipliers. Here $e_{jk}$ is the $j^{\text{th}}$ component of the $k^{\text{th}}$ eigenvector of the matrix $\mathbf{E}$ with entries

$$E_{kj} = \sum_i A_{ik}' A_{ij}'. \tag{S144}$$

This transformation guarantees that the matrix $\mathbf{A}''$ has the property

$$\sum_i A_{ij}'' A_{jk}'' = \beta_j \delta_{jk}, \tag{S145}$$

where $\beta_j$ is the $j^{\text{th}}$ eigenvalue of the matrix $\mathbf{E}$ and $\delta_{jk}$ is the Kronecker delta function. What this means is that, as desired, the constraints are orthogonal to each other, improving the algorithm convergence speed.

#### S5.3 Predicting distributions for simple repression constructs

Having explained the theoretical background along with the practical difficulties and a workaround strategy proposed by Bretthorst, we implemented the inference using the moments obtained from averaging over the variability along the cell cycle (See Appendix S4). Fig. S16 and Fig. S17 present these inferences for both mRNA and protein levels respectively for different values of the repressor-DNA binding energy and repressor copy numbers per cell. From these plots we can easily appreciate that despite the fact that the mean of each distribution changes as the induction level changes, there is a lot of overlap between distributions. This as a consequence means that at the single-cell level cells cannot perfectly resolve between different inputs.

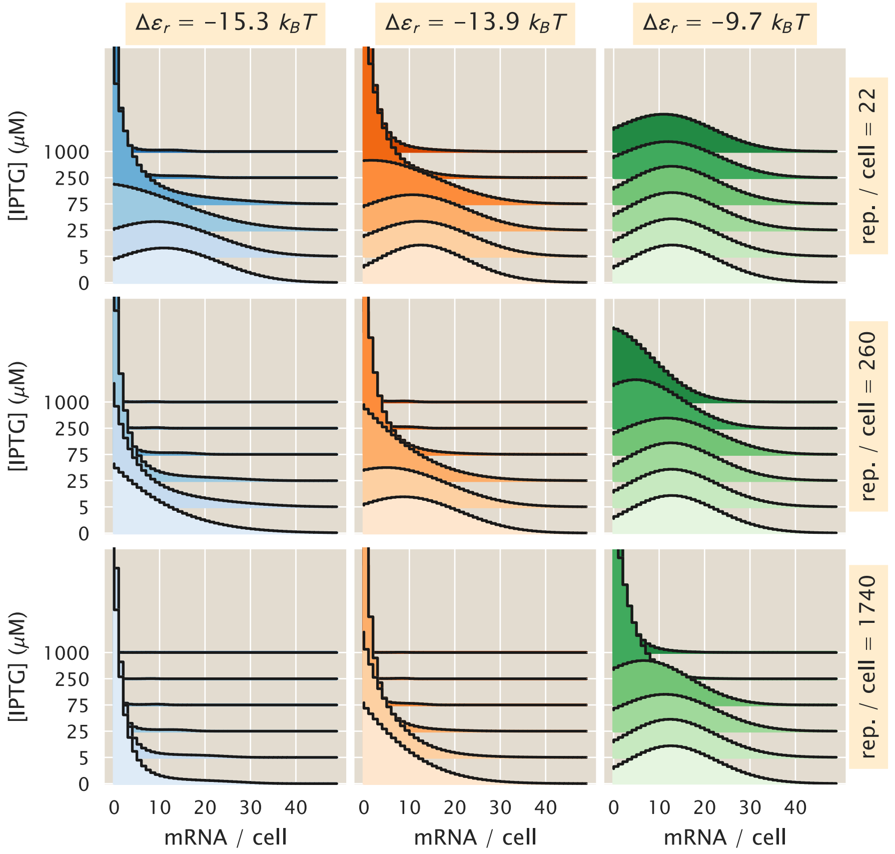

**Figure S16. Maximum entropy mRNA distributions for simple repression constructs.** mRNA distributions for different biophysical parameters. From left to right the repressor-DNA affinity decreases as defined by the three lacI operators O1 ($-15.3\ k_BT$), O2 ($-13.9\ k_BT$), and O3 ($-9.7\ k_BT$). From top to bottom the mean repressor copy number per cell increases. The curves on each plot represent different IPTG concentrations. Each distribution was fitted using the first three moments of the mRNA distribution.

##### S5.4 Comparison with experimental data

Now that we have reconstructed an approximation of the probability distribution $P(m, p)$ we can compare this with our experimental measurements. But just as detailed in Appendix S4.5 the single-cell microscopy measurements are given in arbitrary units of fluorescence. Therefore we cannot compare directly our predicted protein distributions with these values. To get around this issue we use the fact that the fold-change in gene expression that we defined as the ratio of the gene expression level in the presence of the repressor and the expression level of a knockout strain is a non-dimensional quantity. Therefore we normalize all of our single-cell measurements by the mean fluorescence value of the $\Delta lacI$ strain with the proper background fluorescence subtracted as explained in Appendix S4.5 for the noise measurements. In the case of the theoretical predictions of the protein distribution we also normalize each protein value by the predicted mean protein level $\langle p \rangle$, having now non-dimensional scales that can be directly compared. Fig. S18 shows the experimental (color curves) and theoretical (dark dashed line) cumulative distribution functions for the three $\Delta lacI$ strains. As in Fig. S12, we do not expect differences between the operators, but we explicitly plot them separately to make sure that this is the case. We can see right away that as we would expect given the limitations of the model

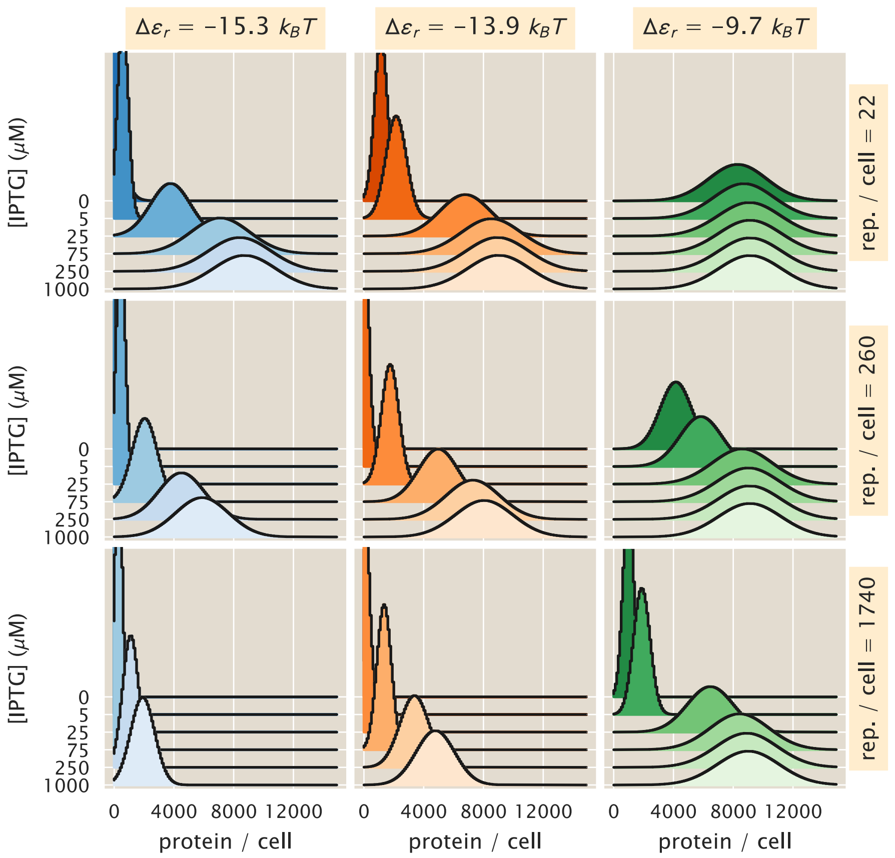

**Figure S17. Maximum entropy protein distributions for simple repression constructs.** Protein distributions for different biophysical parameters. From left to right the repressor-DNA affinity decreases as defined by the three lacI operators O1 ($-15.3\ k_BT$), O2 ($-13.9\ k_BT$), and O3 ($-9.7\ k_BT$). From top to bottom the mean repressor copy number per cell increases. The curves on each plot represent different IPTG concentrations. Each distribution was fitted using the first six moments of the protein distribution.

to accurately predict the noise and skewness of the distribution, the model doesn't accurately predict the data. Our model predicts a narrower distribution compared to what we measured with single-cell microscopy.

The same narrower prediction applies to the regulated promoters. Fig. S19, shows the theory-experiment comparison of the cumulative distribution functions for different repressor binding sites (different figures), repressor copy numbers (rows), and inducer concentrations (columns). In general the predictions are systematically narrower compared to the actual experimental data.

### S6 Gillespie simulation of master equation

(Note: The Python code used for the calculations presented in this section can be found in the following link as an annotated Jupyter notebook)

So far we have generated a way to compute an approximated form of the joint distribution of protein and mRNA $P(m, p)$ as a function of the moments of the distribution $\langle m^x p^y \rangle$. This is a non-conventional form to work with the resulting distribution of the master equation. A more conventional approach

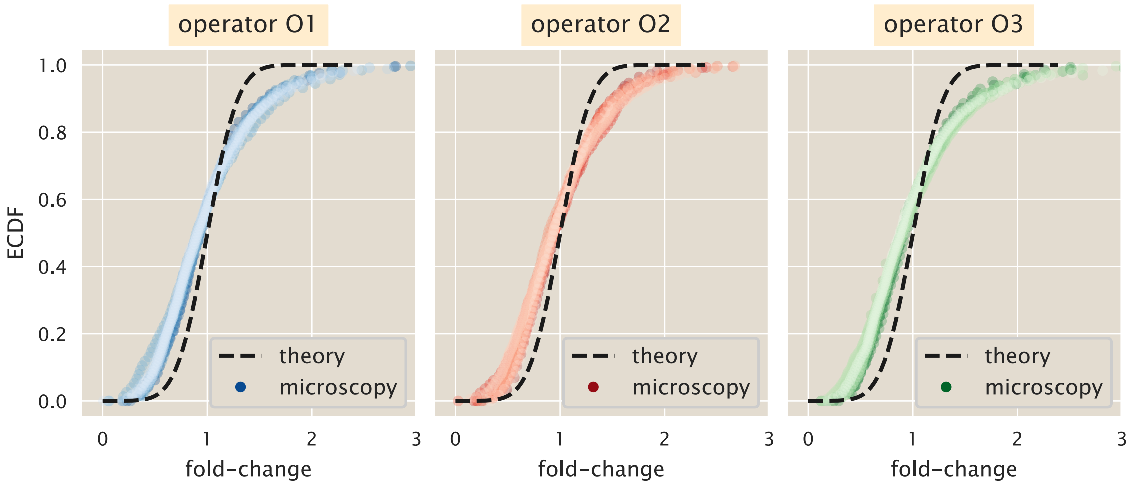

**Figure S18. Experiment vs. theory comparison for $\Delta lacI$ strain.** Example fold-change empirical cumulative distribution functions (ECDF) for strains with no repressors and different operators. The color curves represent single-cell microscopy measurements while the dashed black lines represent the theoretical distributions as reconstructed by the maximum entropy principle. The theoretical distributions were fitted using the first six moments of the protein distribution.

to work with master equations whose closed-form solutions are not known or not computable is to use stochastic simulations commonly known as Gillespie simulations. To benchmark the performance of our approach based on distribution moments and maximum entropy we implemented the Gillespie algorithm. Our implementation as detailed in the corresponding Jupyter notebook makes use of just-in-time compilation as implemented with the Python package numba.

#### S6.1 mRNA distribution with Gillespie simulations

To confirm that the implementation of the Gillespie simulation was correct we perform the simulation at the mRNA level for which the closed-form solution of the steady-state distribution is known as detailed in Appendix S2. Fig. S20 shows example trajectories of mRNA counts. Each of these trajectories were computed over several cell cyles, where the cell division was implemented generating a binomially distributed random variable that depended on the last mRNA count before the division event.

To check the implementation of our stochastic algorithm we generated several of these stochastic trajectories in order to reconstruct the mRNA steady-state distribution. These reconstructed distributions for a single- and double-copy of the promoter can be compared with Eq. S10 - the steady-state distribution for the two-state promoter. Fig. S21 shows the great agreement between the stochastic simulation and the analytical result, confirming that our implementation of the Gillespie simulation is correct.

#### S6.2 Protein distribution with Gillespie simulations

Having confirmed that our implementation of the Gillespie algorithm that includes the binomial partitioning of molecules reproduces analytical results we extended the implementation to include protein counts. Fig. S22 shows representative trajectories for both mRNA and protein counts over several cell cycles. Specially for the protein we can see that it takes several cell cycles for counts to converge to the dynamical steady-state observed with the deterministic moment equations. Once this steady-state is reached, the ensemble of trajectories between cell cycles look very similar.

From these trajectories we can compute the protein steady-state distribution, taking into account the cell-age distribution as detailed in Appendix S5. Fig. S23 shows the comparison between this distribution and the one generated using the maximum entropy algorithm. Despite the notorious differences between the distributions, the Gillespie simulation and the maximum entropy results are indistinguishable in terms of the mean, variance, and skewness of the distribution. We remind the reader that the maximum entropy is an approximation of the distribution that gets better the more moments we add. We therefore claim that the approximation works sufficiently well for our purpose. The enormous advantage of the maximum entropy approach comes from the computation time. for the number of distributions that were needed for our calculations the Gillespie algorithm proved to be a very inefficient method given the large sample space. Our maximum entropy approach reduces the computation time by several orders of magnitude, allowing us to extensively explore different parameters of the regulatory model.

### S7 Computational determination of the channel capacity

(Note: The Python code used for the calculations presented in this section can be found in the following link as an annotated Jupyter notebook)

In this section we detail the computation of the channel capacity of the simple genetic circuit shown in Fig. 5. As detailed in Section 1.6 the channel capacity is defined as the mutual information between input $c$ and output $p$ maximized over all possible input distributions $P(c)$ [3]. In principle there are an infinite number of input distributions, so the task of finding $\hat{P}(c)$, the input distribution at channel capacity, requires an algorithmic approach that guarantees the convergence to this distribution. Tkačik, Callan and Bialek developed a clever analytical approximation to find the $\hat{P}(c)$ distribution [17]. The validity of their so-called small noise approximation requires the standard deviation of the output distribution $P(p \mid c)$ to be much smaller than the domain of the distribution. For our particular case such condition is not satisfied given the spread of the inferred protein distributions shown in Fig. 4.

Fortunately there exists a numerical algorithm to approximate $\hat{P}(c)$ for discrete distributions. In 1972 Richard Blahut and Suguru Arimoto independently came up with an algorithm mathematically shown to converge to $\hat{P}(c)$ [44]. To compute both the theoretical and the experimental channel capacity shown in Fig. 5, we implemented Blahut's algorithm. In the following section we detail the definitions needed for the algorithm. Then we detail how to compute the experimental channel capacity when the bins of the distribution are not clear given the intrinsic arbitrary nature of microscopy fluorescence measurements.

#### S7.1 Blahut's algorithm

Following [44] we implemented the algorithm to compute the channel capacity. We define $\mathbf{p_c}$ to be an array containing the probability of each of the input inducer concentrations (twelve concentrations, See Methods). Each entry $j$ of the array is then of the form

$$p_c^{(j)} = P(c = c_j), \tag{S146}$$

with $j \in \{1, 2, \ldots, 12\}$. The objective of the algorithm is to find the entries $p_c^{(j)}$ that maximize the mutual information between inputs and outputs. We also define $\mathbf{Q}$ to be a $|\mathbf{p_c}|$ by $|\mathbf{p_{p|c}}|$ matrix, where $|\cdot|$ specifies the length of the array, and $\mathbf{p_{p|c}}$ is an array containing the probability distribution of an output given a specific value of the input. In other words, the matrix $\mathbf{Q}$ recollects all of the individual output distribution arrays $\mathbf{p_{p|c}}$ into a single object. Then each entry of the matrix $\mathbf{Q}$ is of the form

$$Q^{(i,j)} = P(p = p_i \mid c = c_j). \tag{S147}$$

For the case of the theoretical predictions of the channel capacity (Solid lines in Fig. 5) the entries of matrix $\mathbf{Q}$ are given by the inferred maximum entropy distributions as shown in Fig. 4. In the next section we will discuss how to define this matrix for the case of the single-cell fluorescence measurements. Having defined these matrices we proceed to implement the algorithm shown in Figure 1 of [44].

##### S7.2 Channel capacity from arbitrary units of fluorescence

A difficulty when computing the channel capacity between inputs and outputs from experimental data is that ideally we would like to compute

$$C(g; c) \equiv \sup_{P(c)} I(g; c), \tag{S148}$$

where $g$ is the gene expression level, and $c$ is the inducer concentration. But in reality we are computing

$$C(f(g); c) \equiv \sup_{P(c)} I(f(g); c), \tag{S149}$$

where $f(g)$ is a function of gene expression that has to do with our mapping from the YFP copy number to some arbitrary fluorescent value as computed from the images taken with the microscope. The data processing inequality, as derived by Shannon himself, tells us that for a Markov chain of the form $c \rightarrow g \rightarrow f(g)$ it must be true that [3]

$$I(g; c) \geq I(f(g); c), \tag{S150}$$

meaning that information can only be lost when mapping from the real relationship between gene expression and inducer concentration to a fluorescence value.

On top of that, given the limited number of samples that we have access to when computing the channel capacity, there is a bias in our estimate given this undersampling. The definition of accurate unbiased descriptors of the mutual information is still an area of active research. For our purposes we will use the method described in [68]. The basic idea of the method is to write the mutual information as a series expansion in terms of inverse powers of the sample size, i.e.

$$I_{\text{biased}} = I_{\infty} + \frac{a_1}{N} + \frac{a_2}{N^2} + \cdots, \tag{S151}$$

where $I_{\text{biased}}$ is the biased estimate of the mutual information as computed from experimental data, $I_{\infty}$ is the quantity we would like to estimate, being the unbiased mutual information when having access to infinity number of experimental samples, and the coefficients $a_i$ depend on the underlying distribution of the signal and the response. This is an empirical choice to be tested. Intuitively this choice satisfies the limit that as the number of samples from the distribution grows, the empirical estimate of the mutual information $I_{\text{biased}}$ should get closer to the actual value $I_\infty$.

In principle for a good number of data points the terms of higher order become negligible. So we can write the mutual information as

$$I_{\text{biased}} \approx I_\infty + \frac{a_1}{N} + \mathcal{O}(N^{-2}). \tag{S152}$$

This means that if this particular arbitrary choice of functional form is a good approximation, when computing the mutual information for varying number of samples - by taking subsamples of the experimental data - we expect to find a linear relationship as a function of the inverse of these number of data points. From this linear relationship the intercept is a bias-corrected estimate of the mutual information. We can therefore bootstrap the data by taking different sample sizes and then use the Blahut-Arimoto algorithm we implemented earlier to estimate the biased channel capacity. We can then fit a line and extrapolate for when $1/N = 0$ which corresponds to our unbiased estimate of the channel capacity.

Let's go through each of the steps to illustrate the method. Fig. S24 show a typical data set for a strain with an O2 binding site ($\Delta\varepsilon_r = -13.9\ k_BT$) and $R = 260$ repressors per cell. Each of the distributions in arbitrary units is binned into a specified number of bins to build matrix $\mathbf{Q}$.

Given a specific number of bins used to construct $\mathbf{Q}$, we subsample a fraction of the data and compute the channel capacity for such matrix using the Blahut-Arimoto algorithm. Fig. S25 shows an example where 50% of the data on each distribution from Fig. S24 was sampled and binned into 100 equal bins. The counts on each of these bins are then normalized and used to build matrix $\mathbf{Q}$ that is then fed to the Blahut-Arimoto algorithm. We can see that for these 200 bootstrap samples the channel capacity varies by $\approx 0.1$ bits. Not a significant variability, nevertheless we consider that it is important to bootstrap the data multiple times to get a better estimate of the channel capacity.

Eq. S152 tells us that if we subsample each of the distributions from Fig. S24 at different fractions, and plot them as a function of the inverse sample size we will find a linear relationship if the expansion of the mutual information is valid. To test this idea we repeated the bootstrap estimate of Fig. S25 sampling 10%, 20%, and so on until taking 100% of the data. We repeated this for different number of bins since a priori for arbitrary units of fluorescence we do not have a way to select the optimal number of bins. Fig. S26 shows the result of these estimates. We can see that the linear relationship proposed in Eq. S152 holds true for all number of bins selected. We also note that the value of the intercept of the linear regression varies depending on the number of bins.

To address the variability in the estimates of the unbiased channel capacity $I_\infty$ we again follow the methodology suggested in [68]. We perform the data subsampling and computation of the channel capacity for a varying number of bins. As a control we perform the same procedure with shuffled data, where the structure that connects the fluorescence distribution to the inducer concentration input is lost. The expectation is that this control should give a channel capacity of zero if the data is not "over-binned." Once the number of bins is too high, we would expect some structure to emerge in the data that would cause the Blahut-Arimoto algorithm to return non-zero channel capacity estimates.

Fig. S27 shows the result of the unbiased channel capacity estimates obtained for the data shown in Fig. S24. For the blue curve we can distinguish three phases:

1. A rapid increment from 0 bits to about 1.5 bits as the number of bins increases.

2. A flat region between ≈ 50 and 1000 bins.
3. A second rapid increment for large number of bins.

We can see that the randomized data presents two phases only:

1. A flat region where there is, as expected no information being processed since the structure of the data was lost when the data was shuffled.
2. A region with fast growth of the channel capacity as the over-binning generates separated peaks on the distribution, making it look like there is structure in the data.

We take the flat region of the experimental data (≈ 100 bins) to be our best unbiased estimate of the channel capacity from this experimental dataset.

##### S7.3 Assumptions involved in the computation of the channel capacity

An interesting suggestion by Professor Gasper Tkacik was to dissect the different physical assumptions that went into the construction of the input-output function $P(p \mid c)$, and their relevance when comparing the theoretical channel capacities with the experimental inferences. In what follows we describe the relevance of four important aspects that all affect the predictions of the information processing capacity.

**(i) Cell cycle variability.** We think that the inclusion of the gene copy number variability during the cell cycle and the non-Poissoninan protein degradation is a key component to our estimation of the input-output functions and as a consequence of the channel capacity. This variability in gene copy number is an additional source of noise that systematically decreases the ability of the system to resolve different inputs. The absence of the effects that the gene copy number variability and the protein partition has on the information processing capacity leads to an overestimate of the channel capacity as shown in Fig. S28. Only when these noise sources are included in our inferences is that we get to capture the experimental channel capacities with no further fit parameters.

**(ii) Non-Gaussian noise distributions.** For the construction of the probability distributions used in the main text (Fig. 4) we utilized the first 6 moments of the protein distribution. The maximum entropy formalism tells us that the more constraints we include in the inference, the closer the maximum entropy distribution will be to the real distribution. But *a priori* there is no way of knowing how many moments should be included in order to capture the essence of the distribution. In principle two moments could suffice to describe the entire distribution as happens with the Gaussian distribution. To compare the effect that including more or less constraints on the maximum entropy inference we constructed maximum entropy distributions using an increasing number of moments from 2 to 6. We then computed the Kullback-Leibler divergence $D_{KL}$ of the form

$$D_{KL}(P_6(p \mid c)||P_i(p \mid c)) = \sum_p P_6(p \mid c) \log_2 \frac{P_6(p \mid c)}{P_i(p \mid c)}, \tag{S153}$$

where $P_i(p \mid c)$ is the maximum entropy distribution constructed with the first $i$ moments, $i \in \{2, 3, 4, 5, 6\}$. Since the Kullback-Leibler divergence $D_{KL}(P||Q)$ can be interpreted as the amount of information lost by assuming the incorrect distribution $Q$ when the correct distribution is $P$, we used this metric as a way of how much information we would have lost by using less constraints compared to the six moments used in the main text.

Fig. S29 shows this comparison for different operators and repressor copy numbers. We can see from here that using less moments as constraints gives basically the same result. This is because most of the values of the Kullback-Leibler divergence are significantly smaller than 0.1 bits, and the entropy of these distributions is in general $> 10$ bits, so we would lose less than 1% of the information contained in these distributions by utilizing only two moments as constraints. Therefore the use of non-Gaussian noise is not an important feature for our inferences.

**(iii) Multi-state promoter.** This particular point is something that we are still exploring from a theoretical perspective. We have shown that in order to capture the single-molecule mRNA FISH data a single-state promoter wouldn't suffice. This model predicts a Poisson distribution as the steady-state and the data definitely shows super Poissonian noise. Given the bursty nature of gene expression we opt to use a two-state promoter where the states reflect effective transcriptionally "active" and "inactive" states. We are currently exploring alternative formulations of this model to turn it into a single state with a geometrically distributed burst-size.

**(iv) Optimal vs Log-flat Distributions.** The relevance of having use the Blahut-Arimoto algorithm to predict the maximum mutual information between input and outputs was just to understand the best case scenario. We show the comparison between theoretical and experimental input-output functions $P(p \mid c)$ in Fig. S19. Given the good agreement between these distributions we could compute the mutual information $I(c; p)$ for any arbitrary input distribution $P(c)$ and obtain a good agreement with the corresponding experimental mutual information.

The reason we opted to specifically report the mutual information at channel capacity was to put the results in a context. By reporting the upper bound in performance of these genetic circuits we can start to dissect how different molecular parameters such as repressor-DNA binding affinity or repressor copy number affect the ability of this genetic circuit to extract information from the environmental state.

### S8 Empirical fits to noise predictions

(Note: The Python code used for the calculations presented in this section can be found in the following link as an annotated Jupyter notebook)

In Fig. 3(C) in the main text we show that our minimal model has a systematic deviation on the gene expression noise predictions compared to the experimental data. This systematics will need to be addressed on an improved version of the minimal model presented in this work. To guide the insights into the origins of this systematic deviation in this appendix we will explore empirical modifications of the model to improve the agreement between theory and experiment.

#### S8.1 Multiplicative factor for the noise

The first option we will explore is to modify our noise predictions by a constant multiplicative factor. This means that we assume the relationship between our minimal model predictions and the data for noise in gene expression are of the from

$$\text{noise}_{\text{exp}} = \alpha \cdot \text{noise}_{\text{theory}}, \tag{S154}$$

where $\alpha$ is a dimensionless constant to be fit from the data. The data, especially in Fig. S12 suggests that our predictions are within a factor of $\approx$ two from the experimental data. To further check that intuition we performed a weighted linear regression between the experimental and theoretical noise measurements. The weight for each datum was taken to be proportional to the bootstrap errors in the noise estimate, this to have poorly determined noises weigh less during the regression. The result of this regression with no intercept shows exactly that a factor of two systematically improves the theoretical vs. experimental predictions. Fig. S30 shows the improved agreement when the theoretical predictions for the noise are multiplied by $\approx 1.5$.

For completeness Fig. S31 shows the noise in gene expression as a function of the inducer concentration including this factor of $\approx 1.5$. It is clear that overall a simple multiplicative factor improves the predictive power of the model.

#### S8.2 Additive factor for the noise

As an alternative way to empirically improve the predictions of our model we will now test the idea of an additive constant. What this means is that our minimal model underestimates the noise in gene expression as

$$\text{noise}_{\text{exp}} = \beta + \text{noise}_{\text{theory}}, \tag{S155}$$

where $\beta$ is an additive constant to be determined from the data. As with the multiplicative constant we performed a regression to determine this empirical additive constant comparing experimental and theoretical gene expression noise values. We use the error in the 95% bootstrap confidence interval as a weight for the linear regression. Fig. S32 shows the resulting theoretical vs. experimental noise where $\beta \approx 0.2$. We can see a great improvement in the agreement between theory and experiment with this additive constant

For completeness Fig. S33 shows the noise in gene expression as a function of the inducer concentration including this additive factor of $\beta \approx 0.2$. If anything, the additive factor seems to improve the agreement between theory and data even more than the multiplicative factor.

#### S8.3 Correction factor for channel capacity with multiplicative factor

As seen in Appendix S4 a constant multiplicative factor can reduce the discrepancy between the model predictions and the data with respect to the noise (standard deviation / mean) in protein copy number. To find the equivalent correction would be for the channel capacity requires gaining insights from the so-called small noise approximation [17]. The small noise approximation assumes that the input-output function can be modeled as a Gaussian distribution in which the standard deviation is small. Using these assumptions one can derive a closed-form for the channel capacity. Although our data and model predictions do not satisfy the requirements for the small noise approximation, we can gain some intuition for how the channel capacity would scale given a systematic deviation in the cell-to-cell variability predictions compared with the data.

Using the small noise approximation one can derive the form of the input distribution at channel capacity $P^*(c)$. To do this we use the fact that there is a deterministic relationship between the input inducer concentration $c$ and the mean output protein value $\langle p \rangle$, therefore we can work with $P(\langle p \rangle)$ rather than $P(c)$ since the deterministic relation allows us to write

$$P(c)dc = P(\langle p \rangle)d\langle p \rangle\,. \tag{S156}$$

Optimizing over all possible distributions $P(\langle p \rangle)$ using calculus of variations results in a distribution of the form

$$P^*(\langle p \rangle) = \frac{1}{\mathcal{Z}}\frac{1}{\sigma_p(\langle p \rangle)}, \tag{S157}$$

where $\sigma_p(\langle p \rangle)$ is the standard deviation of the protein distribution as a function of the mean protein expression, and $\mathcal{Z}$ is a normalization constant defined as

$$\mathcal{Z} \equiv \int_{\langle p(c=0)\rangle}^{\langle p(c\rightarrow\infty)\rangle} \frac{1}{\sigma_p(\langle p \rangle)} d\langle p \rangle\,. \tag{S158}$$

Under these assumptions the small noise approximation tells us that the channel capacity is of the form [17]

$$I = \log_2\left(\frac{\mathcal{Z}}{\sqrt{2\pi e}}\right). \tag{S159}$$

From the theory-experiment comparison in Appendix S4 we know that the standard deviation predicted by our model is systematically off by a factor of two compared to the experimental data, i.e.

$$\sigma_p^{\text{exp}} = 2\sigma_p^{\text{theory}}. \tag{S160}$$

This then implies that the normalization constant $\mathcal{Z}$ between theory and experiment must follow a relationship of the form

$$\mathcal{Z}^{\text{exp}} = \frac{1}{2}\mathcal{Z}^{\text{theory}}. \tag{S161}$$

With this relationship the small noise approximation would predict that the difference between the experimental and theoretical channel capacity should be of the form

$$I^{\text{exp}} = \log_2\left(\frac{\mathcal{Z}^{\text{exp}}}{\sqrt{2\pi e}}\right) = \log_2\left(\frac{\mathcal{Z}^{\text{theory}}}{\sqrt{2\pi e}}\right) - \log_2(2). \tag{S162}$$

Therefore under the small noise approximation we would expect our predictions for the channel capacity to be off by a constant of 1 bit ($\log_2(2)$) of information. Again, the conditions for the small noise approximation do not apply to our data given the intrinsic level of cell-to-cell variability in the system, nevertheless what this analysis tells is is that we expect that an additive constant should be able to explain the discrepancy between our model predictions and the experimental channel capacity. To test this hypothesis we performed a "linear regression" between the model predictions and the experimental channel capacity with a fixed slope of 1. The intercept of this regression, -0.56 bits, indicates the systematic deviation we expect should explain the difference between our model and the data. Fig. S34 shows the comparison between the original predictions shown in Fig. 5(A) and the resulting predictions with this shift. Other than the data with zero channel capacity, this shift is able to correct the systematic deviation for all data. We therefore conclude that our model ends up underestimating the experimentally determined channel capacity by a constant amount of 0.43 bits.

### S9 Derivation of the cell age distribution

E. O. Powell first derive in 1956 the distribution of cell age for a cell population growing steadily in the exponential phase [40]. This distribution is of the form

$$P(a) = \ln(2) \cdot 2^{1-a}, \tag{S163}$$

where $a \in [0, 1]$ is the fraction of the cell cycle, 0 being the moment right after the mother cell divides, and 1 being the end of the cell cycle just before cell division. In this section we will reproduce and expand the details on each of the steps of the derivation.

For an exponentially growing bacterial culture, the cells satisfy the growth law

$$\frac{dn}{dt} = \mu n, \tag{S164}$$

where $n$ is the number of cells and $\mu$ is the growth rate in units of time$^{-1}$. We begin by defining $P(a)$ to be the probability density function of a cell having age $a$. At time zero of a culture in exponential growth, i.e. the time when we start considering the growth, not the initial condition of the culture, there are $NP(a)da$ cells with age range between $[a, a + da]$. In other words, for $N \gg 1$ and $da \ll a$

$$NP(a \leq x \leq a + da) \approx NP(a)da. \tag{S165}$$

We now define

$$F(\tau) = \int_{\tau}^{\infty} f(\xi) d\xi, \tag{S166}$$

as the fraction of cells whose division time is greater than $\tau$. This is because in principle not all cells divide exactly after $\tau$ minutes, but there is a distribution function $f(\tau)$ for the division time after birth. Empirically it has been observed that a generalize Gamma distribution fits well to experimental data on cell division time, but we will worry about this specific point later on.

From the definition of $F(\tau)$ we can see that if a cell reaches an age $a$, the probability of surviving to an age $a + t$ without dividing is given by

$$P(\text{age} = (a + t) \mid \text{age} = a) = F(a + t \mid a) = \frac{F(a + t)}{F(a)}. \tag{S167}$$

This result comes simply from the definition of conditional probability. Since $F(a)$ is the probability of surviving $a$ or more minutes without dividing, by the definition of conditional probability we have that

$$F(a + t \mid a) = \frac{F(a, a + t)}{F(a)}, \tag{S168}$$

where $F(a, a+t)$ is the joint probability of surviving $a$ minutes and $a+t$ minutes. But the probability of surviving $a + t$ minutes or more implies that the cell already survived $a$ minutes, therefore the information is redundant and we have

$$F(a, a + t) = F(a + t). \tag{S169}$$

This explains Eq. S167. From this equation we can find that out of the $NP(a)da$ cells with age $a$ only a fraction

$$[NP(a)da] \, F(a + t \mid a) = NP(a)\frac{F(a + t)}{F(a)}da \tag{S170}$$

will survive without dividing until time $a + t$. During that time interval $t$ the culture has passed from $N$ cells to $Ne^{\mu t}$ cells given the assumption that they are growing exponentially. The survivors $NP(a)F(a + t \mid a)da$ then represent a fraction of the total number of cells

$$\frac{\#\text{ survivors}}{\#\text{ total cells}} = \frac{[NP(a)da]\,F(a+t \mid a)}{Ne^{\mu t}} = P(a)\frac{F(a+t)}{F(a)}da\frac{1}{e^{\mu t}}, \tag{S171}$$

and their ages lie in the range $[a + t, a + t + da]$. Since we assume that the culture is in steady state then it follows that the fraction of cells that transitioned from age $a$ to age $a + t$ must be $P(a + t)da$. Therefore we have a difference equation - the discrete analogous of a differential equation - of the form

$$P(a+t)da = P(a)\frac{F(a+t)}{F(a)}e^{-\mu t}da. \tag{S172}$$

What this equation shows is a relationship that connects the probability of having a life time of $a + t$ with a probability of having a shorter life time $a$ and the growth of the population. If we take $t$ to be very small, specifically if we assume $t \ll \mu^{-1}$ we can Taylor expand around $a$ the following terms:

$$F(a+t) \approx F(a) + \frac{dF}{da}t, \tag{S173}$$

$$P(a+t) \approx P(a) + \frac{dP}{da}t, \tag{S174}$$

and

$$e^{-\mu t} \approx 1 - \mu t. \tag{S175}$$

Substituting these equations into Eq. S172 gives

$$P(a) + \frac{dP}{da}t = P(a)\left(\frac{F(a) + \frac{dF}{da}t}{F(a)}\right)(1 - \mu t). \tag{S176}$$

This can be rewritten as

$$\frac{1}{P(a)}\frac{dP}{da} = \frac{1}{F(a)}\frac{dF}{da} - \mu - \frac{\mu t}{F(a)}\frac{dF}{da}. \tag{S177}$$

Since we assumed $t \ll \mu^{-1}$ we then approximate the last term to be close to zero. We can then simplify this result into

$$\frac{1}{P(a)}\frac{dP}{da} = \frac{1}{F(a)}\frac{dF}{da} - \mu. \tag{S178}$$

Integrating both sides of the equation with respect to $a$ gives

$$\ln P(a) = \ln F(a) - \mu a + C, \tag{S179}$$

where $C$ is the integration constant. Exponentiating both sides gives

$$P(a) = C' F(a) e^{-\mu a}. \tag{S180}$$

Where $C' \equiv e^{C}$. To obtain the value of the unknown constant we recall that $F(0) = 1$ since the probability of having a life equal or longer than zero must add up to one, therefore we have that $P(0) = C'$. This gives then

$$P(a) = P(0)e^{-\mu a}F(a). \tag{S181}$$

Substituting the definition of $F(a)$ gives

$$P(a) = P(0)e^{-\mu a} \int_a^\infty f(\xi) d\xi. \tag{S182}$$

The last step of the derivation involves writing $P(0)$ and the growth rate $\mu$ in terms of the cell cycle length distribution $f(\tau)$.

The growth rate of the population cell number (not the growth of cell mass) is defined as the number of cell doublings per unit time divided by the number of cells. This is more clear to see if we write Eq. S164 as a finite difference

$$\frac{N(t + \Delta t) - N(t)}{\Delta t} = \mu N(t). \tag{S183}$$

If the time $\Delta t$ is the time interval it takes to go from $N$ to $2N$ cells we have

$$\frac{2N - N}{\Delta t} = \mu N. \tag{S184}$$

Solving for $\mu$ gives

$$\mu = \overbrace{\frac{2N - N}{\Delta t}}^{\text{\# doubling events per unit time}} \overbrace{\frac{1}{N}}^{\frac{1}{\text{population size}}}. \tag{S185}$$

We defined $F(a)$ to be the probability of a cell reaching an age $a$ or greater. For a cell to reach an age $a + da$ we can then write

$$F(a + da) = \int_{a+da}^\infty f(\xi) d\xi = \int_a^\infty f(\xi) d\xi - \int_a^{a+da} f(\xi) d\xi. \tag{S186}$$

We can approximate the second term on the right hand side to be

$$\int_a^{a+da} f(\xi) d\xi \approx f(a) da, \tag{S187}$$

for $da \ll a$, obtaining

$$F(a + da) \approx F(a) - f(a) da. \tag{S188}$$

What this means is that from the original fraction of cells $F(a)$ with age $a$ or greater a fraction $f(a)da/F(a)$ will not reach age $(a + da)$ because they will divide. So out of the $NP(a)$ cells that reached exactly age $a$, the number of doubling events on a time interval $da$ is given by

$$\text{\# doublings of cells of age } a \text{ on interval } da = \overbrace{NP(a)}^{\text{\# cells of age } a} \overbrace{\frac{f(a)da}{F(a)}}^{\text{fraction of doublings per unit time}}. \tag{S189}$$

The growth rate then is just the sum (integral) of each age contribution to the total number of doublings. This is

$$\mu = \frac{1}{N} \int_0^\infty NP(a) \frac{f(a)da}{F(a)}. \tag{S190}$$

Substituting Eq. S181 gives

$$\mu = \int_0^\infty [P(0)e^{-\mu a}F(a)]\frac{f(a)da}{F(a)} = \int_0^\infty P(0)e^{-\mu a}f(a)da. \tag{S191}$$

We now have the growth rate $\mu$ written in terms of the cell cycle length probability distribution $f(a)$ and the probability $P(0)$. Since $P(a)$ is a probability distribution it must be normalized, i.e.

$$\int_0^\infty P(a)da = 1. \tag{S192}$$

Substituting Eq. S181 into this normalization constraint gives

$$\int_0^\infty P(0)e^{-\mu a}F(a)da = 1. \tag{S193}$$

From here we can integrate the left hand side by parts. We note that given the definition of $F(a)$, the derivative with respect to $a$ is $-f(a)$ rather than $f(a)$. This is because if we write the derivative of $F(a)$ we have

$$\frac{dF(a)}{da} \equiv \lim_{da\to 0}\frac{F(a+da)-F(a)}{da}. \tag{S194}$$

Substituting the definition of $F(a)$ gives

$$\frac{dF(a)}{da} = \lim_{da\to 0}\frac{1}{da}\left[\int_{a+da}^\infty f(\xi)d\xi - \int_a^\infty f(\xi)d\xi\right]. \tag{S195}$$

This difference in the integrals can be simplified to

$$\lim_{da\to 0}\frac{1}{da}\left[\int_{a+da}^\infty f(\xi)d\xi - \int_a^\infty f(\xi)d\xi\right] \approx \frac{-f(a)da}{da} = -f(a). \tag{S196}$$

Taking this into account we now perform the integration by parts obtaining

$$P(0)\left[\frac{e^{-\mu t}}{-\mu}F(a)\right]_0^\infty - P(0)\int_0^\infty \frac{e^{-\mu a}}{-\mu}(-f(a))da = 1. \tag{S197}$$

On the first term on the left hand side we have that as $a \to \infty$, both terms $e^{-\mu a}$ and $F(a)$ go to zero. We also have that $e^{\mu 0} = 1$ and $F(0) = 1$. This results in

$$\frac{P(0)}{\mu} - P(0)\int_0^\infty \frac{e^{-\mu a}}{\mu}f(a)da = 1. \tag{S198}$$

The second term on the left hand side is equal to Eq. S191 since

$$\mu = \int_0^\infty P(0)e^{-\mu a}f(a)da \Rightarrow 1 = \int_0^\infty P(0)\frac{e^{-\mu a}}{\mu}f(a)da. \tag{S199}$$

This implies that on Eq. S197 we have

$$\frac{P(0)}{\mu} - 1 = 1 \Rightarrow P(0) = 2\mu. \tag{S200}$$

With this result in hand we can rewrite Eq. S182 as

$$P(a) = 2\mu e^{-\mu a} \int_a^{\infty} f(\xi) d\xi. \tag{S201}$$

Also we can rewrite the result for the growth rate $\mu$ on Eq. S191 as

$$\mu = 2\mu \int_0^{\infty} e^{-\mu a} f(a) da \Rightarrow 2 \int_0^{\infty} e^{-\mu a} f(a) da = 1. \tag{S202}$$

As mentioned before the distribution $f(a)$ has been empirically fit to a generalize Gamma distribution. But if we assume that our distribution has almost negligible dispersion around the mean average doubling time $a = \tau_d$, we can approximate $f(a)$ as

$$f(a) = \delta(a - \tau_d), \tag{S203}$$

a Dirac delta function. Applying this to Eq. S202 results in

$$2 \int_0^{\infty} e^{-\mu a} \delta(a - \tau_a) da = 1 \Rightarrow 2e^{-\mu \tau_d} = 1. \tag{S204}$$

Solving for $\mu$ gives

$$\mu = \frac{\ln 2}{\tau_d}. \tag{S205}$$

This delta function approximation for $f(a)$ has as a consequence that

$$F(a) = \begin{cases} 1 \text{ for } a \in [0, \tau_d], \\ 0 \text{ for } a > \tau_d. \end{cases} \tag{S206}$$

Fianlly we can rewrite Eq. S201 as

$$P(a) = 2 \left( \frac{\ln 2}{\tau_d} \right) e^{-\frac{\ln 2}{\tau_d} a} \int_a^{\infty} \delta(\xi - \tau_d) d\xi \Rightarrow= 2 \ln 2 \cdot 2^{\frac{-a}{\tau_d}}. \tag{S207}$$

Simplifying this we obtain

$$P(a) = \begin{cases} \ln 2 \cdot 2^{1 - \frac{a}{\tau_d}} \text{ for } a \in [0, \tau_d], \\ 0 \text{ otherwise.} \end{cases} \tag{S208}$$

This is the equation we aimed to derive. The distribution of cell ages over the cell cycle.

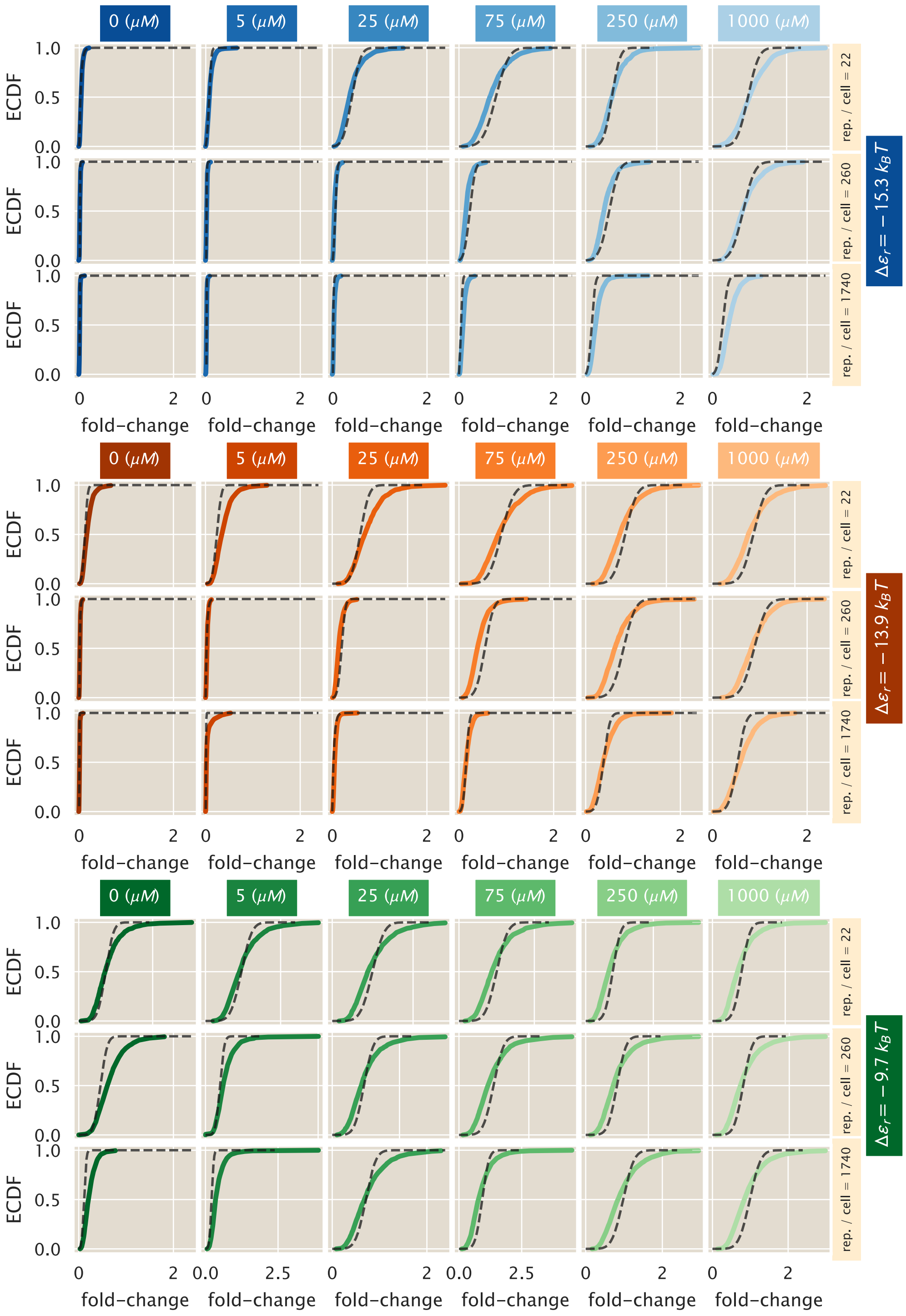

**Figure S19. Experiment vs. theory comparison for regulated promoters.** Example fold-change empirical cumulative distribution functions (ECDF) for regulated strains with the three operators (different colors) as a function of repressor copy numbers (rows) and inducer concentrations (columns). The color curves represent single-cell microscopy measurements while the dashed black lines represent the theoretical distributions as reconstructed by the maximum entropy principle. The theoretical distributions were fitted using the first six moments of the protein distribution

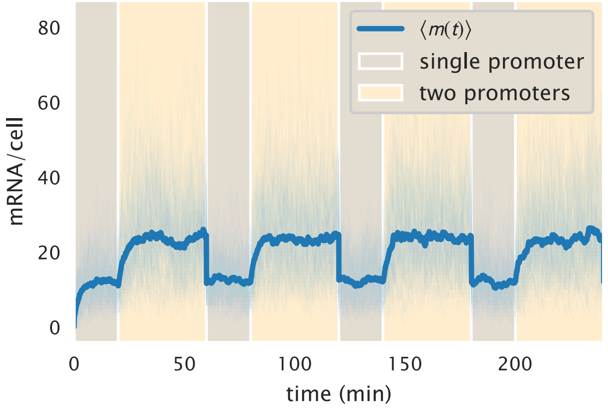

**Figure S20. Stochastic trajectories of mRNA counts.** 100 stochastic trajectories generated with the Gillespie algorithm for mRNA counts over time for a two-state unregulated promoter. Cells spend a fraction of the cell cycle with a single copy of the promoter (light brown) and the rest of the cell cycle with two copies (light yellow). When trajectories reach a new cell cycle, the mRNA counts undergo a binomial partitioning to simulate the cell division.

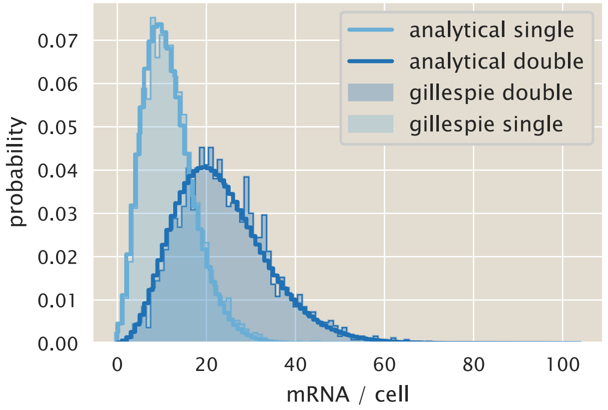

**Figure S21. Comparison of analytical and simulated mRNA distribution.** Solid lines show the steady-state mRNA distributions for one copy (light blue) and two copies of the promoter (dark blue) as defined by Eq. S10. Shaded regions represent the corresponding distribution obtained using 2500 stochastic mRNA trajectories and taking the last cell-cyle to approximate the distribution.

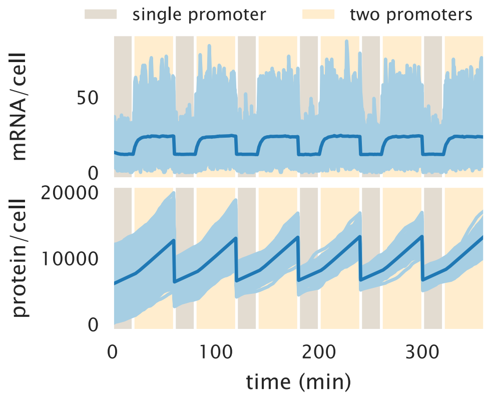

**Figure S22. Stochastic trajectories of mRNA and protein counts.** 2500 protein counts over time for a two-state unregulated promoter. Cells spend a fraction of the cell cycle with a single copy of the promoter (light brown) and the rest of the cell cycle with two copies (light yellow). When trajectories reach a new cell cycle, the molecule counts undergo a binomial partitioning to simulate the cell division.

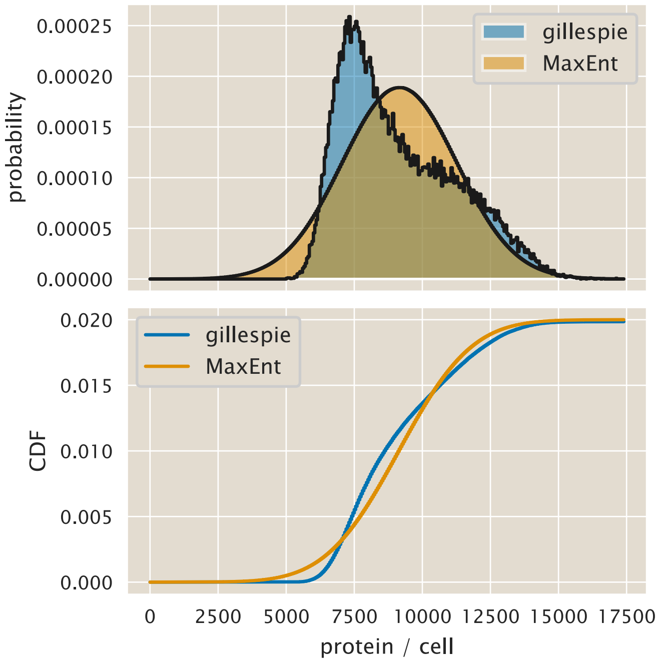

**Figure S23. Comparison of protein distributions.** Comparison of the protein distribution generated with Gillespie stochastic simulations (blue curve) and the maximum entropy approach presented in Appendix S5 (orange curve). The upper panel shows the probability mass function. The lower panel compares the cumulative distribution functions.

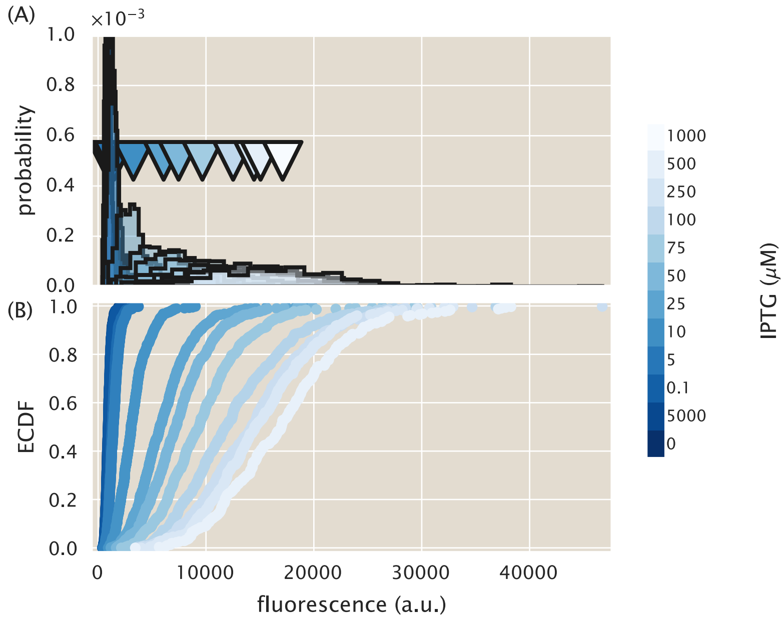

**Figure S24. Single cell fluorescence distributions for different inducer concentrations.** Fluorescence distribution histogram (A) and cumulative distribution function (B) for a strain with 260 repressors per cell and a binding site with binding energy $\Delta\varepsilon_r = -13.9\ k_BT$. The different curves show the single cell fluorescence distributions under the 12 different IPTG concentrations used throughout this work. The triangles in (A) show the mean of each of the distributions.

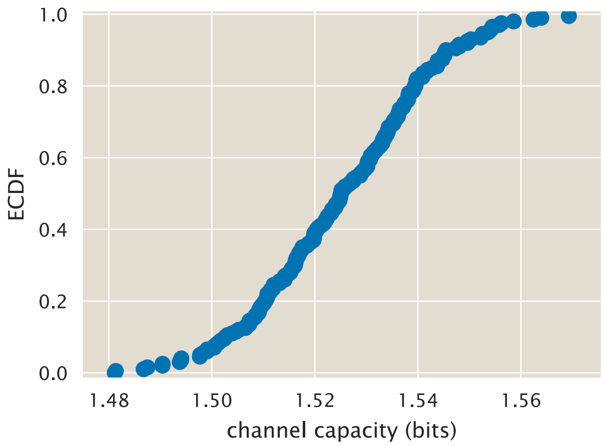

**Figure S25. Channel capacity bootstrap for experimental data.** Cumulative distribution function of the resulting channel capacity estimates obtained by subsampling 200 times 50% of each distribution shown in Fig. S24, binning it into 100 bins, and feeding the resulting **Q** matrix to the Blahut-Arimoto algorithm.

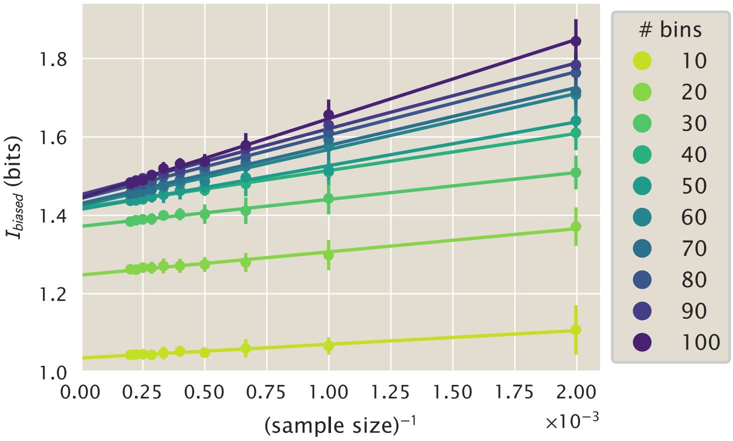

**Figure S26. Inverse sample size vs channel capacity.** As indicated in Eq. S152 if the channel capacity obtained for different subsample sizes of the data is plotted against the inverse sample size there must exist a linear relationship between these variables. Here we perform 15 bootstrap samples of the data from Fig. S24, bin these samples using different number of bins, and perform a linear regression (solid lines) between the bootstrap channel capacity estimates, and the inverse sample size.

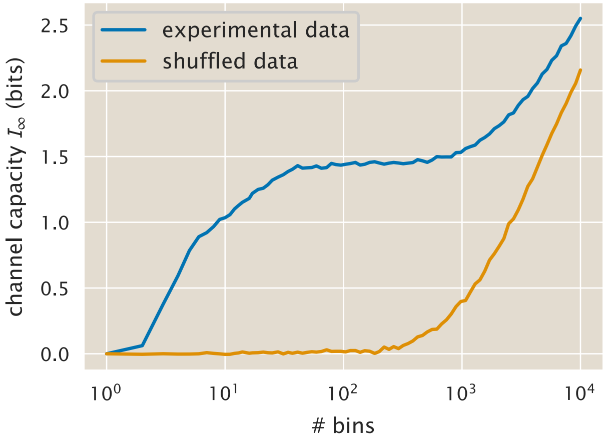

**Figure S27. Channel capacity as a function of the number of bins.** Unbiased channel capacity estimates obtained from linear regressions as in Fig. S26. The blue curve show the estimates obtained from the data shown in Fig. S24. The orange curve is generated from estimates where the same data is shuffled, loosing the relationship between fluorescence distributions and inducer concentration.

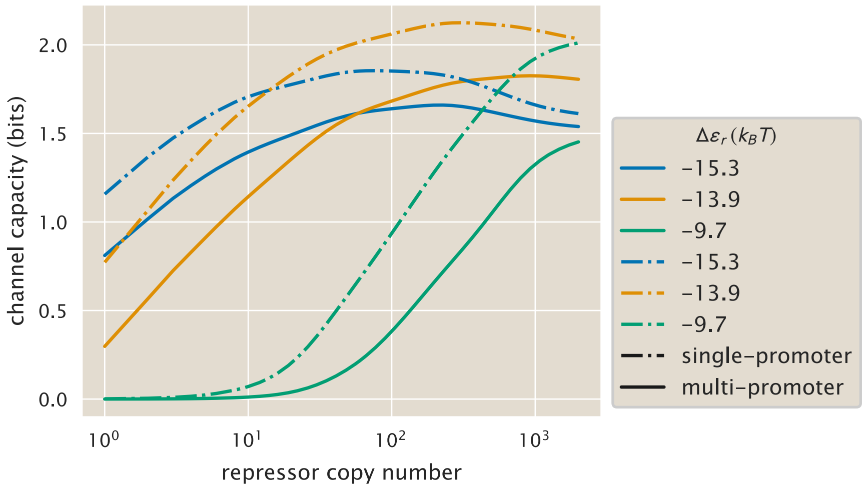

**Figure S28. Comparison of channel capacity predictions for single- and multi-promoter models.** Channel capacity for the multi-promoter model (solid lines) vs. the single-promoter steady state model (dot-dashed lines) as a function of repressor copy numbers for different repressor-DNA binding energies. The single-promoter model assumes Poissonian protein degradation ($\gamma_p > 0$) and steady state, while the multi-promoter model accounts for gene copy number variability and during the cell cycle and has protein degradation as an effect due to dilution as cells grow and divide.

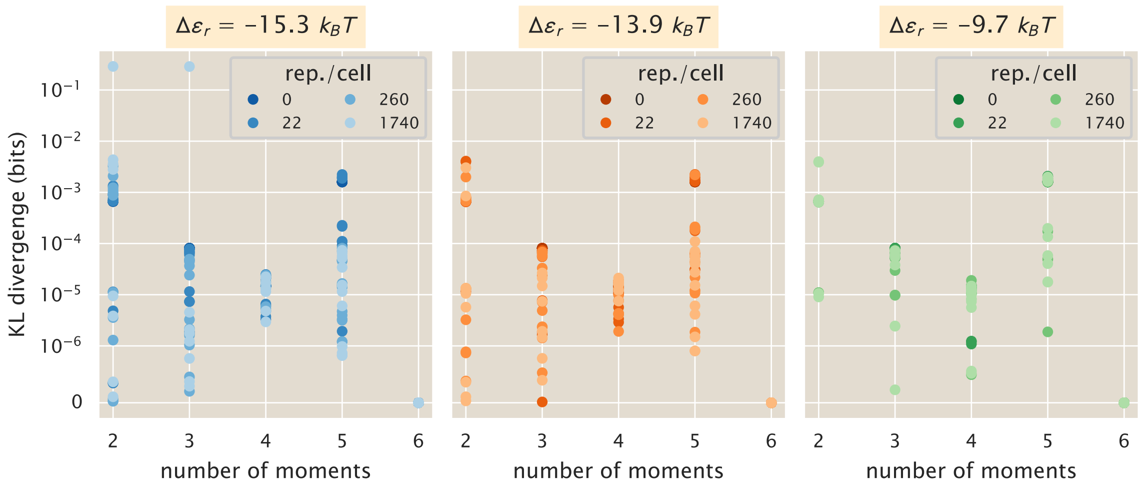

**Figure S29. Measuring the loss of information by using different number of constraints.** The Kullback-Leibler divergence was computed between the maximum entropy distribution constructed using the first 6 moments of the distribution and a variable number of moments.

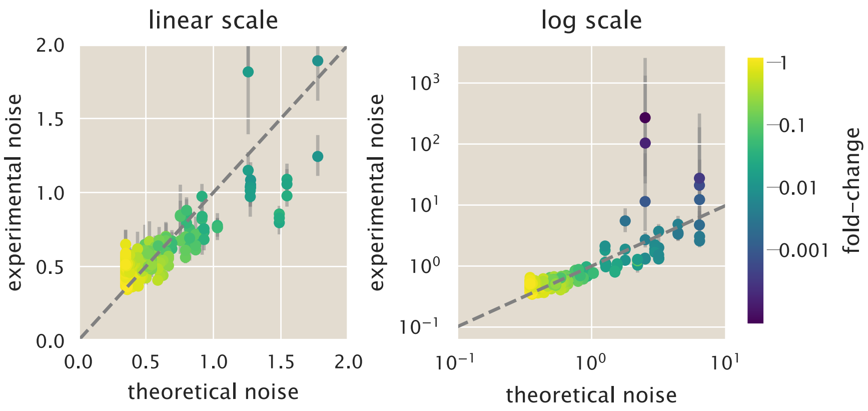

**Figure S30. Multiplicative factor to improve theoretical vs. experimental comparison of noise in gene expression.** Theoretical vs. experimental noise both in linear (left) and log (right) scale. The dashed line shows the identity line of slope 1 and intercept zero. All data are colored by the corresponding value of the experimental fold-change in gene expression as indicated by the color bar. The $x$-axis was multiplied by a factor of $\approx 1.5$ as determined by a linear regression from the data in Fig. S11. Each datum represents a single date measurement of the corresponding strain and IPTG concentration with $\geq 300$ cells. The points correspond to the median, and the error bars correspond to the 95% confidence interval as determined by 10,000 bootstrap samples.

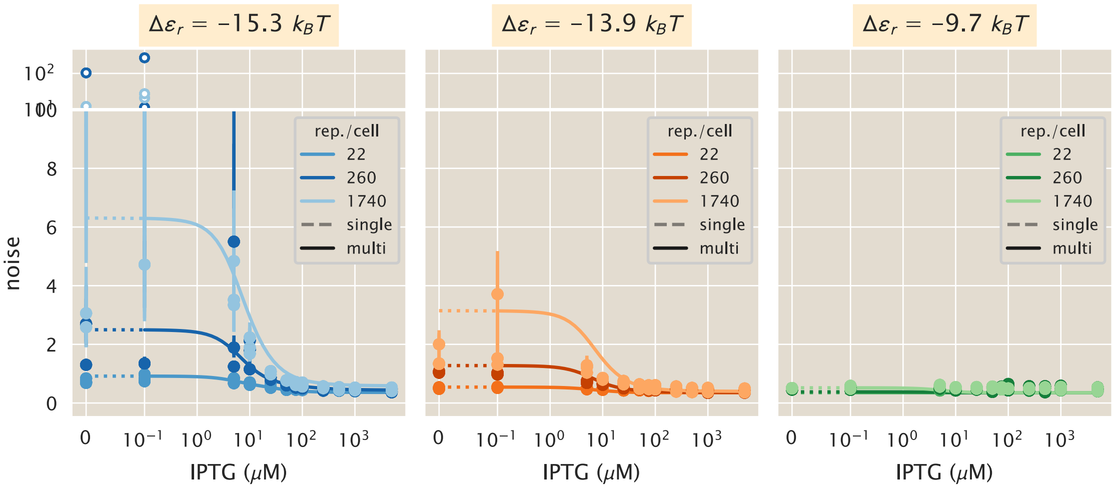

**Figure S31. Protein noise of the regulated promoter with multiplicative factor.** Comparison of the experimental noise for different operators ((A) O1, $\Delta\varepsilon_r = -15.3\ k_BT$, (B) O2, $\Delta\varepsilon_r = -13.9\ k_BT$, (C) O3, $\Delta\varepsilon_r = -9.7\ k_BT$) with the theoretical predictions for the the multi-promoter model. A linear regression revealed that multiplying the theoretical noise prediction by a factor of $\approx 1.5$ would improve agreement between theory and data. Points represent the experimental noise as computed from single-cell fluorescence measurements of different *E. coli* strains under 12 different inducer concentrations. Dotted line indicates plot in linear rather than logarithmic scale. Each datum represents a single date measurement of the corresponding strain and IPTG concentration with $\geq 300$ cells. The points correspond to the median, and the error bars correspond to the 95% confidence interval as determined by 10,000 bootstrap samples. White-filled dots are plot at a different scale for better visualization.

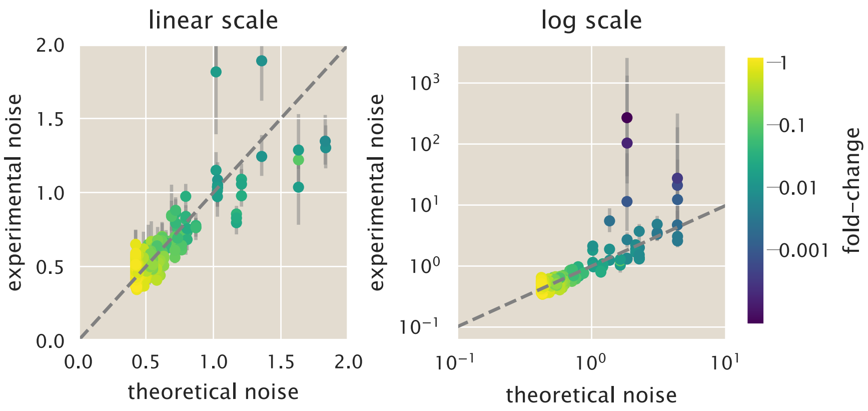

**Figure S32. Additive factor to improve theoretical vs. experimental comparison of noise in gene expression.** Theoretical vs. experimental noise both in linear (left) and log (right) scale. The dashed line shows the identity line of slope 1 and intercept zero. All data are colored by the corresponding value of the experimental fold-change in gene expression as indicated by the color bar. A value of $\approx 0.2$ was added to all values in the $x$-axis as determined by a linear regression from the data in Fig. S11. Each datum represents a single date measurement of the corresponding strain and IPTG concentration with $\geq 300$ cells. The points correspond to the median, and the error bars correspond to the 95% confidence interval as determined by 10,000 bootstrap samples.

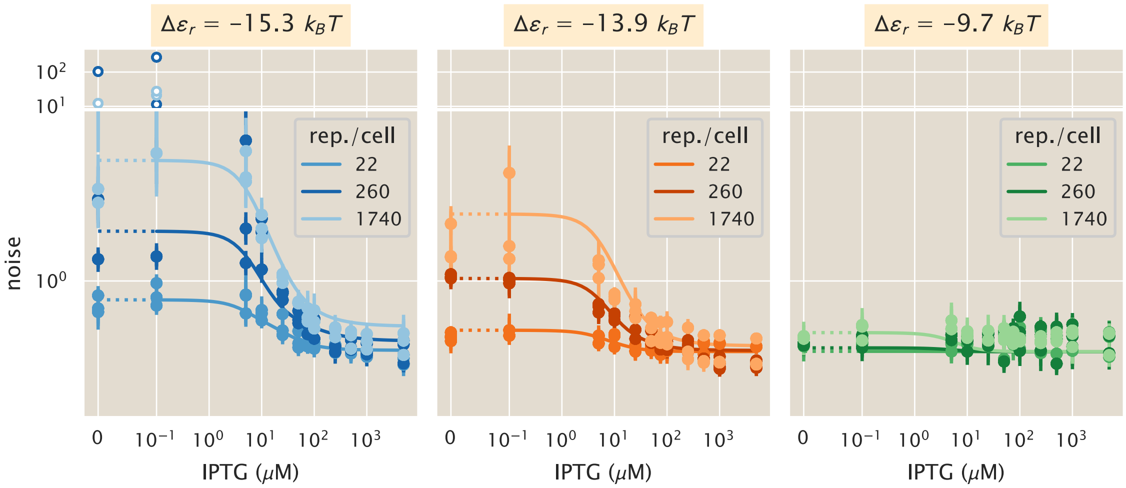

**Figure S33. Protein noise of the regulated promoter with additive factor.** Comparison of the experimental noise for different operators ((A) O1, $\Delta\varepsilon_r = -15.3\ k_BT$, (B) O2, $\Delta\varepsilon_r = -13.9\ k_BT$, (C) O3, $\Delta\varepsilon_r = -9.7\ k_BT$) with the theoretical predictions for the the multi-promoter model. A linear regression revealed that an additive factor of $\approx 0.2$ to the the theoretical noise prediction would improve agreement between theory and data. Points represent the experimental noise as computed from single-cell fluorescence measurements of different *E. coli* strains under 12 different inducer concentrations. Dotted line indicates plot in linear rather than logarithmic scale. Each datum represents a single date measurement of the corresponding strain and IPTG concentration with $\geq 300$ cells. The points correspond to the median, and the error bars correspond to the 95% confidence interval as determined by 10,000 bootstrap samples. White-filled dots are plot at a different scale for better visualization.

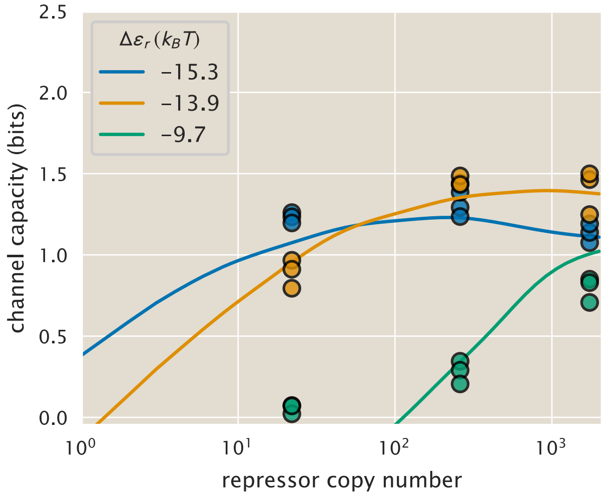

**Figure S34. Additive correction factor for channel capacity.** Solid lines represent the theoretical predictions of the channel capacity shown in Fig. 5(A). The dashed lines show the resulting predictions with a constant shift of -0.43 bits. Points represent single biological replicas of the inferred channel capacity.
